# Future Range Shifts and Diversity Patterns of Antarctic Lecideoid Lichens Under Climate Change Scenarios

**DOI:** 10.1101/2025.06.23.661079

**Authors:** Anna Götz, Mikhail Andreev, Robert R. Junker, Lea Maislinger, Leopoldo G. Sancho, Wolfgang Trutschnig, Ulrike Ruprecht

## Abstract

Being uniquely adapted to extreme environmental conditions, rock-dwelling lecideoid lichens are a diverse and major component of terrestrial vegetation in Antarctica. Climate change is reshaping Antarctic ecosystems, forcing cold-adapted species to migrate to maintain their climatic niche. The study surveyed the circum Antarctic lecideoid lichen diversity and modelled the impacts of two climate change scenarios on their distributional range shifts across Antarctica.

Myco- and photobiont symbionts of lecideoid lichen species from a circum-Antarctic sampling were classified using classical barcoding methods. The climatic niches of nine common mycobiont species and four photobiont OTUs were modelled, and range shifts were projected across four Antarctic bioregions under two Shared Socioeconomic Pathways: (a) SSP1-2.6: sustainable development (2.6 Wm^-2^, +3°C) and (b) SSP5-8.5: continued dependence on fossil fuels (8.5 Wm^-2^, +5.1°C).

DNA barcoding revealed altogether 34 species of lecideoid lichens and 9 photobiont OTUs for the Antarctic continent. In addition to the already known lichen species in Antarctica, three newly detected species of the genus *Lecidella* could be identified. The calculated climate change scenarios across bioregions predict greater range expansion for mycobiont species and photobiont OTUs than climate-induced habitat loss. While niche contraction is expected in maritime Antarctica, species distributions are expected to expand in continental regions, primarily due to inland shifts. These inland areas may serve as emerging climatic refugia for certain mycobiont species.

Overall, these results suggest that, under future warming, lecideoid lichens undergo an overall range expansion, particularly in previously uncolonized inland areas in continental Antarctica.

## Introduction

The biogeographical and bioclimatic characteristics of the Antarctic continent only allow life on a very reduced scale. With an area of 14 Mio km^2^ Antarctica is the 5^th^ largest continent on Earth, but only about 0.81% of the whole continent are ice-free (Burton-Johnson et al., 2016) leaving a highly fragmented habitat as potential area for terrestrial life (Lee et al., 2022a; Peat, Clarke, & Convey, 2007). Those terrestrial ecosystems are mainly limited to non-vascular vegetation such as lichens, fungi, mosses and algae across the continent. Just two vascular plants (*Deschampsia antarctica* Desv. and *Colobanthus quitensis* (Kunth) Bartl.) are restricted to the milder, ocean-influenced areas of maritime Antarctica (Lee et al., 2022a; Peat et al., 2007). In the more climatically extreme areas of continental Antarctica, rock-dwelling (saxicolous) lichens are one of the most successful vegetation-forming pioneer organisms(Castello, 2003; Hertel, 2007; Wagner et al., 2021).

Given the extreme environmental conditions and the limited extent of ice-free areas, the structure and distribution of Antarctic terrestrial ecosystems are particularly sensitive to climatic changes. The effects of climate change vary across different regions of the continent and therefore affecting terrestrial species depending on their spatial distribution. In maritime Antarctica and in particular on the Western Antarctic Peninsula, climate change is occurring faster than in continental Antarctica and exceeds the global average rate of change (Siegert et al., 2019; Turner et al., 2009), contributing to increased glacial melt, more precipitation and the expansion of available habitats (Lee et al., 2017; Lenaerts et al., 2016; Vignon et al., 2021). Recent studies observed an expansion of native vascular plants and invasive plant species into bryophyte and lichen dominated areas in the past decades, but found no evidence for future increase of terrestrial vegetation coverage, so far (Aleksandrov, Andreev, & Kurbatova, 2012; Andreev et al., 2015; Cannone et al., 2022; Chown et al., 2012; Hughes et al., 2020; Parnikoza et al., 2013; Rocha et al., 2024; Torres Mellado, Jaňa, & Casanova-Katny, 2011). Long-term observations in maritime Antarctica indicate that an increase in precipitation seems to affect lichen distribution and coverage negatively due to the enhanced snow cover in maritime Antarctica (Miranda et al., 2020; Sancho et al., 2017) and less available sunlight due to increased cloud cover (Sancho, Pintado, & Green, 2019). In contrast, in the continental part of Antarctica lichen cover has increased with rising temperature and dryness (Brabyn et al., 2005; Johansson & Thor, 2008; Melick & Seppelt, 1997; Olech & Słaby, 2016; Sancho et al., 2017). Particularly the Transantarctic Mountains which currently contain the largest ice-free areas, is expected to undergo relatively limited warming and associated snowmelt (Lee et al., 2017; Turner et al., 2014). Nevertheless, enhanced snowfall is predicted for East Antarctica and increased melt occurs predominantly in West Antarctica (Lenaerts et al., 2016). Biodiversity, however, is expected to remain relatively stable, also allowing for the persistence of endemic species and even promoting the range expansion of native and less motile species (Lee et al., 2017; Lee et al., 2022a).

Among the organisms expected to benefit from these relatively stable conditions are lichens, which form complex mini-ecosystems (Schneider et al., 2011) consisting of a dominant fungus (mycobiont) and a dominant photosynthetic partner (green algae and/or cyanobacteria; photobiont, classified in Operational Taxonomic Units (OTUs)). Additionally, they harbour highly diverse bacterial (Aschenbrenner et al., 2016), endolichenic or lichenicolous fungal communities, basidiomycete yeasts (Arnold et al., 2009; Lawrey & Diederich, 2003; Spribille et al., 2010; Spribille et al., 2016) and green algae (Casano et al., 2011; Guzow-Krzemińska, 2006; Molins et al., 2021; Moya et al., 2017; Peksa & Škaloud, 2011; Ruprecht, Brunauer, & Turk, 2014; Werth & Sork, 2010).

One of the few species groups distributed throughout Antarctica is the heterogenous and rock-dwelling group of lecideoid lichens (Ruprecht et al., 2020; Fryday et al., 2024). The distribution of these lichens ranges from the northernmost, very mild habitats such as Livingston Island (62°S; Hertel 2007, Ruprecht *et al*., 2024), across the Antarctic Peninsula to the extremely cold and dry areas in higher latitudes such as the McMurdo Dry Valleys (78°S; Wagner *et al*., 2020; Pérez-Ortega *et al*., 2023) or the Darwin Area (80S°; Wagner et al., 2021). In particular, the western side facing the Ross Sea, and the Ross Ice Shelf of the Transantarctic Mountains currently contain the largest amount of ice-free area on a bioregional scale (Lee et al., 2017), where a high species diversity of saxicolous lecideoid lichens has been found over the last years (Pérez-Ortega et al., 2023; Wagner et al., 2020; Wagner et al., 2021). Therefore, they serve as a very useful model studying these eco systems with a single species group (Ellis, 2019; Sancho et al., 2019; Wagner et al., 2021).

Previous studies in the McMurdo Dry Valleys (MDV; S78°) showed that the mycobionts *Lecidea cancriformis* C.W. Dodge & G.E. Baker and *Rhizoplaca macleanii* (Dodge) Castello were significantly more abundant at the more humid, colder and higher elevations along the mountain ridges than *Carbonea vorticosa* (Hertel) Hertel and *Lecidea polypycnidophora* U. Rupr. & Türk at the drier valley floor (Wagner et al., 2020). Furthermore, one of the most common photobionts, the Antarctica endemic OTU *Tr*_S18 (Ruprecht et al., 2020; Wagner et al., 2021) is strongly specialized for the climatically most extreme areas in the continent such as the Darwin Area (S80°; Wagner 2021). As the saxicolous lichens depend mainly on atmospheric humidity (clouds, fog, dew) for their growth (Wagner et al., 2020), the species composition also varies based on the local moisture content. Given that the increased humidity caused by climate change does not lead to more snowfall and the covering of potential growing sites, it can therefore be assumed that it could potentially promote the spread but also significantly change the species composition (Lenaerts et al., 2016; Ruprecht et al., 2012b; Sancho et al., 2017; Wagner et al., 2020; Wagner et al., 2021).

Reliable predictions of changes in lichen diversity and distribution in Antarctica can only be made on a robust taxonomic basis. The lecideoid lichen diversity has been studied for over 50 years, and many of these difficult-to-classify species have been described in the early stages (e.g.; Dodge, 1973; Filson, 1974; Knoph & Leuckert, 1994; Inoue, 1995). Nevertheless, advances in molecular techniques have since significantly refined these classifications, leading to synonymization, reclassification, and the description of new species (Castello, 2003; Hertel, 2007; Ruprecht et al., 2010; Ruprecht et al., 2012b; Ruprecht et al., 2020; Ruprecht et al., 2024).

Building on these findings and based on an extensive and circum-Antarctic dataset the aim of this study is to (1) investigate the diversity and distribution of the most common lecideoid lichen species of the genera *Carbonea* (Hertel) Hertel, *Lecanora* Ach., *Lecidea* Ach., *Lecidella* Körb., and *Rhizoplaca* Zopf and their photobionts of the genus *Trebouxia* Puymaly; (2) to assess the current climatic niches of each mycobiont and photobiont throughout Antarctica and (3) to predict future range dynamics for both symbionts for 2100 under two priority Shared Socioeconomic Pathways (SSP), a narrative with different emission trajectories based on the Representative Concentration Pathways (RCPs scenarios of the IPCC; Meinshausen et al., 2020; O’Neill et al., 2016), including (a) the scenario SSP1-2.6 of sustainable development with an approximate climate forcing of 2.6 Wm^-2^ and temperature increase of 3°C from pre-industrial times to 2100 and (b) the scenario SSP5-8.5 with continued dependence on fossil fuels without significant climate protection efforts, with an approximate climate forcing of 8.5 Wm^-2^ and a 5.1°C temperature increase from pre-industrial times to 2100 (Meinshausen et al., 2020; O’Neill et al., 2016; Riahi et al., 2017).

## Material and Methods

### Investigated lichen specimens

This study includes 637 lecideoid lichen specimens across the whole Antarctic continent ranging from a latitude 62°S to 86°S (Table S1). For all subsequent analyses the dataset was filtered for mycobiont species with an overall occurrence of n >= 10 such that the combination of mycobiont species and associated photobiont OTUs >=5. This resulted in 205 specimens including 9 mycobiont species (*Carbonea vorticosa, Lecanora fuscobrunnea* C.W. Dodge & G.E. Baker*, Lecidea andersonii* Filson*, Lecidea atrobrunnea* (Ramond ex Lam. DC.) Schaer.*, Lecidea cancriformis, Lecidea polypycnidophora, Lecidella greenii* U. Ruprecht & Türk, *Lecidella siplei* (C.W. Dodge & G.E. Baker) May. Inoue, and *Rhizoplaca macleanii* and 4 photobiont OTUs of the genus *Trebouxia* (*Tr*_A02, *Tr*_S02, *Tr*_I01 and *Tr*_S18), which constitutes one of the most frequent genera of green algae in lichen symbiosis (Rambold, Friedl, & Beck, 1998). All photobiont OTUs were named according to the equivalence table by Medeiros et al. (2024) in order to guarantee a comprehensible classification of the OTU numbers. Due to the lack of further assignment, *Tr*_S18 is the only photobiont OTU named according to Ruprecht et al. (2020).

Sequences of the barcode-marker ITS were obtained of both symbionts for almost all specimens. Two third of the specimen collections have already been published in Ruprecht, Brunauer, and Printzen (2012a); Ruprecht et al. (2010); Ruprecht et al. (2012b); Wagner et al. (2020); Wagner et al. (2021). Further samples were collected by Mikhail Andreev (Komarov Botanical Institute, St. Petersburg, Russia, 1989 - 2015) at several stations around Continental Antarctica and by Ulrike Ruprecht (University Salzburg, Austria, 2018) at Livingston Island. Additional herbarium material was added from the British Antarctic Survey (BAS) and the Australian Antarctic Division (AAS). The entire list of samples including voucher IDs, species names, OTU numbers and accession numbers can be found in Table S1.

As it is standard practice in Antarctic field surveys, the sampling strategy depended strongly on the accessibility of the areas and the respective research campaigns. All collections were conducted with a representative sample of available lichen specimens collected with a fine chisel on siliceous rock. All voucher specimens are stored in the herbarium of the University of Salzburg (SZU) except for samples collected by Mikhail Andreev which are deposited at the Komarov Botanical Institute, Russian Academy of Sciences, Saint Petersburg, Russia (LE). The study complied with the institutional and national guidelines of the countries, which funded the research projects at the relevant research stations in Antarctica. The PIs of the projects hold permits to collect the lichens in the respective research areas.

### Study Area

To model the climatic niches of lecideoid lichens in Antarctica, we selected four of the continent’s largest ice-free areas. These are defined according the Antarctic Conservation Biogeographic Regions (or bioregions; Terauds & Lee, 2016) and include: (1) Maritime Antarctica (comprising the bioregions Alexander Island, Eastern Peninsula, North East Peninsula, Northern Peninsula, South Orkney Islands and South Shetland Islands), (2) Pensacola Mountains, (3) Prince Charles Mountains and (4) Transantarctic Mountains (including Southern Transantarctic Mountains, Victoria Land North Cape Hallett and Victoria Land South Dry Valleys; Fig. 1a).

**Figure 1:**
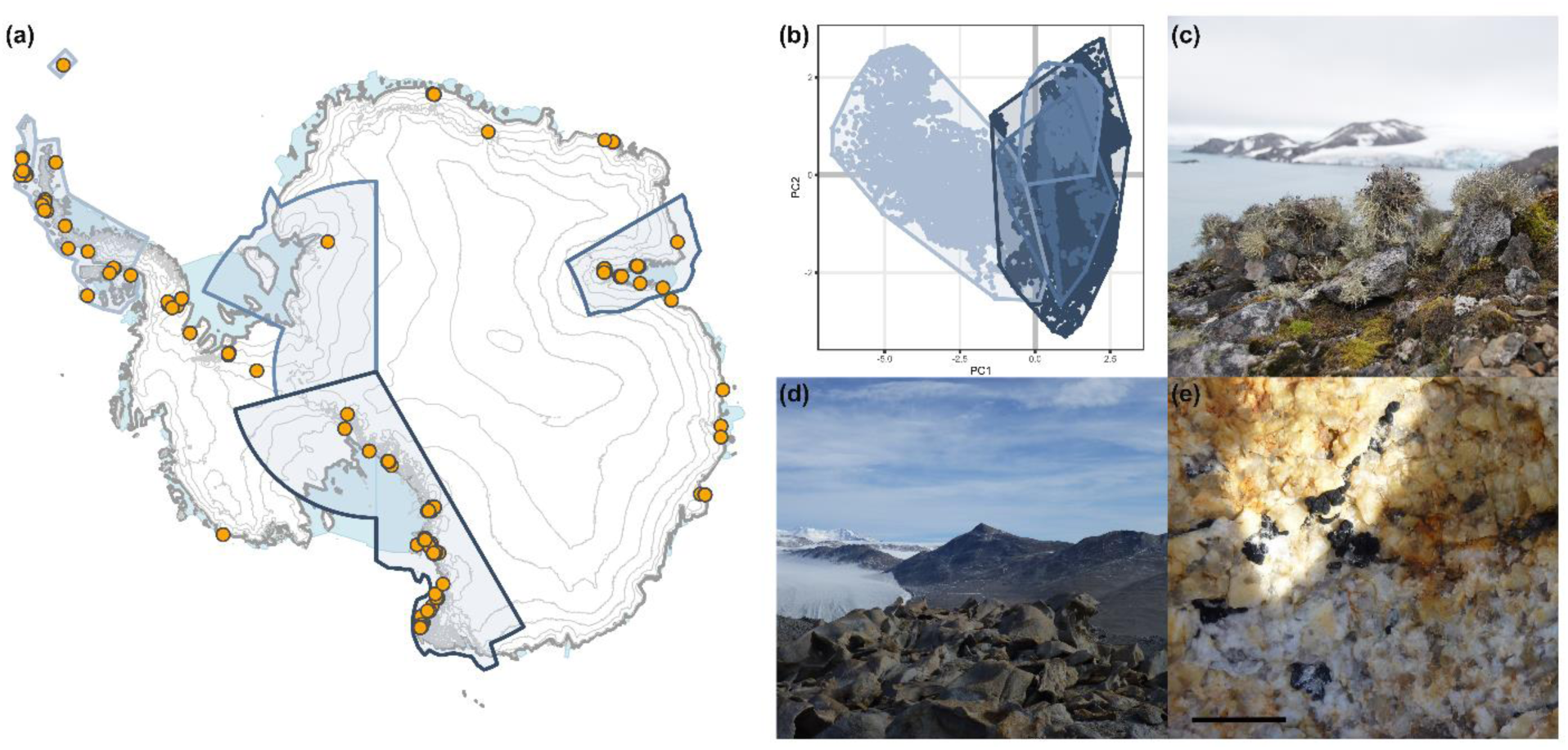
(a) Bioregions and sampling sites selected for this study. Yellow dots represent locations of the collected samples. (b) PCA of the 7 bioclimatic variables (bio1, bio2, bio7, bio8, bio12, bio15, bio18) of the Antarctic rock outcrop, clustered and coloured b y bioregion to show the climatical differentiation between the selected bioregions. (c) Lichen and moss community at Base Juan Carlos I, Livingston Island, maritime Antarctica (S62°). (d) Miers Valley, McMurdo Dry Valleys, continental Antarctica (S78°). (e) Lecideoid liche n, southern ridge Miers valley. Scale bar = 1 cm.

Maritime Antarctica differs most distinctly from the continental sites in terms of both current climate and predicted future climatic changes (Fig. 1b). This region shows relatively mild temperatures and the highest level of precipitation, much of which falls as rain during the austral summer and occasionally during the colder months (Turner et al., 2020; Vignon et al., 2021; Wagner et al., 2018). In contrast, the Pensacola Mountains, Prince Charles Mountains, and Transantarctic Mountains, part of continental Antarctica, generally experience lower mean annual temperatures and less to no precipitation. Most moisture arrives as snow, with little to no rainfall (Scarchilli et al., 2020).

Among the continental regions, the Prince Charles Mountains exhibit slightly higher annual mean temperatures compared to the Pensacola Mountains and the Transantarctic Mountains. The climate in the Transantarctic Mountains is largely driven by extremely dry and strong katabatic winds (Turner et al., 2019; Vignon et al., 2021). Particularly on the southern slopes of the Transantarctic Mountains, these winds generate a pronounced rain-shadow effect on the leeward side, resulting in drier and colder conditions near the ice sheet interior (Lenaerts et al., 2016; Nicolas & Bromwich, 2014; Vignon et al., 2021). With temperatures being constantly subzero, the majority of precipitation only occurs as snow (Scarchilli et al., 2020) which often sublimates directly and is therefore in most cases not available as a water source for terrestrial vegetation growing on rocks above the melting streams (Wagner et al., 2020).

In sheltered microhabitats, the water retention capacity of the substrate, melting snow, as well as the available humidity of dew, fog and clouds are another key factors for possible lichen growth in the extreme habitats of the continent, in addition to wind and sun protection (Wagner et al., 2021; Yung et al., 2014; Zucconi et al., 2016). Microclimatic conditions are also dependent on local and further on macroclimatic conditions, vary considerably within these regions and can strongly influence local lichen diversity and abundance. For example, in the Prince Charles Mountains near Lake Radok (Fig. S1), the valley is sheltered from prevailing southerly winds, creating a relatively temperate microclimate. The presence of a polynya on the lake surface increases local humidity, supporting a richer vegetation cover. Similarly, in the Clemens Massif (Fig. S1), meltwater from the adjacent Lambert Glacier sustains rivers and lakes, generating a highly humid environment that supports extensive moss and lichen communities, including large moss cushions. In contrast, the Glybovyye Mountains (Rymill, Stinear, and Bloomfield ranges; Fig. S1,) experience significantly harsher and drier conditions, with near-permanent subzero temperatures and sparse or absent vegetation (Andreev, 2023).

Although microhabitats play a key role in supporting lichen communities, their characteristics are shaped by local and broader macroclimatic and geographic conditions which are also represented in the climate models. As a result, large-scale climate gradients still largely determine the availability and distribution of suitable microclimatic refugia (Wagner et al., 2021)

### DNA amplification, sequencing and species delimitation

Genomic DNA was extracted from individual thalli by using DNeasy Plant Mini Kit (Qiagen, Hilden, Germany) following the manufacturer’s instructions. For all samples, the internal transcribed spacer (ITS) region of the mycobionts’ and photobionts’ nuclear ribosomal DNA (nrITS) were sequenced and amplified. The primers used for amplifying the nrITS region are shown in Table S2.

All reactions were performed with standard methods as described in Ruprecht et al. (2020). Unpurified PCR - products were sent to Eurofins Genomics/Germany for sequencing. The sequences of the mycobionts’ and photobiont’ ITS region were assembled and edited with Geneious version 6.1.8 (https://www.geneious.com) and aligned with MAFFT v7.017 (Katoh et al., 2002) using pre-set settings. Assignment of the specimens (myco-/photobiont) were conducted using a maximum likelihood (ML) approach on the IQ-TREE web server (Trifinopoulos et al., 2016) based on the existing phylogenies of Ruprecht et al. (2020); Wagner et al. (2020) with default settings. In order to ensure the accurate molecular assignment to the corresponding taxa of the examined lichen samples, they were morphologically examined using microscopic and chemical methods (Ruprecht et al., 2024).

### Environmental variables and Climate forcing scenarios

All data analysis was conducted in the R environment version 4.3.3 (RCoreTeam, 2024). Climatic data (bioclim variables) for present and future (2071-2100 time period) conditions were obtained from CHELSA v2.1 database (Karger et al., 2017). Bioclimatic variables were grouped into correlation clusters using raster.cor.matrix() from the package ENMTools (Warren et al., 2021) and visualized as dendrogram shown in Fig. S2. From each cluster, the most relevant variable for the Antarctic region and its species was selected (Fig. S2), which resulted in working with the bioclimatic variables 1 (Annual Mean Temperature), 2 (Mean Diurnal Range), 7 (Temperature Annual Range), 8 (Mean Temperature of Driest Quarter), 12 (Annual Precipitation), 15 (Precipitation Seasonality), 18 (Precipitation of Warmest Quarter) for all subsequent analyses. The variable importance (the aggregated variable importance of each individual model weighted by their contribution) of each bioclim variable to the built the ensemble models was obtained with get_variable_importance() from the package biomod2 and can be found in Fig. S3.

Two different Shared Socio-Economic Pathways (SSPs) with different Climate Forcing Scenarios (RCPs) were used to model future species niches for the period 2071-2100: (1) SSP1-2.6: a scenario of sustainable development with an approximate climate forcing of 2.6 Wm^-2^ and temperature increase of 3°C from pre-industrial times to 2100 and (2) SSP5-8.5: a scenario with continued dependence on fossil fuels without significant climate protection efforts, with an approximate climate forcing of 8.5 Wm^-2^ and a 5.1°C temperature increase from pre-industrial times to 2100 (Meinshausen et al., 2020; O’Neill et al., 2016; Riahi et al., 2017).

For modelling future species distributions for the two respective SSP-RCPs all global climate models (GCMs) of the Coupled Model Intercomparison Project Phase 6 (CMIP6) (O’Neill et al., 2016; Tebaldi et al., 2021) used, which are available at CHELSA database v2.1: GFDL-ESM4 (Krasting et al., 2018), IPSL-CM6A-LR (Boucher et al., 2018), MPI-ESM1-2-HR (Schupfner et al., 2019), MRI-ESM2-0 (Yukimoto et al., 2019) and UKESM1-0-LL (Tang et al., 2019). In order to visualize the overall range dynamics, the mean results of the individual niche models of each GCM were merged into an ensemble mean, min and max.

### Spatial Processing

All climatic data, additional layers and specimen locations were projected and processed in the CRS EPSG:3031 WGS 84 / Antarctic Polar Stereographic. Initial resolution of obtained CHELSA raster layers was 0.008333333 to 0.008333333 degrees in the WGS84 projection (EPSG 4326). Using the function aggregate()from the package terra (Hijmans et al., 2022) the raster layers were downscaled by aggregating 4 pixels horizontally and 5 vertically into their mean. This resulted in a resolution of 2380 m to 3680 m per pixel in the Antarctic Stereographic Projection EPSG 3031.

For modelling species distributions under current and SSP1-2.6 climate conditions, all climate data were cropped to the Antarctic rock outcrop (“Medium resolution vector polygons of Antarctic rock outcrop” from SCAR Antarctic Digital Database (ADD; Gerrish, 2020), representing the available space for all terrestrial organisms on the Antarctic continent. In order to model species distributions under the climate conditions of the SSP5-8.5 scenario, the additional rock outcrop exposed by snowmelt at warmer temperatures has to be taken into account. Therefore, the future rock outcrop (PS_RCP85_BestFuture_IceFree) provided by Lee et al. (2017) on the Australian Antarctic Data Centre (ADDC; https://data.aad.gov.au/) was considered.

### Ensemble models

To obtain current and future species distributions and calculate range shift, the weighted Ensemble Mean (EMwmean) probabilities of occurrence were calculated using the biomod2 package (Thuiller et al., 2016). Parallel to the biomod2 ensemble modelling, all niche calculations and projections were also done with the Maxent modelling algorithm from the package dismo (Hijmans et al., 2017). Based on the comparison of the model performance (AUC/TSS) we selected the biomod2 ensemble models for further downstream analyses.

In the biomod2 modelling process the final models with True Skill Statistic (TSS) > 0.6 and Receiver Operator characteristics (ROC) > 0.9 were selected for ensemble modelling, which resulted in a total of 11 models: ANN (Artificial Neural Network), CTA (Classification Tree Analysis), FDA (Flexible Discriminant Analysis), GAM (Generalized Additive Model), GBM (Boosted Regression Trees), GLM (Generalized Linear Model), MARS (Multiple Adaptive Regression Splines), MAXNET (Maximum Entropy), RF (Random Forest), SRE (Surface Range Envelop), XGBOOST (eXtreme Gradient Boosting Training). The weighted Ensemble Mean (EMwmean, weighted by model TSS value) was used for all downstream analysis to ensure that models with higher accuracy are upweighted.

### Calculation of niche area change

For each climate model of CMIP6 (see Climate data and models) the future ensemble projection was created and range differences to the current distribution were calculated per pixel via the function BIOMOD_RangeSize() by applying the formula *proj.future − 2 * proj.current*. This resulted in four different discrete states for each pixel for each climate forcing scenario: −2 = predicted to be lost; −1 = predicted to remain occupied; 0 = predicted to remain unoccupied and 1 = predicted to be gained. EM area gains (pixel state = 1) and losses (pixel state = −2), as well as overall current (pixel state = −2 and −1) future probable niche area (pixel state = −1 and 1) and the delta probable niche area (future - current niche area) were calculated. Each was calculated per CMIP6 climate model (GFDL-ESM4, IPSL-CM6A-LR, MPI-ESM1-2-HR, MRI-ESM2-0, UKESM1-0-LL) with a pixel size of 8.76km^2^.

For further analysis and visualization, the mean niche gain and loss of all climate models was considered with the respective minimum and maximum of each climate model. Current niche overlap (Warrens’ I; Warren, Glor, & Turelli, 2008) was calculated using the function nicheOverlap() from the package dismo (Hijmans et al., 2017).

For each pixel, distance from coast for each pixel was calculated using the function st_distance() from the package sf (Pebesma, 2018) with the High resolution vector polylines of the Antarctic coastline by SCAR Antarctic Digital Database (ADD).

When predicting future species niches, several constraints must be considered. Species distribution models (SDMs) inherently produce probabilistic estimates rather than definitive/crisp outcomes. These values do not represent the actual distribution but rather a probability of appearance based on the input data, including species records, pseudo-absences, and environmental variables. Area gains and losses, which were calculated as proposed beforehand are therefore also just probabilities Alimited or incomplete species record, which is often the case for Antarctic species, is a well-recognized challenge in SDMs (Fois et al., 2018). We attempted to address this problem by working with ensemble models integrating multiple modelling approaches and hence, enhancing the robustness of predictions. However, small-scale niche predictions, such as those concerning the future distribution of *Tr*_S18 in maritime Antarctica, must be interpreted with caution.

## Results

### Species diversity

In total, the dataset contained 34 mycobiont species (Fig. 2a; Fig. S4a, Table 1) and 9 photobiont OTUs (Fig2b; Fig.4b; Table 2) distributed across maritime and continental Antarctica. Most of the mycobiont species listed in Table 1 have already been described for the entire Antarctic continent in former studies (Castello, 2003; Dodge, 1973; Filson, 1974; Hertel, 2007; Inoue, 1995; Øvstedal & Smith, 2001; Ruprecht et al., 2020; Ruprecht et al., 2010; Ruprecht et al., 2012b; Wagner et al., 2021).

**Figure 2:**
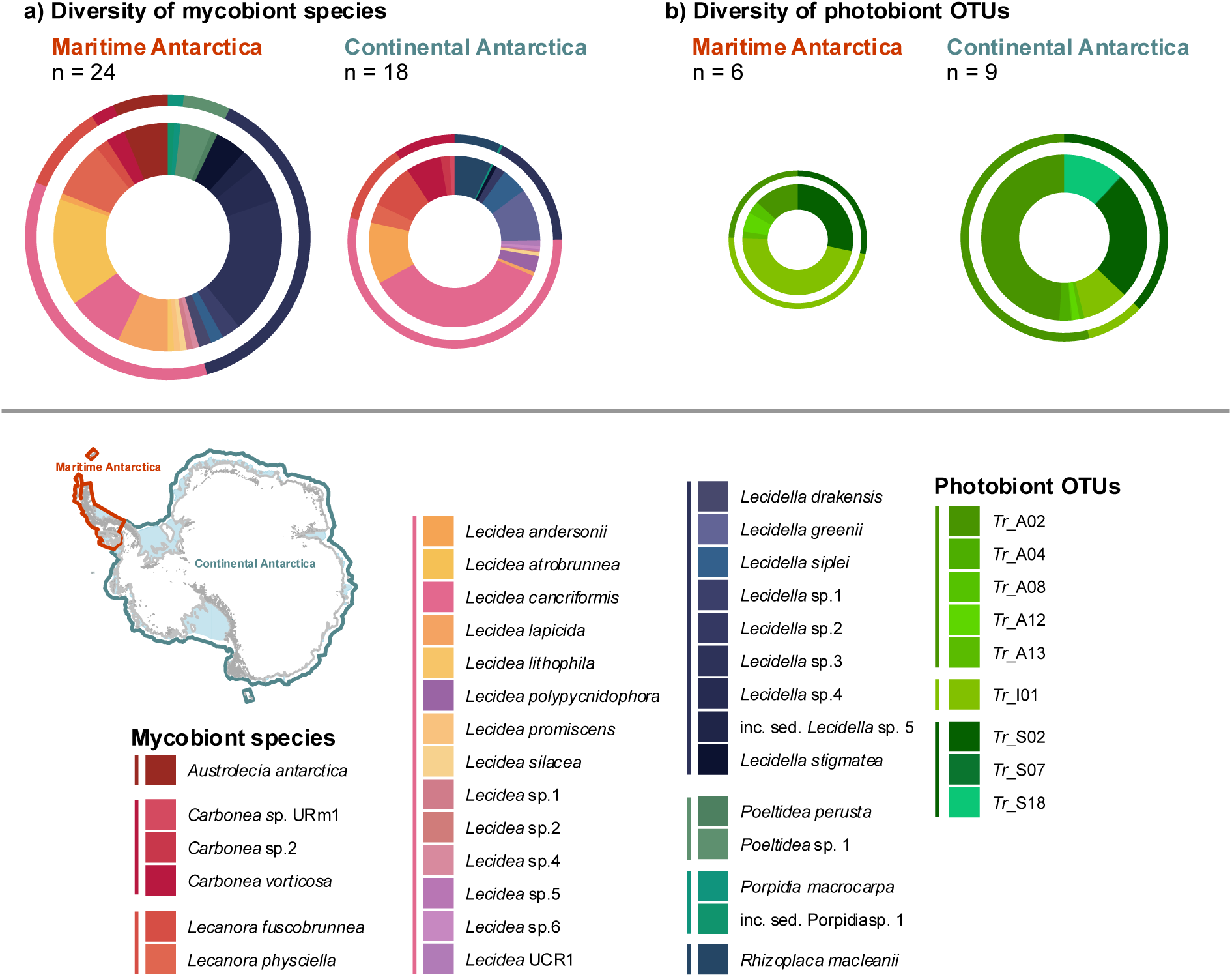
Diversity of mycobiont species and photobiont OTUs in the maritime Antarctica and continental Antarctica. a) Overview of the distinction between maritime and continental Antarctica; b) Composition and number of identified mycobiont species in maritime and continental Antarctica in this study. Inner ring shows the species level, outer ring shows the genus level; c) Composition and number of identified photobiont OTUs in maritime and continental Antarctica in this stud. Inner ring shows the OTU level, outer ring shows the respective Trebouxia clade. Size of the pie charts correspond to the observed species/OTU number.

**Table 1:**
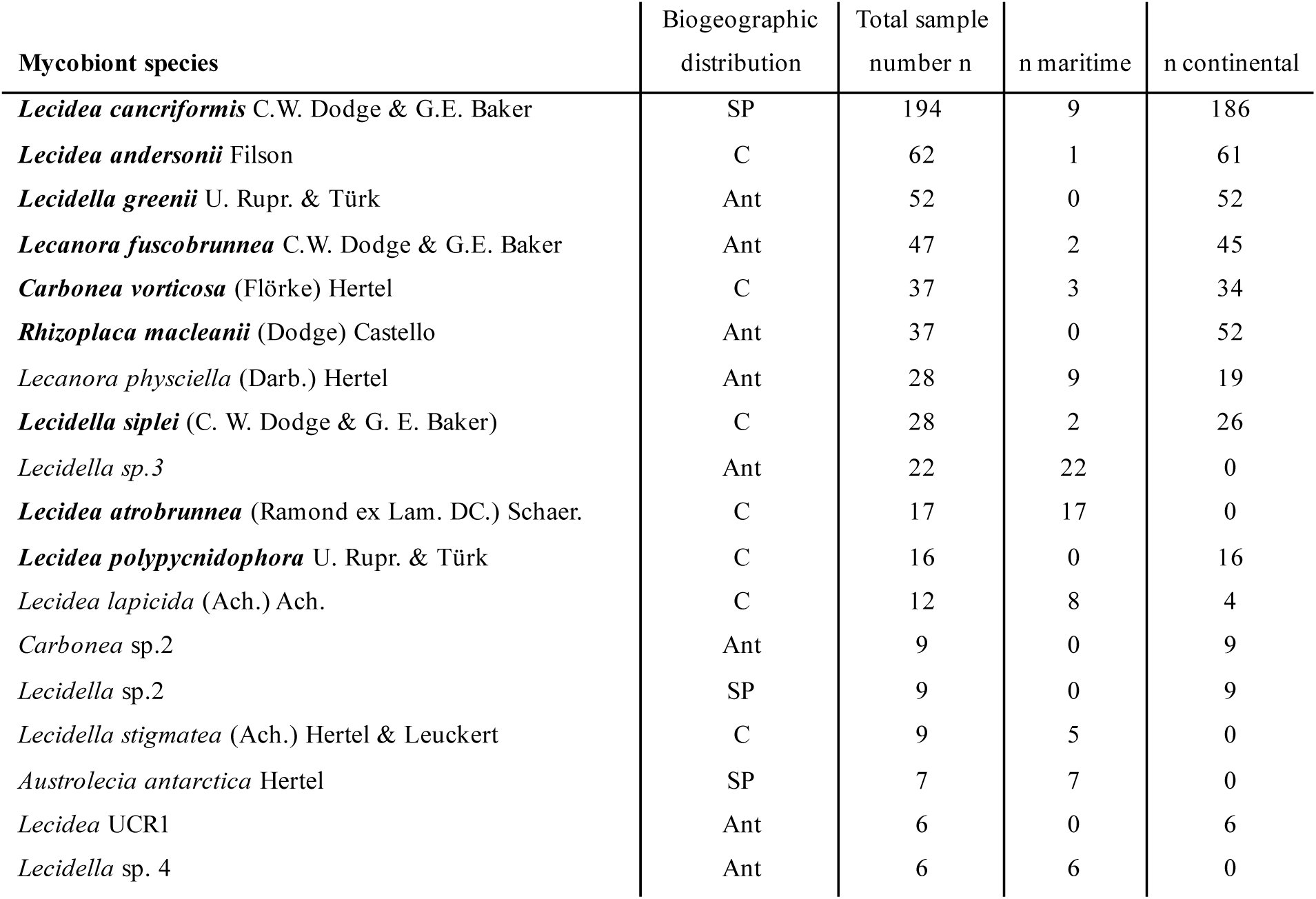
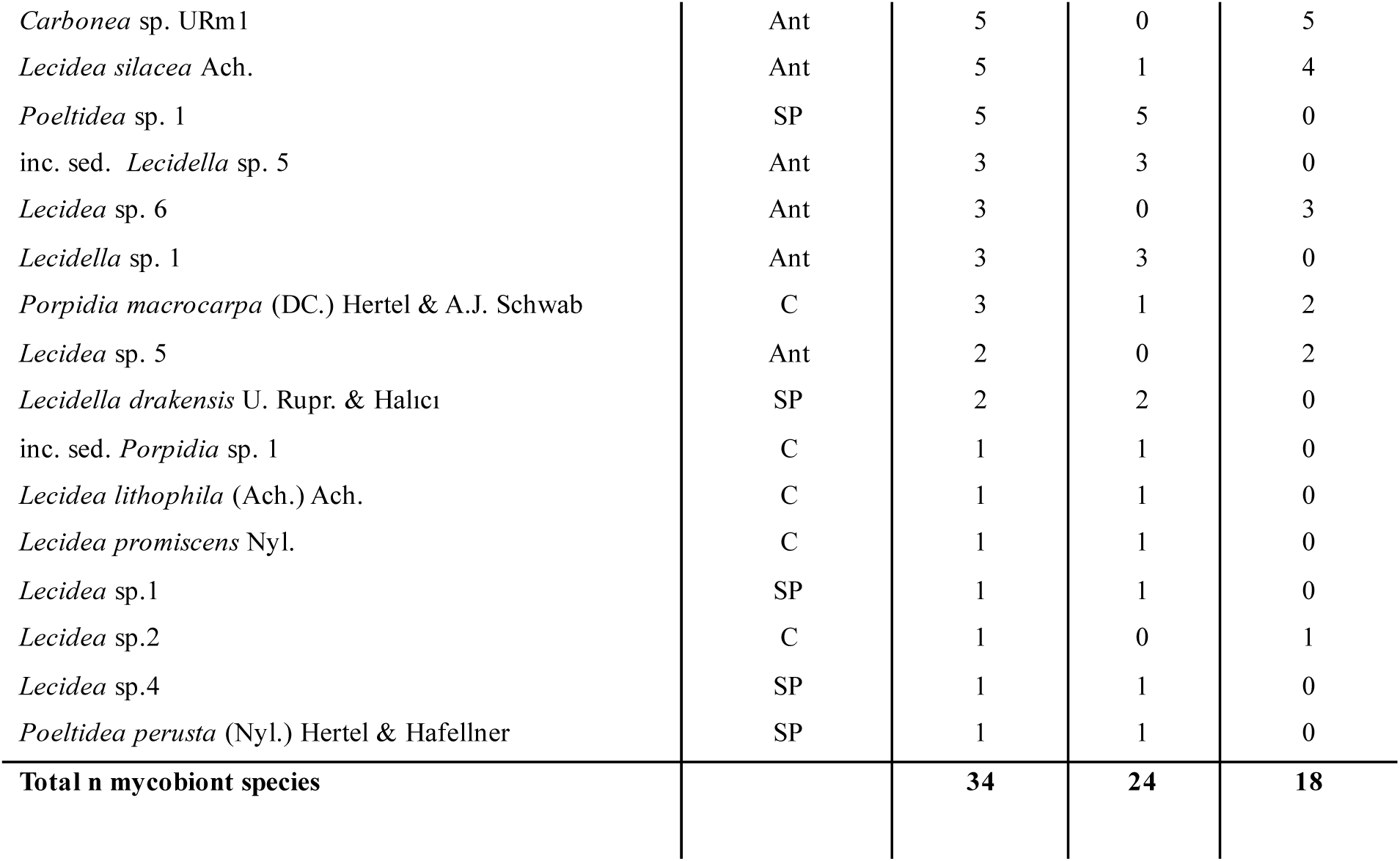
Number n of all main mycobiont species for maritime and continental Antarctica (Fig. 2b). Bold names indicate the mycobiont species which were used in the subsequent niche modelling. Lowest row of the table gathers the total photobiont OTU count per largescale region. Abbreviations for the biogeographic distribution information for each species: C - cosmopolitan; SP - southern Polar; Ant - Antarctica.

| Mycobiont species | Biogeographic distribution | Total sample number *n* | *n* maritime | *n* continental |
| --- | --- | --- | --- | --- |
| ***Lecidea cancriformis*** C.W. Dodge & G.E. Baker | SP | 194 | 9 | 186 |
| ***Lecidea andersonii*** Filson | C | 62 | 1 | 61 |
| ***Lecidella greenii*** U. Rupr. & Türk | Ant | 52 | 0 | 52 |
| ***Lecanora fuscobrunnea*** C.W. Dodge & G.E. Baker | Ant | 47 | 2 | 45 |
| ***Carbonea vorticosa*** (Flörke) Hertel | C | 37 | 3 | 34 |
| ***Rhizoplaca macleanii*** (Dodge) Castello | Ant | 37 | 0 | 52 |
| *Lecanora physciella* (Darb.) Hertel | Ant | 28 | 9 | 19 |
| ***Lecidella siplei*** (C. W. Dodge & G. E. Baker) | C | 28 | 2 | 26 |
| *Lecidella sp.3* | Ant | 22 | 22 | 0 |
| ***Lecidea atrobrunnea*** (Ramond ex Lam. DC.) Schaer. | C | 17 | 17 | 0 |
| ***Lecidea polypycnidophora*** U. Rupr. & Türk | C | 16 | 0 | 16 |
| *Lecidea lapicida* (Ach.) Ach. | C | 12 | 8 | 4 |
| *Carbonea sp.2* | Ant | 9 | 0 | 9 |
| *Lecidella sp.2* | SP | 9 | 0 | 9 |
| *Lecidella stigmatea* (Ach.) Hertel & Leuckert | C | 9 | 5 | 0 |
| *Austrolecia antarctica* Hertel | SP | 7 | 7 | 0 |
| *Lecidea* UCR1 | Ant | 6 | 0 | 6 |
| *Lecidella sp. 4* | Ant | 6 | 6 | 0 |
| *Carbonea* sp. URm1 | Ant | 5 | 0 | 5 |
| *Lecidea silacea* Ach. | Ant | 5 | 1 | 4 |
| *Poeltidea* sp. 1 | SP | 5 | 5 | 0 |
| inc. sed. *Lecidella* sp. 5 | Ant | 3 | 3 | 0 |
| *Lecidea* sp. 6 | Ant | 3 | 0 | 3 |
| *Lecidella* sp. 1 | Ant | 3 | 3 | 0 |
| *Porpidia macrocarpa* (DC.) Hertel & A.J. Schwab | C | 3 | 1 | 2 |
| *Lecidea* sp. 5 | Ant | 2 | 0 | 2 |
| *Lecidella drakensis* U. Rupr. & Halıcı | SP | 2 | 2 | 0 |
| inc. sed. *Porpidia* sp. 1 | C | 1 | 1 | 0 |
| *Lecidea lithophila* (Ach.) Ach. | C | 1 | 1 | 0 |
| *Lecidea promiscens* Nyl. | C | 1 | 1 | 0 |
| *Lecidea* sp.1 | SP | 1 | 1 | 0 |
| *Lecidea* sp.2 | C | 1 | 0 | 1 |
| *Lecidea* sp.4 | SP | 1 | 1 | 0 |
| *Poeltidea perusta* (Nyl.) Hertel & Hafellner | SP | 1 | 1 | 0 |
| **Total n mycobiont species** |  | **34** | **24** | **18** |

**Table 2:** Number n of all photobiont OTUs for maritime and continental Antarctica (Fig. 2c). Bold names indicate the photobiont OTUs which were used in the subsequent niche modelling. Lowest row of the table contains the total photobiont OTU count per largescale region. Abbreviations for the biogeographic distribution information for each OTU: C - cosmopolitan; SP - southern Polar; Ant - Antarctica.

| Photobiont OTU | Biogeographic distribution | Total sample number *n* | *n* maritime | *n* continental |
| --- | --- | --- | --- | --- |
| ***Tr_A02*** | C | 218 | 7 | 211 |
| ***Tr_S02*** | C | 123 | 15 | 108 |
| ***Tr_I01*** | C | 64 | 25 | 39 |
| ***Tr_S18*** | Ant | 51 | 0 | 51 |
| *Tr_A04* | C | 10 | 0 | 10 |
| *Tr_A12* | C | 9 | 3 | 6 |
| *Tr_A13* | C | 5 | 1 | 4 |
| *Tr_A08* | C | 3 | 2 | 1 |
| *Tr_S07* | C | 1 | 0 | 1 |
| **Total n photobiont OTUs** |  | **9** | **6** | **9** |

In addition to two genetically identified but not yet morphologically described species of the genus *Lecidella* (*L*. sp. 1 and *L*. sp. 2; Ruprecht *et al.,* 2020), three further previously unrecognized species (*L.* sp. 3, *L.* sp. 4, inc. sed. *L*. sp. 5) have been identified (Fig. S5) from the Maritime Antarctic, the region with the highest recorded species richness (24), so far for this group. Species diversity for continental Antarctica was lower with 18 species identified. Altogether eight species were recorded in both parts of the continent such as the most common *Lecidea cancriformis* (195 specimens), *L. andersonii* (62)*, Lecanora fuscobrunnea* (47), *Lecidella siplei* (28), *Lecanora physciella* (28), and two rare species such as *Lecidea lapicida* and *Porpidia macrocarpa*. The number of Antarctic endemic (13) and southern polar distributed (8) species covers almost two thirds of all 34 species studied (Ruprecht et al., 2020; Table 1).

Nine OTUs of the genus *Trebouxia* were identified for the photobionts with a clearly higher diversity in continental Antarctica (9) than in maritime Antarctica (6). Four very common OTUs are *Tr*_01 which is dominant in maritime Antarctica and *Tr*_A02 in continental Antarctica followed by the in both parts evenly distributed *Tr*_S02. All OTUs exhibit a cosmopolitan distribution, except for one - *Tr*_S18 - which is endemic to the most climatically extreme areas of the continent (Ruprecht et al., 2020).

### Current species distribution

For modelling the current niches, the most common mycobiont species (n = 9; Table 1) and photobiont OTUs (n = 4; Table 2) were used for all subsequent analyses.

Current mycobiont species niches show a high overlap, except for *Lecidea atrobrunnea*, which is exclusively distributed in maritime Antarctica (Fig. 3a). The rest of the probable niches of mycobiont species form a nested pattern, where climatically restricted species like *Rhizoplaca macleanii* are nested within the probable distributional range of more widely distributed species like *Lecidea cancriformis* and *Lecidea andersonii* (Fig. 3a). The photobiont OTUs also exhibit a similar nested distribution pattern, with *Tr*_I01 predicted to be the most widespread photobiont OTU, likely due to its dominance in the northern Antarctic Peninsula. In contrast, *Tr*_S18 appears to be the most climatically restricted photobiont OTU, with its ecological niche limited to continental Antarctica, mostly within the Transantarctic Mountains. The hotspots of the majority of the mycobiont species and photobiont OTUs are predicted to be in the Transantarctic Mountains, whereas *Lecidea atrobrunnea* and *Tr*_I01 are widely distributed in the maritime Antarctica (Fig. S6).

**Figure 3:**
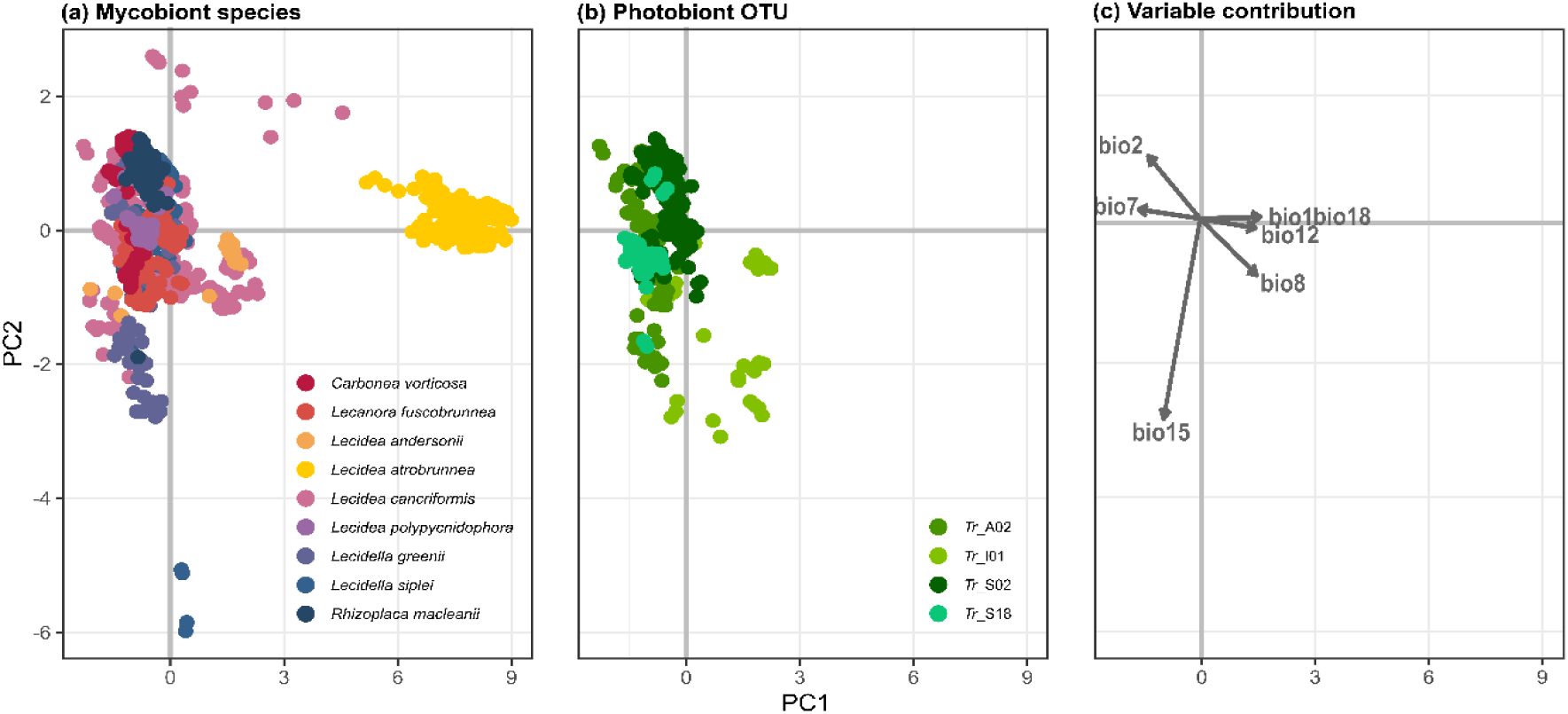
PCA of all pixels with probability of occurrence ≥ 0.75 (current probable niches) of all main (a) mycobiont species and (b) photobiont OTUs in the (bio-)climatic space. (c) Direction of contribution of the used bioclim variables in the niche modelling. Length and angle of the lines show the contribution of each environmental variable to the PC axes. Values of contribution of the bioclim variables for PC1 and PC2 are shown in Table S3.

The primary contributors to the differentiation along the PC1 axis, and thus main discriminators between species with dominant niches in warm and humid conditions versus those in colder and drier conditions, are mainly bio1 and bio18, as well as bio 7 with high negative contribution and bio12 with a slightly weaker contribution (Fig. 3c and Table S3). Bio15 shows a high contribution to the climatic differentiation between the cold adapted species along PC2 (Table S3).

### Species-specific and regional range shifts

Our analysis indicates a general expansion of all species/OTU ranges across all regions for both scenarios (Fig. 4a and Fig. S7). Major changes in species distributions in the maritime Antarctic are largely driven by an area expansion of *Lecidea atrobrunnea* (Fig. 4a and Fig. S8; Table S4) and by photobiont OTU *Tr*_I01 (Fig. 4; Table S4). Species such as *Lecidea andersonii* and *Lecidea cancriformis*, are also predicted to occupy suitable areas in this region in the future, although the area loss of *Lecidea cancriformis* exceeds the area gain (Fig. 4b and Table S4), which results in an overall area reduction. The future distribution of those species is predicted to be primarily located in the fragmented parts of the maritime Antarctic (mainly on Alexander Island), whereas the major northwestern coastal regions of maritime Antarctica represent the core area of the expansion of *Lecidea atrobrunnea* and *Tr*_I01 (Fig. S8). Furthermore, on the photobiont side, OTUs *Tr*_A02 and *Tr*_S18 are predicted to slightly expand their potential niche areas in the maritime Antarctic, which is also highly fragmented.

**Figure 4:**
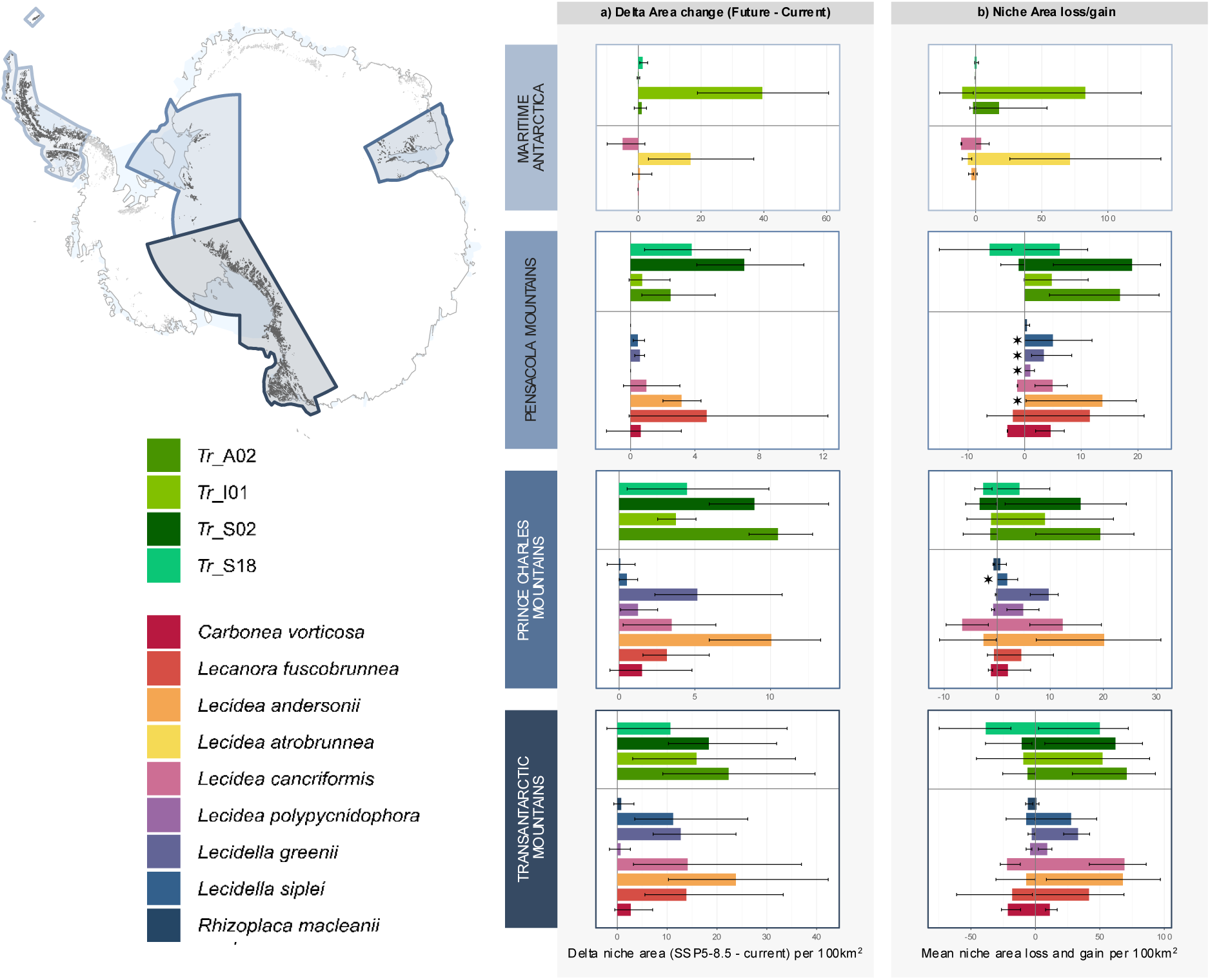
Change of probable niche area (mean of probable niche area for each GCM per 100km^2^) per mycobiont species/photobiont OTU for each selected region for SSP5-8.5: Maritime Antarctica, Pensacola Mountains, Prince Charles Mountains and, Transantarctic Mountains. a) Total area change (SSP5-8.5 area – current area) and b) gained and lost niche area and per mycobiont species/photobiont OTUs for each bioregion. In b) the bars above x - axis (0) depict the absolute gain of probable niche area (niche expansion), and bars below the x-axis the absolute loss of probable niche area (niche reduction). The four bioregions are shown in the upper left map of Antarctica and are colour coded in the boxes of the respective plots. Mycobiont species and photobiont OTUs are colour coded according to the legend in the left lower corner. Error bars represent the minimum and the maximum value, resulting from different GCMs used (see Methods: Climatic Data and Models). Bars with asterisks present the mycobiont species/photobiont OTUs, which are predicted to establish an entirely new niche in this area. Please note the different scales per region for a) and b). The respective analysis with SSP1-2.6 and SSP5-8.5 combined can be found in Fig. S7, and the absolute values of area change in Table S4.

In the three studied regions of continental Antarctica, niche area expansion exceeds niche area reduction for all mycobiont species and photobiont OTUs (Fig. 4a). The Transantarctic Mountains showed the greatest absolute area gain of new species niches (Fig. 4a and Table S4). Any mycobiont species or photobiont OTUs with a current probable niche in the Transantarctic Mountains region are predicted to expand its niche considerably (Fig. 4a), as the newly gained areas exceed the lost areas due to climate change (Fig. 4b). A similar pattern emerges in the Prince Charles Mountains, where each mycobiont species/photobiont OTU, which is already occurring expands its niche (Fig. 4a; Table S4). Additionally, *Lecidella siplei,* which was not predicted to have a current niche in this region, is increasing its future distributional area in this region (Fig. 4a). It is expected to establish most of the new areas (in the Pensacola Mts. and Prince Charles Mts.; Fig. 4a). Proportional to the currently occupied area, the Pensacola Mountains exhibit the highest increase in probable niche area expansion (Table S4). This trend is particularly evident for the mycobiont species *Lecidea andersonii, L. polypycnidophora, Lecidella greenii, L. siplei,* and *Rhizoplaca macleanii*, and the photobiont OTU *Tr*_A02, which are predicted to establish new niches in this region (Fig. 4a and Table S4). Additionally, the already existing species *Carbonea vorticosa, Lecanora fuscobrunnea,* and *Lecidea cancriformis* are also expected to expand their probable niches within the area (Fig. 4a).

Overall, the presented future projections for SSP5-8.5 and SSP1-2.6 indicate a general trend of major niche expansion, which notably exceeds the loss of niche area due to climate warming (Fig. 4a, Fig. S7). The extent of niche expansion and reduction is greater for SSP5-8.5, while SSP1-2.6 shows smaller changes in niche area for mycobiont species and photobiont OTUs (Figure S7 and Table S4).

The projected probabilities of occurrence show no consistent patterns across different GCMs (Fig. S8a and b). Predictions with MPI-ESM1-2-HR show fundamentally different outcomes than predictions with other used GCMs. Especially when projecting the mycobiont species *Lecanora fuscobrunnea, Lecidea andersonii* and *Lecidella siplei* and all photobiont OTUs using MPI-ESM1-2-HR there is a prediction for a major area reduction instead of the predicted area expansion when using all other GCMs or the Ensemble Mean (Fig. S8b).

Future projections for SSP1-2.6 and SSP5-8.5 scenarios indicate not only an overall trend of area expansion for lecideoid lichens species (Fig. 5a) and their photobionts but also a distinct spatial shift of those predicted ranges, with areas of overlapping mycobiont nichesshifting from coastal areas toward the inland (Fig. 5b). Notably, SSP5-8.5 regions with 3 and 4 overlapping mycobiont niches (Fig. 5a; Table S5) show the largest shift towards the inland. The zones with the maximal number 8 of overlapping probable niches experience the smallest inwards shift (Fig. 5a, Table S5) in SSP5-8.5 and the second smallest for SSP1-2.6 (Fig. 5a, Table S5). Overall, a significant trend emerges, where areas with higher species richness are predicted to shift inland, while a decline in species richness is to be expected in coastal regions. For SSP1-2.6 a similar, yet weaker pattern can be found (Fig. 5a).

**Figure 5:**
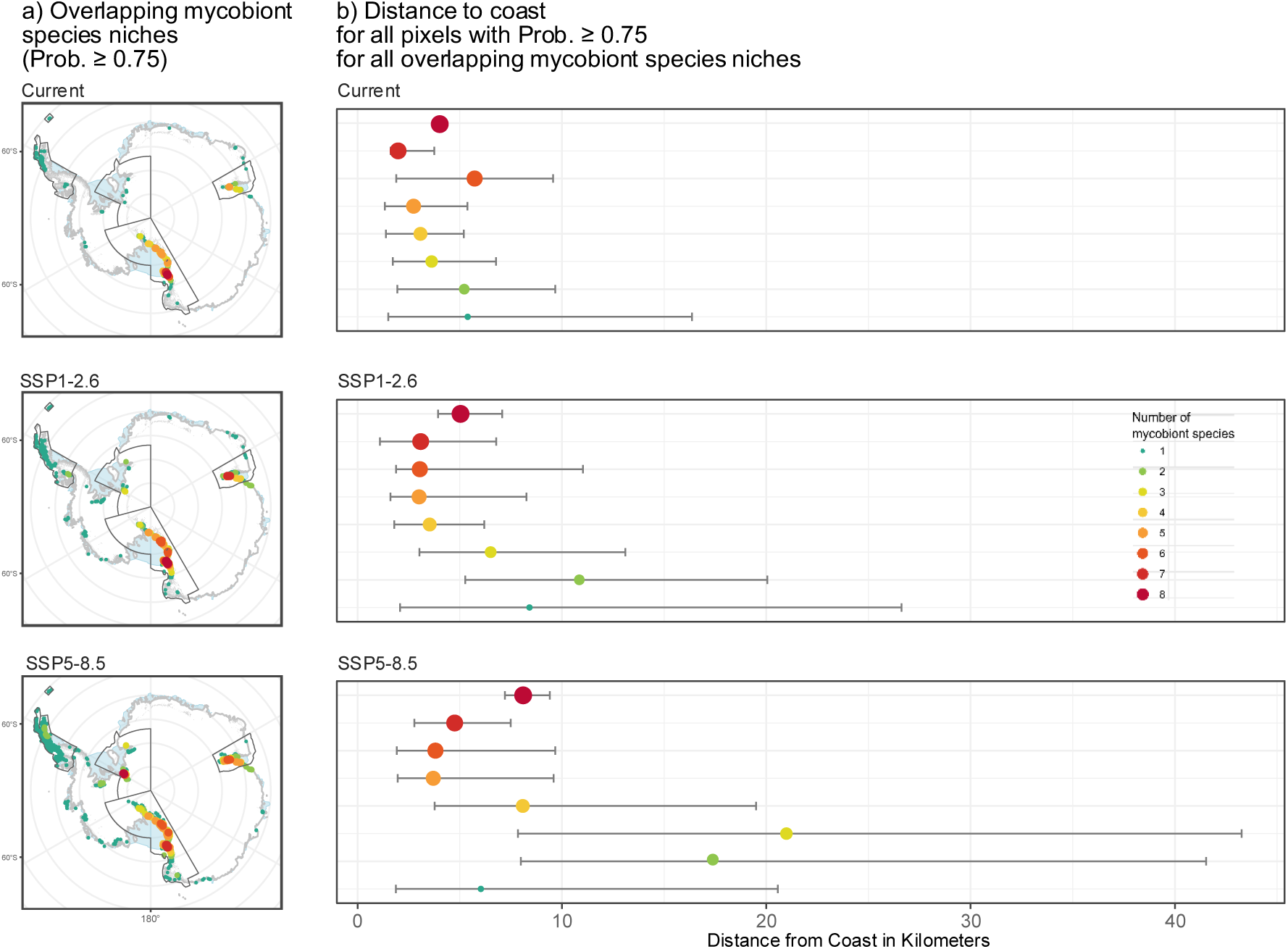
(a) Overlapping mycobiont species niches (Prob. ≥ 0.75) across the selected four bioregions. Note that each dots represents one pixel, which is not to scale. The size of the dots is proportional to the number of overlapping mycobiont species niches. (b) Distance from the coast for each pixel with ensemble mean probability ≥ 0.75 of mycobiont species for current conditions, the SSP1-2.6 and the SSP5-8.5 scenario. Colours and size of the dots indicate the number of overlapping mycobiont species niches.

## Discussion

The biogeographical and bioclimatic characteristics of the Antarctic continent only allow life on a scattered and very reduced scale. Nevertheless, there is a diverse flora of lecideoid lichens occurring as pioneer vegetation in almost all habitat types of the Antarctic continent (e.g. Wagner 2021, Ruprecht 2010, Castello 2003, Hertel 2007, Andreev 2023). The extensive and circum-Antarctic data set of this study, comprising 637 lecideoid lichen specimens revealed a total of 34 lecideoid mycobiont species for Antarctica reflecting the substantial progress made over the past 50 years in documenting this well-studied group, which had previously been recorded on a much smaller scale (Castello, 2003; Hertel, 2007; Ruprecht et al., 2020). Nevertheless, three further species of the genus *Lecidella* were detected on Livingston Island (S62°) by molecular methods (Fig. S5) but have yet to be described morphologically. The level of endemism for lecideoid lichens in Antarctica is quite high, with 13 species and 8 species from the southern polar regions, although the numbers keep changing depending on the level of species studied worldwide (Castello, 2003; Hale et al., 2019; Hertel, 2007; Øvstedal & Smith, 2001; Ruprecht et al., 2012b).

Most of the photobiont OTUs of the genus *Trebouxia* identified here have also been documented in previous studies. Earlier documentation of these OTUs, however, has predominantly been limited to studies conducted in the Transantarctic Mountains (Pérez-Ortega et al., 2023; Ruprecht et al., 2012a; Wagner et al., 2020; Wagner et al., 2021). Interestingly, almost all of these OTUs are characterized by a widespread and cosmopolitan distribution (Medeiros et al., 2024; Muggia et al., 2020; Ruprecht et al., 2020). One notable exception is the photobiont OTU *Tr*_S18, which has so far only been recorded in Antarctica and is therefore still considered endemic to this continent (Ruprecht et al., 2020). Despite the cosmopolitan nature of most photobionts, certain OTUs - particularly *Tr*_S02 and *Tr*_I01 - exhibit distinct Antarctic subgroups – most likely a consequence of geographic isolation and specialized adaptation processes, driven by the uniquely harsh climatic conditions prevailing in Antarctica (Ruprecht et al., 2012a; Ruprecht et al., 2020; Wagner et al., 2021).

Considering that lichens respond sensitively to changes in its occurrences of (macro-) climatic conditions (Mallen-Cooper et al., 2023; Sancho et al., 2019; Wagner et al., 2020), it seems natural to use them as model organisms in order to evaluate climate-induced changes in species diversity and range dynamics based on their established climatic niches. With climate warming, the cold desert ecosystems of Antarctica undergo significant changes (Colesie et al., 2023; Convey et al., 2012; McGaughran, Laver, & Fraser, 2021), which will not only alter but also affect the distributions of mycobiont species and photobiont OTUs.

### Niche expansion and inland-directed shift caused by future climate change

While mountainous regions worldwide, such as the Himalayas, where suitable habitats for cold-adapted species will be limited in the future (Manish et al. (2016), climate change in Antarctica is expected to create additional opportunities for colonization, as both previously uncolonized inland areas and newly exposed rock outcrops become climatically suitable for terrestrial organisms. The predicted expansion of Antarctic lecideoid lichens is in line with the global projections of lichen ranges by Mallen-Cooper et al. (2023), who also stated a major increase in niche area for boreal - arctic lichens. Further studies predict an inland directed shift of Antarctic endolithic communities (Zucconi et al., 2016) and an overall expansion of Antarctic native insects (Contador et al., 2020) due to climate change.

This study strongly suggests that the distribution of lecideoid lichens will expand significantly in response to climate change under the two scenarios SSP1-2.6 and SSP5-8.5. Furthermore, they also observed that the newly gained niche area exceeds and therefore overcompensates for the niche area losses in future scenarios, which our study also predicts for Antarctic lichens.

The species-specific distribution patterns (Fig. 3), and consequently their responses to climate change (Fig. 4), are primarily shaped by the climatic divide between maritime and continental Antarctica. Therefore, predicted future range shifts highlight ‘winners’ and ‘losers’ among Antarctic terrestrial vegetation under climate change. While some species may benefit from newly exposed areas, others are at risk of significant habitat loss.

Mycobiont species and photobiont OTUs (i.e. *Lecidea atrobrunnea* and *Tr*_I01) that are predominantly distributed in milder and wetter areas will benefit most from climate change (‘winners’), mainly due to milder temperatures, increased precipitation and the expansion of ice-free areas in maritime Antarctica (Fig. 4a, Fig. S7 and Table S4), which is predicted to have the highest rate of melting (Lee et al., 2017). However, besides the climatic favourable changes, it can be expected that the interspecific competition increases with climate warming as vascular vegetation and invasive species will expand especially over maritime Antarctica (Cannone et al., 2022; Chown et al., 2012; Hughes et al., 2020; Pertierra et al., 2017) presenting an additional threat for highly cold - adapted, slow-growing species like lecideoid lichens. Conversely, we predict a substantial reduction in probable niche area for *Carbonea vorticosa, Lecidea andersonii*, and *Lecidea cancriformis* under the high-emission scenario SSP5-8.5. These species are specialized for cold, arid conditions - climatic niches that are expected to become limited across much of maritime Antarctica, even under the more optimistic SSP1-2.6 scenario. Under SSP5-8.5, it is assumed that their preferred environmental conditions will have nearly all but disappeared.

Continental Antarctica shows a distinct pattern in terms of future range dynamics, with relatively minor differences among the individual regions (Fig. 4a, b) and an overall trend toward range expansion under climate change. At the species level, no strongly divergent or species-specific response patterns are evident; rather, a general expansion of suitable niche area is predicted. Each mycobiont species, except *Lecidea atrobrunnea*, and all photobiont OTUs are expected to expand their niche area mainly in the large continental rock outcrop areas (Fig. 4a), shifting from coastal regions to the inland (Fig. 5 and Fig. S8).

When comparing the three continental bioregions, nearly all species are predicted to experience a significant expansion of suitable habitat. In particular, the Pensacola Mountains are expected to show the greatest increase in newly available niches, while simultaneously exhibiting only minor habitat losses. Proportionally to its area, the Pensacola Mountains may exhibit the highest niche expansion among the regions examined. To date, the Pensacola Mountains remain less studied compared to the Prince Charles Mountains, and particularly the widely surveyed Transantarctic Mountains. To account for these differences, we calculated future range dynamics based on the predicted current probable niches. Species that are currently widespread across continental Antarctica are likely to already occur in the Pensacola Mountains, despite the absence of records. With increased sampling efforts, it seems possible that the distribution patterns observed in the Transantarctic Mountains can also be detected in the Pensacola Mountains, albeit at a different scale than in the Prince Charles Mountains.

In the Transantarctic Mountains, where many species currently have their largest niche areas, future range expansion is expected primarily inland, while reductions will mostly affect coastal zones. The predicted inland shift is also consistent with previous observations for endolithic communities (Yung et al., 2014; Zucconi et al., 2016). The shift will occur on already exposed substrates in the still uncolonized inland, as a certain degree of weathering is necessary for these sites to be colonizable (Friedmann, 1982; Zucconi et al., 2016). Particularly, south-facing slopes, influenced by colder temperatures, katabatic winds, and a pronounced rain-shadow effect, are currently less likely to be colonized (Lenaerts et al., 2016; Nicolas & Bromwich, 2014; Yung et al., 2014). These conditions are expected to persist despite increasing coastal precipitation due to climate change. Those inland areas may serve as future climatic refugia, as coastal regions are likely to experience more rapid and pronounced environmental changes (Zucconi et al., 2016; McGaughran et al., 2021; Tewari et al., 2022).

Directed shifts in species distributions are often observed as a result of climate change (Chen et al., 2011), particularly latitudinal range shifts are highly pronounced in Arctic regions (Mallen-Cooper et al., 2023; van Beest et al., 2023). Equivalent to latitudinal shifts in the Northern Hemisphere, range shifts in Antarctica are directed inland. The climatic gradient in Antarctica is not proportional to latitude, with regions around 80°S being the driest and coldest (Colesie et al., 2014; Wagner et al., 2021). With increasing latitude towards the South Pole, climatic conditions will become comparatively milder again, in contrast to the northern Arctic regions.

### Constraints of Species Distribution Models (SDMs)

When interpreting predicted future species distributions, it is crucial to acknowledge that all environmental inputs are derived from climate models, which are themselves projections. This inherent uncertainty must be considered when interpreting the results. To illustrate the variability among models, we provide individual ensemble model (EM) projections for each General Circulation Model (GCM) in Fig. S8a and b. These projections reveal substantial differences in predicted range dynamics. Notably, the projections with the MPI-ESM1-2-HR model consistently forecast an overall reduction in niche area across all species and OTUs, in contrast to the general pattern of range expansion predicted by the other GCMs. Furthermore, the minor range expansions predicted for *Lecidea andersonii* and *Tr*_S18 in maritime Antarctica may reflect model artifacts, potentially driven by the tendency of GCMs to vary in the projected precipitation in maritime Antarctica and coastal areas. These findings highlight the necessity of cautious interpretation when projecting future distributions under climate change scenarios.

In addition to macroclimatic factors such as temperature and precipitation/atmospheric humidity, the ab initio establishment of a lichen depends as well on suitable microhabitats, where sheltered conditions prevail (Colesie et al., 2023; Green et al., 2011; Matos et al., 2024). Rock-dwelling lichens depend on moisture retained in weathered rock, from the air and by melting snow. Due to this reliance, broader climatic changes that alter atmospheric moisture levels can have profound effects on their distribution. On the Antarctic continent, the loss of sea ice and alterations in wind systems due to climate change are estimated to increase atmospheric humidity (Lenaerts et al., 2016; Medley & Thomas, 2019; Winkelmann et al., 2012). Along with continent-wide increasing precipitation (Vignon et al., 2021) and temperatures contributing to surface melt (Green et al., 2011; Lenaerts et al., 2016), these changes are expected to enhance the availability of water and humidity, potentially creating favourable conditions for future micro habitats. Additionally, increased liquid precipitation may initially promote lichen growth as result in a shift in species composition, since some lichens are outcompeted by those better adapted to these wetter conditions (Green et al., 2011). These warm-adapted species typically exhibit higher growth rates (Sancho et al., 2019), allowing them to expand more rapidly than slower-growing crustose lichens. However, additional precipitation can also have negative effects on lichen coverage: A major increase in snowfall leads to reduced activity and a subsequent decline in lichen coverage due to the insulating effect of the snowbank, which was observed and stated by Sancho et al. (2017) and Green et al. (2011).

Furthermore, when assessing species range dynamics, it is important to consider whether colonialization, niche area expansion and establishment are possible, depending on the dispersal ability of the species, the local microclimatic conditions and the intra-as well as interspecific competition on site. Especially for terrestrial life in Antarctica, the dispersal ability is key, as the patches of ice-free areas are fragmented by vast snow-covered areas, which can act as dispersal barriers (Koerich et al., 2023; Lee et al., 2022b). Based on the monitoring of fungal in the air it can be considered that lecideoid lichens are effectively dispersed via spores, soredia, and thallus fragments at long distances by wind (Muñoz et al., 2004). For future niche predictions, we therefore assumed, referring to Mallen-Cooper et al. (2023), that reaching a newly suitable habitat might be possible around the current and future ice-free areas across Antarctica. However, a recent study of Neme, England, and McC. Hogg (2022) indicates circumpolar weakening of near-surface winds under high to moderate emission scenarios (SSP5-8.5 and SSP2-4.5). As this is expected to occur mainly during the summer season, it may negatively affect the circumantarctic dispersal of lichen species and thus impede climate-driven niche expansion.

## Conclusion

Our findings confirm the high extent of diversity of lecideoid lichens as well as the further discovery of new species. Despite predicted habitat loss due to climate warming, range expansions are expected to exceed niche reduction for most species, driven largely by a shift toward inland regions. While maritime Antarctica may experience reductions in suitable habitat, continental areas are likely to gain new colonizers as environmental conditions become more favourable.

The extent of future range expansion will depend on the dispersal capacity of these species and the availability of microhabitats offering sufficient moisture and stability. Although effective wind dispersal suggests high potential, successful establishment remains constrained by microclimatic variability. Increasing surface melt may create new ecological opportunities, but local differences in temperature, wind exposure, and precipitation will continue to shape species distributions.

Integrating macro- and microclimatic modelling with species-specific physiological constraints will be essential to improve predictions of Antarctic vegetation dynamics. Understanding these patterns offers key insights into the resilience of cryptogamic organisms under climate change and contributes to broader assessments of biodiversity responses in polar ecosystems.

## Supporting information

Supplementary Information Goetz et al. 2025

## Acknowledgments

We thank Hans-Peter Comes for his valuable advice and support, Matthias Affenzeller (Department of Environment and Biodiversity, University of Salzburg) for support in the niche modelling, and Manfred Mittelböck (Department of Geoinformatics – Z_GIS, University of Salzburg) for support with climate layer preparation.

The study was funded in whole or in part by the Austrian Sciences Fund (FWF) 10.55776/P26638 and 10.55776/P35512, by the project CRYPTOCOVER (Spanish Ministry of Science CTM2015-64728-C2-1-R) as well as by the Russian Antarctic Expedition and by the Komarov Botanical Institute of the Russian Academy of Sciences (grant number 121021600184-6). Additionally, the last two authors gratefully acknowledge the support of the IDA Lab Salzburg (20102/F2300464-KZP).

## Author Contributions

Funding acquisition: UR, RRJ, WT; Conceptualization AG, UR; Formal Analyses AG, UR; Investigation AG, UR; Methodology AG, LM, WT; Resources MA, UR, LS; Visualisation AG; Original Draft Preparation AG, UR; Review & Editing MA, AG, RRJ, LM, UR, LS, WT.

## Data availability

Sequences were uploaded to and NCBI database, the complete dataset can be found in the Supplementary Information.

