## Supplementary Information Goetz et al. 2025 for "Future Range Shifts and Diversity Patterns of Antarctic Lecideoid Lichens Under Climate Change Scenarios"

### Supporting Information

**Fig. S1** Prince Charles Mountain Region with sampling sites of lichen and geographic information based on Andreev (2023)

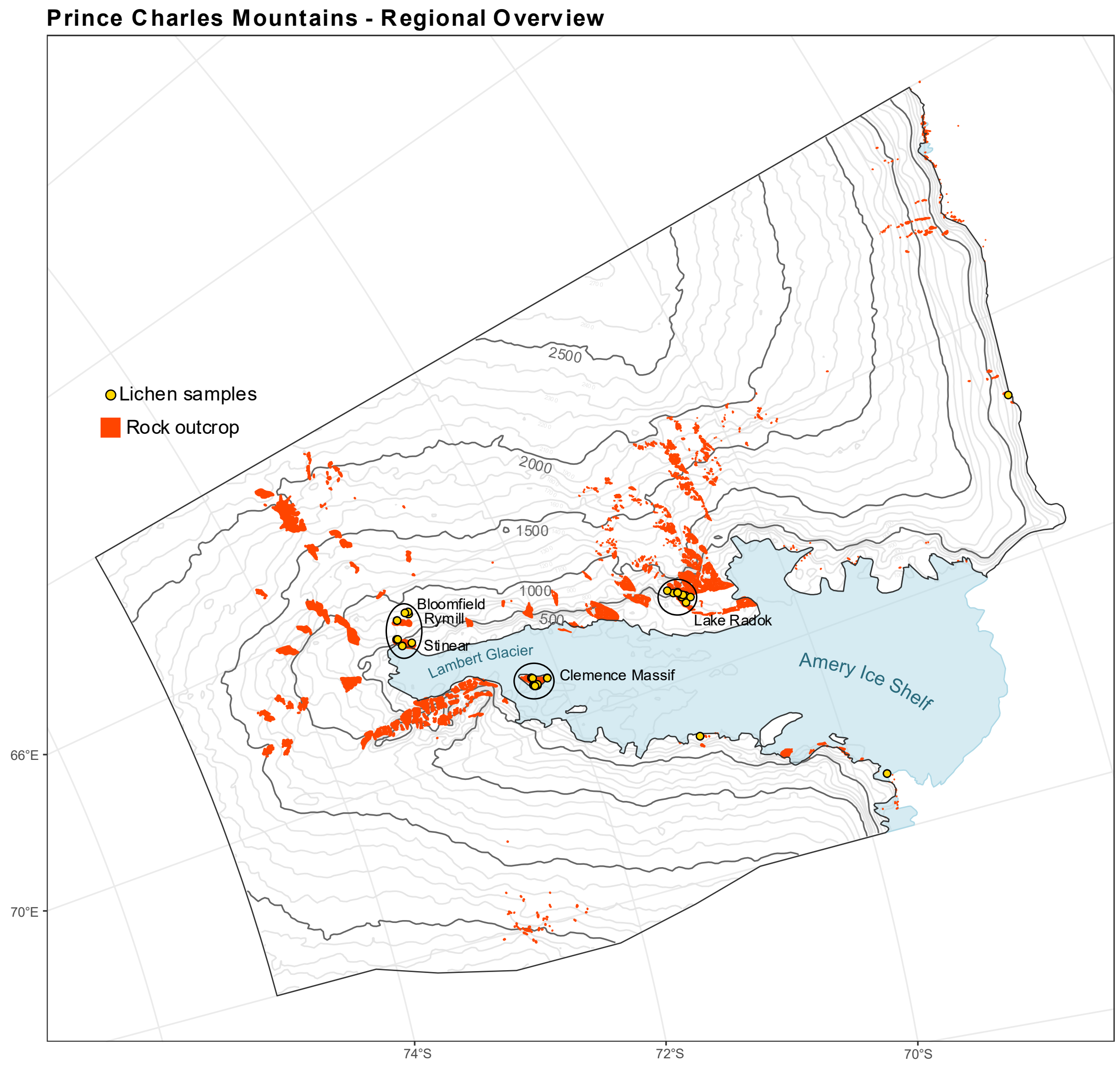

**Fig. S2** Correlation Clusters (in grey boxes) of all bioclim variables based on the current rock outcrop (Medium resolution vector polygons of Antarctic rock outcrop of the SCAR ADD), downscaled to a resolution of 2380m to 3680m in EPSG3031. Coloured bioclim variables where chosen based on the requirements of lecideoid lichens under polar conditions. Reddish coloured boxes correspond to temperature related and blueish coloured boxes to precipitation related bioclim variables.

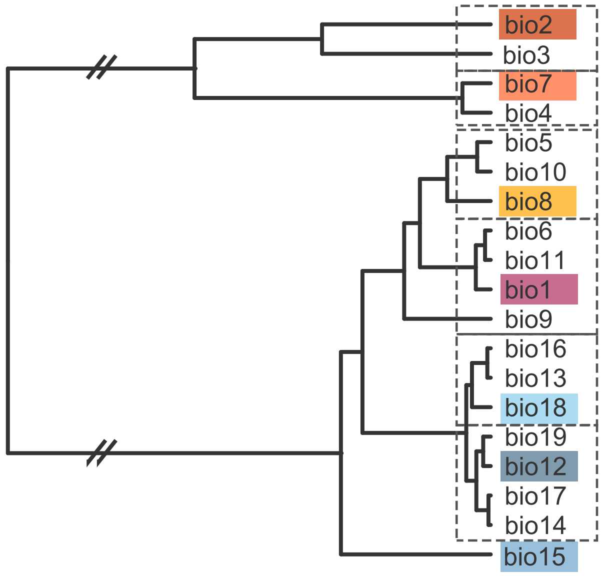

**Fig. S3** Variable Importance (var.imp) of the environmental predictors (bioclim variables) based on the built BIOMOD ensemble model per mycobiont species and photobiont OTU. Reddish coloured boxplots correspond to temperature related and blueish coloured boxplots to precipitation related bioclim variables.

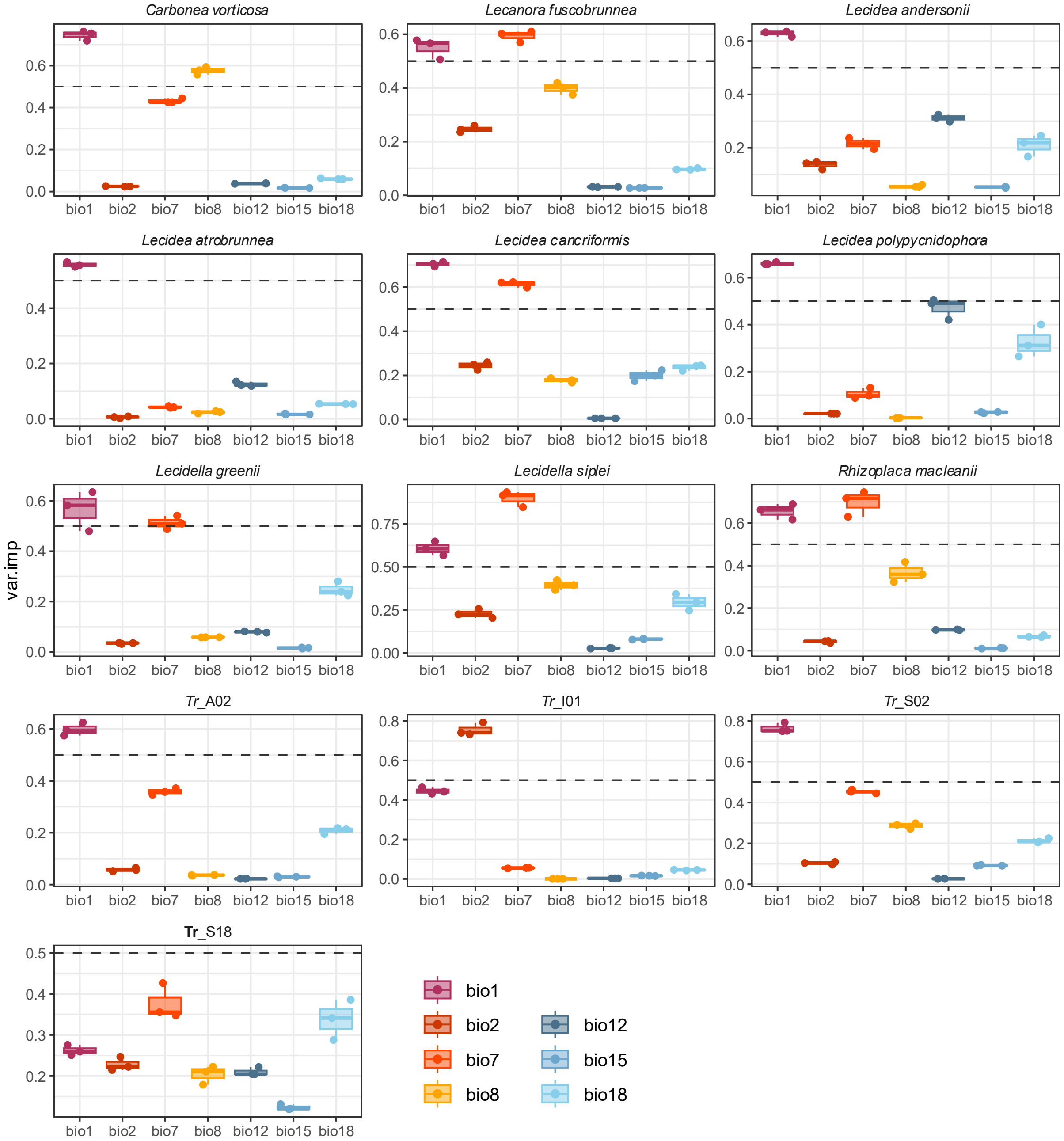

**Fig. S4** Diversity of all sampled (a) mycobiont species and (b) photobiont OTUs for all bioregions by Terauds and Lee (2016). In bold the mycobiont species/photobiont OTUs which were used in the downstream niche modelling. Size of the donut charts are proportional to the number of mycobiont species/photobiont OTU in the respective region.

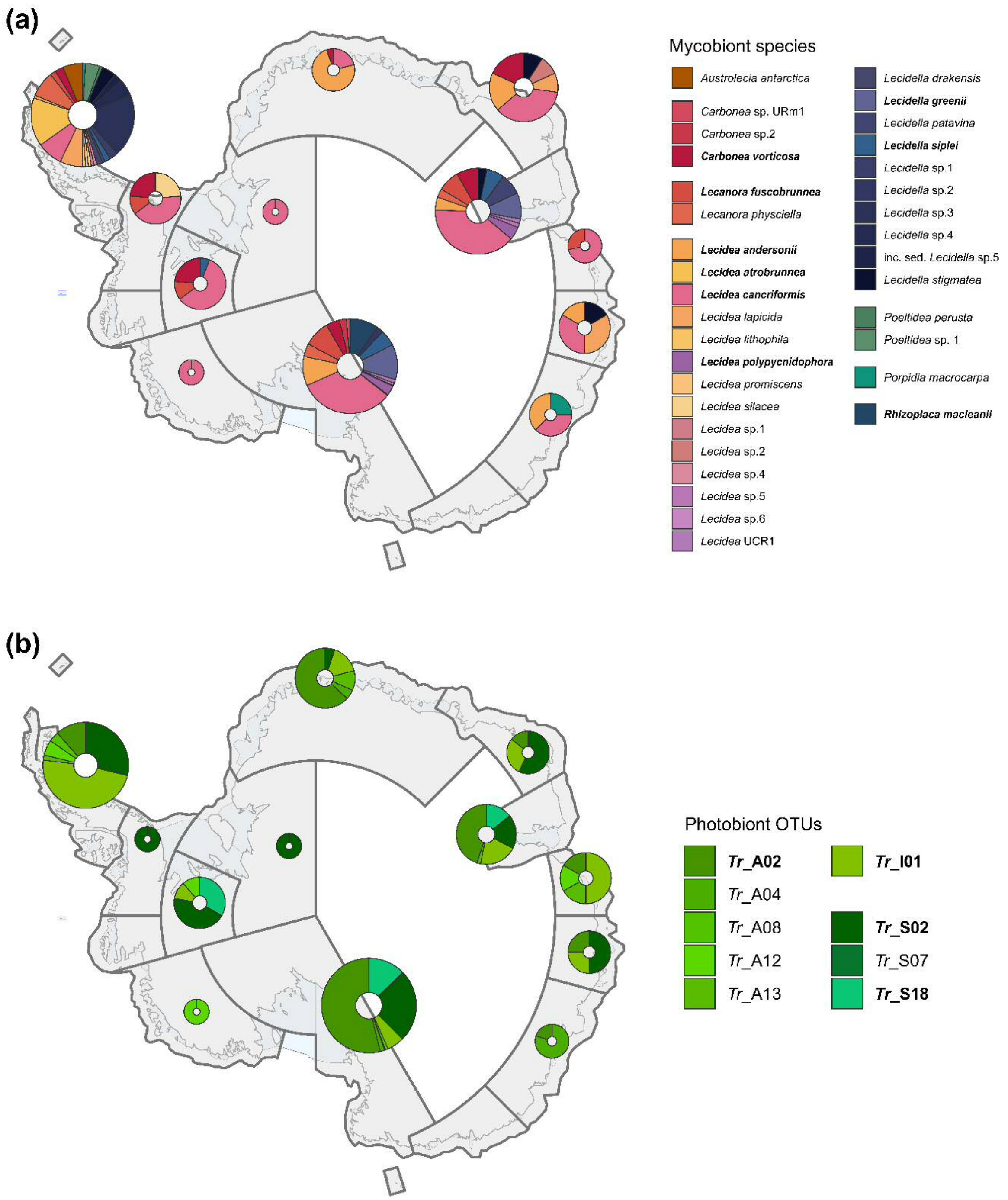

**Fig. S5** Phylogenetic analysis of ITS sequences of the genus Lecidella with the newly developed clades Lecidella sp. 3, L. sp. 4 and inc. sed. L. sp. 5 integrated in the species concepts of Zhao et al. (2016)and Ruprecht et al. (2020). Maximum likelihood (ML) bootstrap values SH-aLRT ≥

80%/UFboot ≥ 95%. All Antarctic accessions are highlighted in bold.

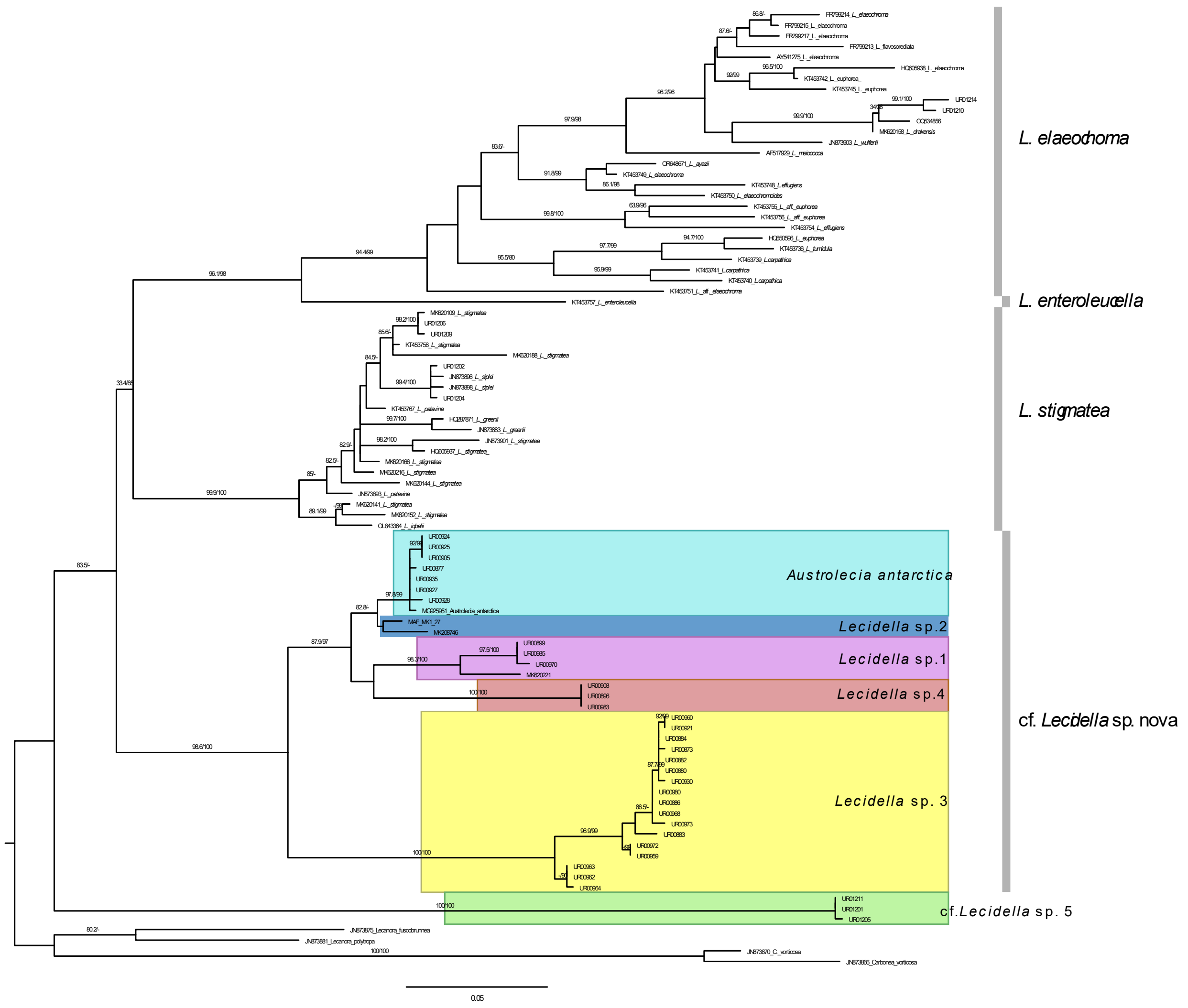

**Fig. S6:** Current Niches per pixel (filtered for all assigned "-2" and "-1" via RangeDiff() in biomod2) of all a) mycobiont species and b) photobiont OTU in maritime Antarctica. Single pixels shown as points for better visibility but note that point sizes are not to scale.

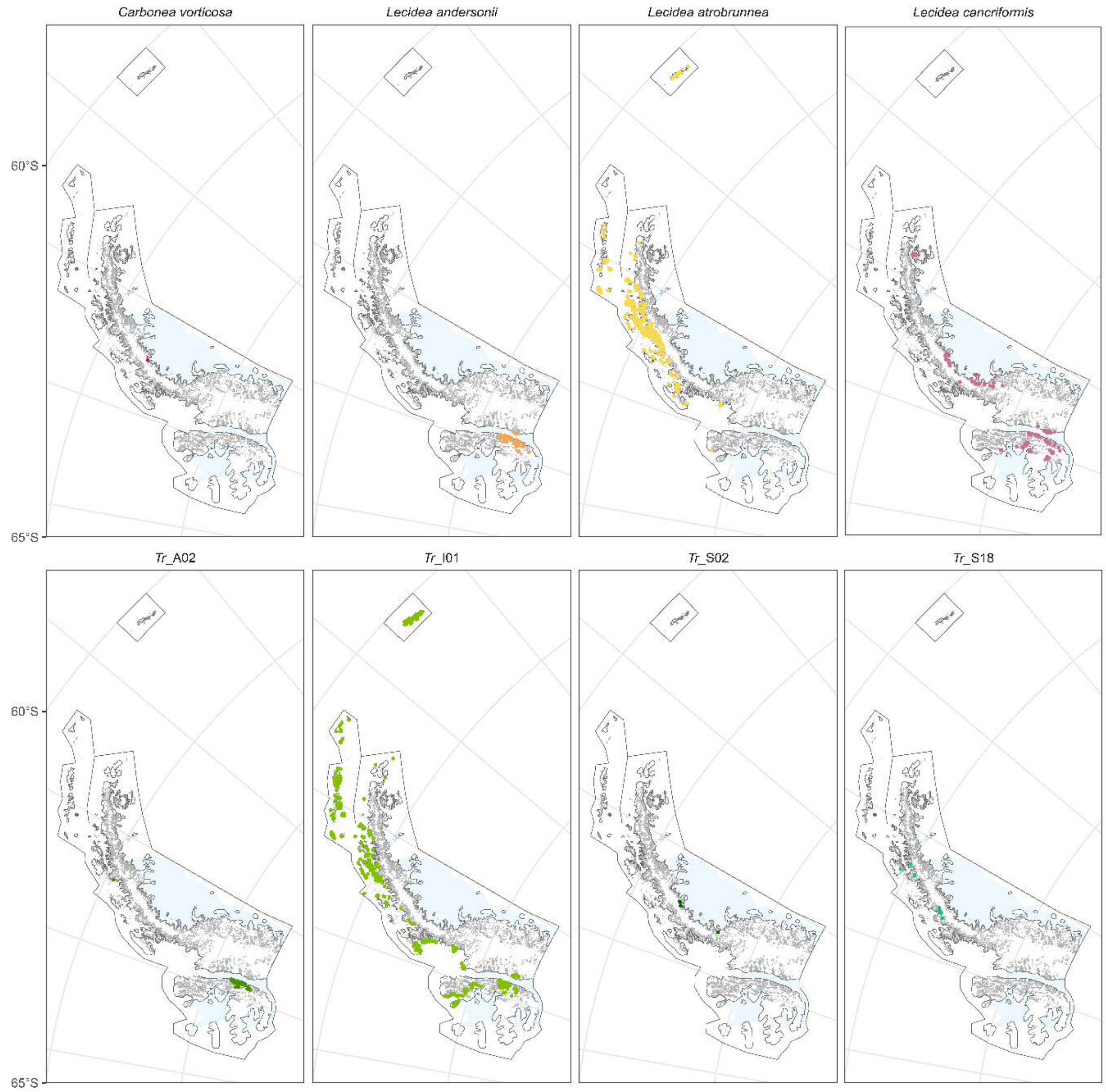

**Fig. S7** a) Niche area change (SSP1-2.6 area/SSP5-8.5 area – current area) and b) niche area gain/loss per mycobiont species/photobiont OTU for each selected region (Maritime Antarctica, Pensacola Mountains, Prince Charles Mountains and Transantarctic Mountains) for SSP1-2.6 and SSP5-8.5. Bars above 0 show for a) an overall niche expansion and for b) the absolute gain of probable niche area), and bars below 0 show for a) an overall niche area reduction and for b) the absolute loss of probable niche area. Mycobiont species and photobiont OTUs are colour coded according to the legend. Light colours for SSP1-2.6 and darker shades for SSP5-8.5. Error bars represent the minimum and the maximum value, resulting from different GCMs used (see Methods: Climatic Data and Models). Bars with Asterisks present the mycobiont species, which are predicted to establish an entirely new niche in this area.

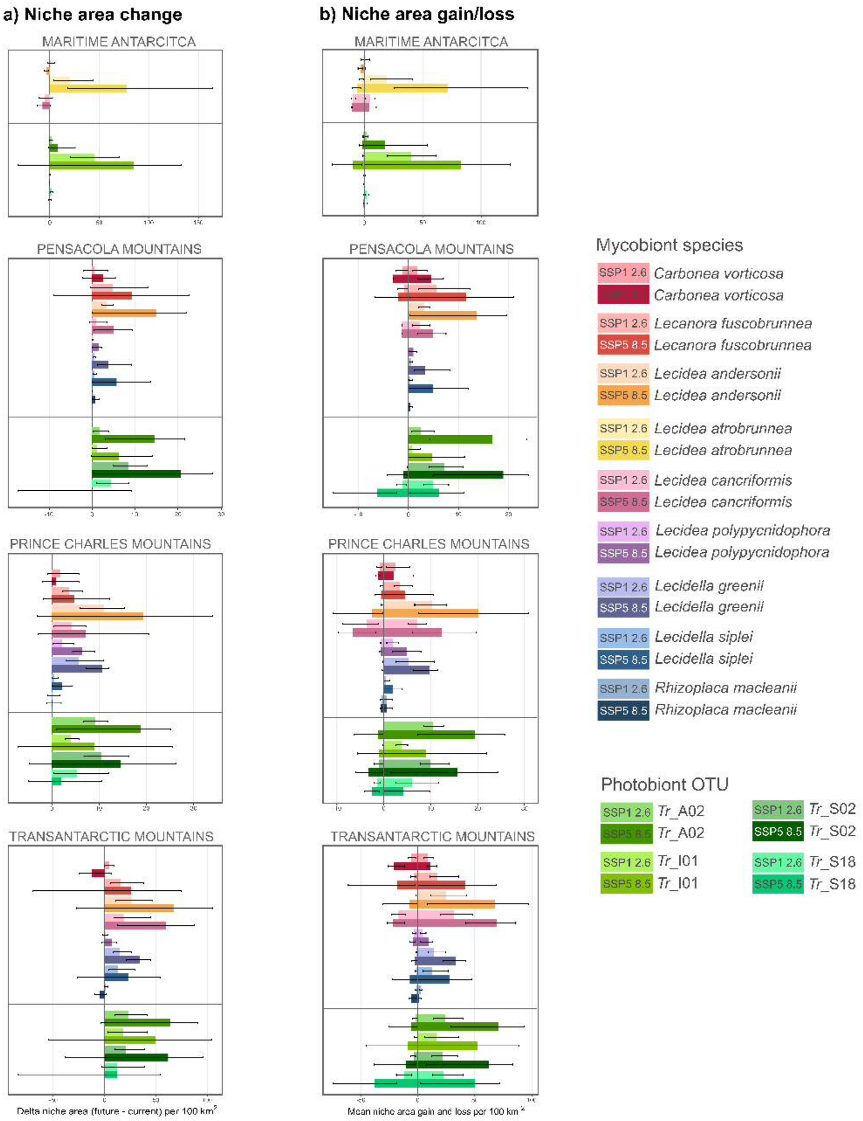

a) Niche area change
b) Niche area gain/loss
MARITIME ANTARCITCA
PENSACOLA MOUNTAINS
PRINCE CHARLES MOUNTAINS
TRANSANTARCTIC MOUNTAINS
Delta niche area (future - current) per 100 km2
Mean niche area gain and loss per 100 km2
Mycobiont species
SSP1 2.6 Carbonea vorticosa
Carbonea vorticosa
SSP1 2.6 Lecanora fuscobrunnea
SSP5 8.5 Lecanora fuscobrunnea
SSP1 2.6 Lecidea andersonii
SSP5 8.5 Lecidea andersonii
SSP1 2.6 Lecidea atrobrunnea
SSP5 8.5 Lecidea atrobrunnea
SSP1 2.6 Lecidea cancriformis
SSP5 8.5 Lecidea cancriformis
SSP1 2.6 Lecidea polypycnidophora
SSP5 8.5 Lecidea polypycnidophora
SSP1 2.6 Lecidella greenii
SSP5 8.5 Lecidella greenii
SSP1 2.6 Lecidella siplei
SSP5 8.5 Lecidella siplei
SSP1 2.6 Rhizoplaca macleanii
SSP5 8.5 Rhizoplaca macleanii
Photobiont OTU
SSP1 2.6 Tr_A02
Tr_A02
SSP1 2.6 Tr_I01
SSP5 8.5 Tr_I01
SSP1 2.6 Tr_S02
SSP5 8.5 Tr_S02
SSP1 2.6 Tr_S18
SSP5 8.5 Tr_S18

**Fig. S8:** a) Delta niche area change (SSP5-8.5 – current) for each GCM (GFDL-ESM4, IPSL-CM6A-LR, MPI-ESM1-2-HR, MRI-ESM2-0, UKESM1-0-LL) across mycobiont species/photobiont OTU across the four selected bioregions. b) Relation between the projected current probabilities of occurrence and future (SSP1-2.6 and SSP5-8.5) probabilities of occurrence for each pixel per mycobiont species and photobiont OTU (pooled).

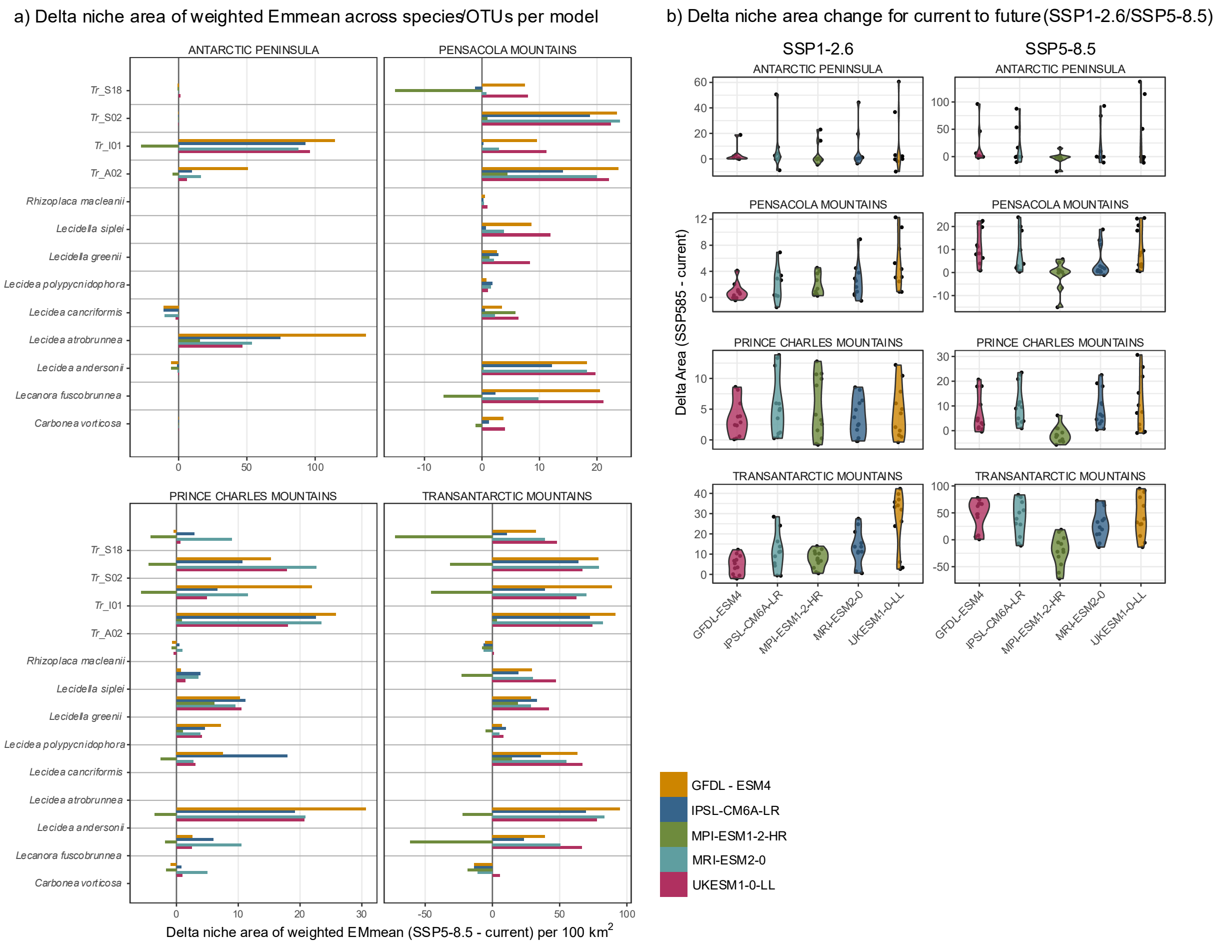

**Table S1** Species List with the Voucher ID, the mycobiont and associated Photobiont (if sequenced), the large-scale sample region, the altitude (alt), latitude (lat) and longitude (lon) of sampling location.

| | | | | | | | Accession Number | |
|---|---|---|---|---|---|---|---|---|
| **Voucher ID** | **Mycobiont species** | **Photobiont OTU** | **Region** | **altitude** | **latitude** | **longitude** | **mycobiont** | **photobiont** |
| UR00877 | *Austrolecia antarctica* | Tr_S02 | MA | 62 | -62.6651 | -60.3736 | PV788468 | - |
| UR00905 | *Austrolecia antarctica* | - | MA | 142 | -62.6666 | -60.392 | PV788469 | - |
| UR00924 | *Austrolecia antarctica* | - | MA | 383 | -62.6816 | -60.3422 | PV788470 | - |
| UR00925 | *Austrolecia antarctica* | - | MA | 383 | -62.6816 | -60.3422 | PV788471 | - |
| UR00927 | *Austrolecia antarctica* | - | MA | 383 | -62.6816 | -60.3422 | PV788468 | - |
| UR00928 | *Austrolecia antarctica* | - | MA | 383 | -62.6816 | -60.3422 | PV788473 | - |
| UR00935 | *Austrolecia antarctica* | - | MA | 443 | -62.6809 | -60.3431 | PV788472 | - |
| MAF_DP1_52 | *Carbonea sp. URm1* | Tr_A02 | CA | 370 | -85.5392 | -151.15 | MK208734 | MK226866 |
| T44626 | *Carbonea sp. URm1* | Tr_I01 | CA | 484 | -79.8819 | 159.3608 | JN873865 | JN204797 |
| T46719 | *Carbonea sp. URm1* | Tr_A02 | CA | 425 | -78.0378 | 163.8044 | MK970657 | MK970698 |
| T48803b | *Carbonea sp. URm1* | Tr_A02 | CA | 775 | -78.1483 | 163.6296 | MK970657 | MK970699 |
| T48806 | *Carbonea sp. URm1* | Tr_A02 | CA | 568 | -78.1417 | 163.6278 | T48806 | MK970699 |
| MAF_DP1_54 | *Carbonea sp.2 UR* | Tr_S02 | CA | 370 | -85.5392 | -151.15 | MK208735 | MK226868 |
| MAF_Sancho3 | *Carbonea sp.2 UR* | Tr_A04 | CA | NA | -83.7606 | 172.755 | MK208767 | JN204839 |
| T35647 | *Carbonea sp.2 UR* | Tr_S02 | CA | 344 | -79.8514 | 159.3408 | JN873866 | JN204751 |
| T48789 | *Carbonea sp.2 UR* | Tr_S02 | CA | 860 | -78.1519 | 163.7388 | MK970654 | MK970692 |

| | | | | | | | | |
|---|---|---|---|---|---|---|---|---|
| T48793b | *Carbonea sp.2 UR* | Tr_S02 | CA | 831 | -78.1529 | 163.7312 | MK970654 | MK970692 |
| T48817b | *Carbonea sp.2 UR* | Tr_A02 | CA | 546 | -78.1127 | 163.7845 | MK970655 | MK970702 |
| T48820 | *Carbonea sp.2 UR* | Tr_A02 | CA | 575 | -78.1137 | 163.7802 | MK970654 | MK970698 |
| T48825 | *Carbonea sp.2 UR* | Tr_A02 | CA | 498 | -78.0969 | 163.6914 | MK970654 | MK970699 |
| T48832 | *Carbonea sp.2 UR* | Tr_A02 | CA | 526 | -78.1119 | 163.8243 | MK970654 | MK970702 |
| AAS_Convey00510 | *Carbonea vorticosa* | Tr_S02 | CA | 860 | -75.8408 | -69.423 | - | PV788567 |
| AAS_Convey00518A | *Carbonea vorticosa* | - | CA | 856 | -75.73 | -71.73 | - | - |
| AAS_Convey00543B | *Carbonea vorticosa* | Tr_S02 | CA | 791 | -75.43 | -72.6 | - | PV788577 |
| AAS_Convey00603 | *Carbonea vorticosa* | - | CA | 1167 | -77.03 | -78.27 | - | - |
| AAS_Convey00630 | *Carbonea vorticosa* | - | CA | 1260 | -75.12 | -72.18 | - | - |
| AAS_Convey00671 | *Carbonea vorticosa* | - | CA | 1760 | -66.78 | -64.88 | - | - |
| AAS_Convey00674 | *Carbonea vorticosa* | Tr_S02 | CA | 1760 | -66.78 | -64.88 | - | PV788580 |
| AAS_Convey01139A | *Carbonea vorticosa* | - | CA | 1351 | -79.88 | -82.59 | - | - |
| AAS_Convey01141B | *Carbonea vorticosa* | Tr_S02 | CA | 1351 | -79.8745 | -82.5962 | - | PV788568 |
| AAS_Convey1173 | *Carbonea vorticosa* | | CA | NA | -79.8333 | -83.65 | - | JN204833 |
| AAS_Smith10093 | *Carbonea vorticosa* | Tr_S02 | CA | NA | -71.9988 | 25.0125 | PV788369 | PV788566 |
| BG_15 | *Carbonea vorticosa* | - | CA | NA | -79.8333 | 83.65 | - | - |
| LE_A0513404 | *Carbonea vorticosa* | Tr_S02 | CA | 100 | -70.8667 | 67.95 | - | - |
| LE_A060601 | *Carbonea vorticosa* | - | mA | NA | -62.2223 | -58.7742 | - | - |
| LE_A112306 | *Carbonea vorticosa* | - | CA | 34 | -67.6629 | 45.87585 | - | - |
| LE_A113302 | *Carbonea vorticosa* | Tr_S02 | CA | 40 | -67.6661 | 45.82648 | - | PV788581 |

| | | | | | | | | |
|---|---|---|---|---|---|---|---|---|
| LE_A130510 | *Carbonea vorticosa* | - | CA | 117 | -72.2236 | 68.80992 | - | - |
| LE_A131304 | *Carbonea vorticosa* | Tr_I01 | CA | 124 | -72.2142 | 68.82712 | - | PV788623 |
| LE_A133804 | *Carbonea vorticosa* | Tr_S02 | CA | 1082 | -72.2214 | 68.58423 | - | PV788569 |
| LE_A150306 | *Carbonea vorticosa* | Tr_I01 | CA | 265 | -72.1022 | 68.73928 | - | - |
| T33418 | *Carbonea vorticosa* | - | CA | 450 | -83.7608 | 172.7402 | - | - |
| T33418 | *Carbonea vorticosa* | - | CA | 450 | -83.7608 | 172.7403 | - | - |
| T33714 | *Carbonea vorticosa* | Tr_S02 | CA | 1180 | -77.6964 | 162.2599 | JN873872 | PV788600 |
| T35649 | *Carbonea vorticosa* | - | CA | 344 | -79.8514 | 159.3408 | - | - |
| T35650 | *Carbonea vorticosa* | Tr_S18 | CA | 344 | -79.8514 | 159.3408 | JN873867 | JN204752 |
| T35686 | *Carbonea vorticosa* | Tr_S02 | CA | 405 | -79.8417 | 159.3631 | JN873868 | PV788573 |
| T43031 | *Carbonea vorticosa* | Tr_A02 | CA | 598 | -77.5701 | 163.1573 | JN873869 | JN204789 |
| T44641 | *Carbonea vorticosa* | Tr_A04 | CA | 491 | -79.8656 | 159.3653 | JN873871 | JN204803 |
| T44655 | *Carbonea vorticosa* | Tr_S02 | CA | 436 | -79.8744 | 159.3389 | MK208787 | MK226915 |
| T44681 | *Carbonea vorticosa* | - | CA | 416 | -79.8497 | 159.8497 | - | - |
| T44693 | *Carbonea vorticosa* | Tr_S02 | CA | 620 | -79.7553 | 158.5225 | - | JN204810 |
| T46651 | *Carbonea vorticosa* | - | CA | 347 | -78.028 | 163.851 | MK970656 | - |
| T46718 | *Carbonea vorticosa* | Tr_A02 | CA | 442 | -78.0397 | 163.8067 | MK970656 | MK970698 |
| T48788 | *Carbonea vorticosa* | Tr_A02 | CA | 397 | -78.1199 | 163.684 | MK970656 | MK970699 |
| T48790a | *Carbonea vorticosa* | Tr_A02 | CA | 415 | -78.1203 | 163.6818 | MK970656 | MK970699 |
| T48823a | *Carbonea vorticosa* | - | CA | 401 | -78.0981 | 163.7096 | MK970656 | - |
| T48831 | *Carbonea vorticosa* | Tr_A02 | CA | 451 | -78.1113 | 163.8584 | - | MK970699 |

| | | | | | | | | |
|---|---|---|---|---|---|---|---|---|
| T48841b | *Carbonea vorticosa* | Tr_A02 | CA | 722 | -78.0681 | 163.8614 | - | MK970698 |
| T48877 | *Carbonea vorticosa* | Tr_A02 | CA | 385 | -78.0241 | 163.8996 | - | MK970698 |
| AAS_Convey00495A | *Lecanora fuscobrunnea* | - | CA | 860 | -75.83 | -69.37 | - | - |
| AAS_Convey00684B | *Lecanora fuscobrunnea* | Tr_S02 | CA | 1334 | -75.0162 | -71.4185 | - | PV788598 |
| AAS_Convey01111 | *Lecanora fuscobrunnea* | Tr_S02 | CA | 1359 | -79.85 | -82.92 | - | PV788603 |
| AAS_Convey01113 | *Lecanora fuscobrunnea* | Tr_A12 | CA | 1395 | -79.85 | -82.92 | - | PV788552 |
| LE_A040301 | *Lecanora fuscobrunnea* | Tr_I01 | CA | 20 | -69.4093 | 76.38333 | - | PV788625 |
| LE_A106102 | *Lecanora fuscobrunnea* | Tr_I01 | CA | 46 | -69.4083 | 76.4 | - | PV788633 |
| LE_A130902 | *Lecanora fuscobrunnea* | Tr_I01 | CA | 127 | -72.218 | 68.81952 | - | PV788624 |
| LE_A131202 | *Lecanora fuscobrunnea* | Tr_S02 | CA | 132 | -72.2149 | 68.82547 | - | PV788592 |
| LE_A131402 | *Lecanora fuscobrunnea* | Tr_A02 | CA | 166 | -72.2281 | 68.7841 | - | PV788493 |
| LE_A131901 | *Lecanora fuscobrunnea* | Tr_I01 | CA | 127 | -72.2124 | 68.82608 | - | - |
| LE_A133805 | *Lecanora fuscobrunnea* | Tr_A02 | CA | 1082 | -72.2214 | 68.58423 | - | PV788544 |
| MAF_DP1_02 | *Lecanora fuscobrunnea* | Tr_S02 | CA | 370 | -85.5392 | -151.15 | MK208710 | MK226845 |
| MAF_DP1_08 | *Lecanora fuscobrunnea* | Tr_A02 | CA | 370 | -85.5392 | -151.15 | MK208714 | MK226848 |
| MAF_DP1_20 | *Lecanora fuscobrunnea* | Tr_A02 | CA | 370 | -85.5392 | -151.15 | MK208720 | MK226853 |
| MAF_DP1_28 | *Lecanora fuscobrunnea* | Tr_S02 | CA | 370 | -85.5392 | -151.15 | MK208724 | MK226857 |
| MAF_DP1_32 | *Lecanora fuscobrunnea* | Tr_A02 | CA | 370 | -85.5392 | -151.15 | MK208726 | MK226859 |
| MAF_DP1_50 | *Lecanora fuscobrunnea* | Tr_A02 | CA | 370 | -85.5392 | -151.15 | MK208732 | PV788495 |
| MAF_DP1_51 | *Lecanora fuscobrunnea* | Tr_A02 | CA | 370 | -85.5392 | -151.15 | MK208733 | MK226865 |
| MAF_MK1_03 | *Lecanora fuscobrunnea* | Tr_A02 | CA | 859 | -83.7748 | 171.8276 | MK208752 | MK226884 |

| | | | | | | | | |
|---|---|---|---|---|---|---|---|---|
| MAF_MK1_40 | *Lecanora fuscobrunnea* | Tr_A02 | CA | 859 | -83.7748 | 171.8276 | MK208760 | MK226893 |
| MAF_MK1_56 | *Lecanora fuscobrunnea* | Tr_S02 | CA | 859 | -83.7748 | 171.8276 | MK208765 | MK2268970 |
| T33713 | *Lecanora fuscobrunnea* | Tr_S02 | CA | 1180 | -77.6964 | 162.2599 | - | JN204745 |
| T35542 | *Lecanora fuscobrunnea* | - | CA | 421 | -79.8381 | 159.3408 | - | - |
| T35544 | *Lecanora fuscobrunnea* | - | CA | 421 | -79.8381 | 159.3408 | JN873873 | - |
| T35664 | *Lecanora fuscobrunnea* | - | CA | 344 | -79.8514 | 159.3408 | JN873874 | - |
| T42994 | *Lecanora fuscobrunnea* | Tr_S18 | CA | 903 | -79.8786 | 157.5389 | GU170839 | MK226904 |
| T44628 | *Lecanora fuscobrunnea* | Tr_S02 | CA | 498 | -79.8653 | 159.3519 | JN873875 | PV788574 |
| T44632 | *Lecanora fuscobrunnea* | Tr_I01 | CA | 490 | -79.8683 | 159.3519 | MK208774 | JN204800 |
| T44650 | *Lecanora fuscobrunnea* | Tr_I01 | CA | 682 | -79.8578 | 159.2881 | MK208784 | MK285376 |
| T44657 | *Lecanora fuscobrunnea* | Tr_S02 | CA | 494 | -79.8633 | 159.3733 | JN873876 | JN204804 |
| T44675 | *Lecanora fuscobrunnea* | Tr_I01 | CA | 401 | -79.8772 | 159.3264 | MK205093 | JN204806 |
| T44687 | *Lecanora fuscobrunnea* | - | CA | 486 | -79.8628 | 159.3789 | MK208799 | - |
| T44688 | *Lecanora fuscobrunnea* | Tr_S02 | CA | 486 | -79.8628 | 159.3817 | JN873877 | JN204807 |
| T44695 | *Lecanora fuscobrunnea* | - | CA | 817 | -79.7608 | 158.5033 | MK208803 | - |
| T44697 | *Lecanora fuscobrunnea* | Tr_S02 | CA | 368 | -79.7575 | 158.7575 | MK208804 | MK226932 |
| T44699 | *Lecanora fuscobrunnea* | Tr_A02 | CA | 381 | -79.7619 | 158.6356 | MK208806 | MK226934 |
| T44700 | *Lecanora fuscobrunnea* | Tr_I01 | CA | 377 | -79.7622 | 158.6369 | MK208807 | MK226935 |
| T44702 | *Lecanora fuscobrunnea* | Tr_A02 | CA | 397 | -79.7558 | 158.61 | MK208809 | MK226937 |
| T44705 | *Lecanora fuscobrunnea* | Tr_S02 | CA | 373 | -79.7556 | 158.6111 | MK208810 | MK226940 |
| T44717 | *Lecanora fuscobrunnea* | Tr_S02 | CA | 385 | -79.7553 | 158.6203 | MK208816 | MK226949 |

| | | | | | | | | |
|---|---|---|---|---|---|---|---|---|
| T44719 | *Lecanora fuscobrunnea* | Tr_S18 | CA | 1461 | -79.9236 | 156.8144 | MK208817 | MK226950 |
| T44720 | *Lecanora fuscobrunnea* | Tr_S02 | CA | 1620 | -79.9486 | 156.7889 | GU170840 | MK226951 |
| T48826 | *Lecanora fuscobrunnea* | Tr_S02 | CA | 424 | -78.0973 | 163.7167 | - | MK970694 |
| T48851 | *Lecanora fuscobrunnea* | Tr_A02 | CA | 587 | -78.0743 | 163.7931 | - | MK970698 |
| T54134 | *Lecanora fuscobrunnea* | Tr_A02 | CA | 15 | -77.0064 | 162.5328 | - | PV788659 |
| UR01200 | *Lecanora fuscobrunnea* | Tr_I01 | MA | 51 | -62.7481 | -60.3236 | PV788484 | - |
| UR01213 | *Lecanora fuscobrunnea* | Tr_I01 | MA | 17 | -62.7512 | -60.3229 | PV788485 | PV788650 |
| AAS_Smith05098 | *Lecanora physciella* | - | MA | 100 | -60.6221 | -45.5777 | - | - |
| AAS_Smith07951 | *Lecanora physciella* | Tr_A02 | MA | 400 | -64.2 | -56.85 | PV788362 | PV788516 |
| AAS_Smith07975 | *Lecanora physciella* | Tr_I01 | MA | 450 | -64.2 | -56.85 | PV788363 | PV788626 |
| AAS_Smith07976A | *Lecanora physciella* | - | MA | 450 | -64.2 | -56.85 | - | - |
| AAS_Smith09573B | *Lecanora physciella* | Tr_A02 | CA | 90 | -74.0117 | 165.2423 | PV788364 | PV788496 |
| AAS_Smith09881 | *Lecanora physciella* | - | CA | 1000 | -74.07 | 164.75 | - | - |
| AAS_Smith09921 | *Lecanora physciella* | - | CA | 100 | -72.63 | 169.37 | - | - |
| AAS_Smith09936A | *Lecanora physciella* | - | CA | 100 | -73.07 | 169.6 | - | - |
| AAS_Smith09948 | *Lecanora physciella* | - | CA | 700 | -72.52 | 169.77 | - | - |
| AAS_Smith10134B | *Lecanora physciella* | Tr_A08 | CA | 25 | -74.33 | 165.13 | PV788370 | PV788550 |
| AAS_Smith10140 | *Lecanora physciella* | Tr_S02 | CA | 650 | -74 | 164.83 | PV788371 | PV788583 |
| AAS_Smith10267 | *Lecanora physciella* | - | CA | 110 | -73.58 | 166.62 | - | - |
| AAS_Smith10301 | *Lecanora physciella* | Tr_A02 | CA | 140 | -74.0667 | 165.15 | PV788372 | PV788497 |
| AAS_Smith10314 | *Lecanora physciella* | - | MA | 0 | -72.08 | -68.55 | - | - |

| | | | | | | | | |
|---|---|---|---|---|---|---|---|---|
| AAS_Smith10521 | *Lecanora physciella* | - | MA | 85 | -69.75 | -75.25 | - | - |
| AAS_Smith10522 | *Lecanora physciella* | Tr_S02 | MA | 85 | -69.75 | -75.25 | PV788373 | PV788582 |
| AAS_Smith10554 | *Lecanora physciella* | - | MA | 125 | -69.75 | -75.25 | - | - |
| LE_A0513602 | *Lecanora physciella* | - | CA | 80 | -70.8667 | 67.95 | - | - |
| LE_A060352 | *Lecanora physciella* | - | MA | 10 | -65.261 | -63.987 | - | - |
| LE_A133911 | *Lecanora physciella* | Tr_A02 | CA | 1252 | -72.2131 | 68.60715 | - | PV788545 |
| MAF_GS1_45 | *Lecanora physciella* | Tr_S02 | CA | 162 | -84.5355 | -174.954 | MK208741 | MK226874 |
| MAF_MK1_63 | *Lecanora physciella* | Tr_S02 | CA | 859 | -83.7748 | 171.8276 | MK208766 | MK226898 |
| T33346 | *Lecanora physciella* | Tr_S18 | CA | 810 | -83.8033 | 172.2567 | JN873878 | JN204728 |
| T35704 | *Lecanora physciella* | Tr_A02 | CA | 25 | -72.3303 | 170.2614 | PV788438 | JN204754 |
| T35723 | *Lecanora physciella* | - | CA | 290 | -72.3178 | 170.2631 | - | - |
| T35769 | *Lecanora physciella* | Tr_A02 | CA | 80 | -72.3292 | 170.2231 | JN873878 | JN204755 |
| T35770 | *Lecanora physciella* | Tr_S02 | CA | 80 | -72.3292 | 170.2231 | JN873880 | JN204756 |
| T35803 | *Lecanora physciella* | Tr_A02 | CA | 70 | -72.3239 | 170.2306 | - | PV788517 |
| AAS_Smith11677 | *Lecidea andersonii* | Tr_A08 | MA | 10 | -60.6221 | -45.5777 | PV788377 | PV788551 |
| ADT17087 | *Lecidea andersonii* | | CA | NA | -66.4167 | 98.96583 | - | JN204825 |
| ADT18513 | *Lecidea andersonii* | Tr_A04 | CA | 50 | -66.2858 | 110.5436 | GU074443 | JN204827 |
| ADT19153 | *Lecidea andersonii* | - | CA | 150 | -76.9437 | 162.3333 | - | - |
| ADT24109 | *Lecidea andersonii* | Tr_A02 | CA | 25 | -66.2825 | 110.5394 | - | JN204828 |
| ADT26423 | *Lecidea andersonii* | Tr_A04 | CA | NA | -66.2653 | 110.5417 | PV788430 | JN204831 |
| LE_A042102 | *Lecidea andersonii* | Tr_A02 | CA | 110 | -70.7667 | 11.83333 | PV788314 | PV788537 |

| | | | | | | | | |
|---|---|---|---|---|---|---|---|---|
| LE_A0514501 | *Lecidea andersonii* | Tr_A02 | CA | 20 | -70.85 | 68.01667 | PV788323 | PV788525 |
| LE_A0515501 | *Lecidea andersonii* | Tr_A02 | CA | 100 | -70.7667 | 11.83333 | - | JN204815 |
| LE_A0515601 | *Lecidea andersonii* | Tr_A04 | CA | 100 | -70.7667 | 11.83333 | GU074445 | JN204816 |
| LE_A0515901 | *Lecidea andersonii* | Tr_A02 | CA | 120 | -70.7667 | 11.83333 | GU074446 | JN204817 |
| LE_A100201 | *Lecidea andersonii* | Tr_A02 | CA | 111 | -70.763 | 11.75615 | PV788324 | PV788538 |
| LE_A101501 | *Lecidea andersonii* | Tr_A02 | CA | 28 | -70.7696 | 11.85245 | PV788315 | PV788539 |
| LE_A102201 | *Lecidea andersonii* | Tr_A02 | CA | 99 | -70.7616 | 11.7871 | PV788316 | PV788540 |
| LE_A102205 | *Lecidea andersonii* | Tr_A02 | CA | 99 | -70.7616 | 11.7871 | PV788317 | PV788541 |
| LE_A102704 | *Lecidea andersonii* | Tr_A02 | CA | 100 | -70.7547 | 11.74405 | PV788318 | PV788542 |
| LE_A103003 | *Lecidea andersonii* | Tr_A13 | CA | 134 | -70.7614 | 11.58773 | PV788319 | PV788560 |
| LE_A103808 | *Lecidea andersonii* | Tr_A02 | CA | 109 | -70.7511 | 11.5309 | PV788325 | PV788522 |
| LE_A104106 | *Lecidea andersonii* | Tr_A02 | CA | 166 | -70.7637 | 11.62745 | PV788320 | PV788523 |
| LE_A104402 | *Lecidea andersonii* | Tr_A02 | CA | 159 | -70.7592 | 11.6359 | PV788321 | PV788543 |
| LE_A105701 | *Lecidea andersonii* | Tr_A02 | CA | 78 | -70.7388 | 11.43937 | MK620084 | PV788498 |
| LE_A110204 | *Lecidea andersonii* | - | CA | 30 | -67.6745 | 45.78878 | - | - |
| LE_A115504 | *Lecidea andersonii* | Tr_A02 | CA | 65 | -67.6656 | 45.89472 | PV788326 | PV788524 |
| LE_A150207 | *Lecidea andersonii* | Tr_S02 | CA | 275 | -72.1023 | 68.739 | - | - |
| LE_A151401 | *Lecidea andersonii* | Tr_A02 | CA | 1209 | -73.0472 | 65.73055 | PV788419 | - |
| MAF_GS1_12 | *Lecidea andersonii* | Tr_S02 | CA | 162 | -84.5355 | -174.954 | MK208738 | MK226871 |
| MAF_GS1_44 | *Lecidea andersonii* | Tr_A02 | CA | 162 | -84.5355 | -174.954 | MK208740 | MK226873 |
| MAF_GS1_61 | *Lecidea andersonii* | Tr_A02 | CA | 162 | -84.5355 | -174.954 | MK208744 | MK226877 |

| | | | | | | | | |
|---|---|---|---|---|---|---|---|---|
| MAF_MG1_16 | *Lecidea andersonii* | Tr_A02 | CA | 230 | -84.5589 | -175.009 | MK208748 | MK226881 |
| MAF_MG1_17 | *Lecidea andersonii* | Tr_A02 | CA | 230 | -84.5589 | -175.009 | MK208749 | MK226882 |
| MAF_MG1_23 | *Lecidea andersonii* | Tr_A02 | CA | 230 | -84.5589 | -175.009 | MK208750 | PV788529 |
| T33001 | *Lecidea andersonii* | Tr_A02 | CA | 40 | -77.0167 | 162.5333 | PV788327 | JN204727 |
| T33002 | *Lecidea andersonii* | - | CA | 40 | -77.0167 | 162.5333 | - | - |
| T33456 | *Lecidea andersonii* | Tr_A02 | CA | 530 | -83.7606 | 172.755 | PV788328 | PV788530 |
| T33696 | *Lecidea andersonii* | Tr_A02 | CA | 60 | -77.6004 | 163.1509 | - | - |
| T35636 | *Lecidea andersonii* | Tr_A02 | CA | 410 | -79.84 | 159.3486 | - | - |
| T35888 | *Lecidea andersonii* | Tr_A02 | CA | 15 | -77.0071 | 162.6298 | PV788341 | PV788526 |
| T35920 | *Lecidea andersonii* | Tr_A12 | CA | 15 | -77.0071 | 162.6298 | PV788329 | PV788553 |
| T35959 | *Lecidea andersonii* | Tr_A02 | CA | 15 | -77.0071 | 162.6298 | PV788342 | PV788499 |
| T42995 | *Lecidea andersonii* | Tr_A02 | CA | 730 | -77.0362 | 161.6032 | GU074447 | JN204772 |
| T42996 | *Lecidea andersonii* | Tr_A02 | CA | 734 | -77.0344 | 161.7308 | GU074448 | PV788500 |
| T42997 | *Lecidea andersonii* | Tr_A02 | CA | 733 | -77.0343 | 161.7318 | PV788330 | JN204773 |
| T42998 | *Lecidea andersonii* | Tr_A02 | CA | 758 | -77.0333 | 161.7325 | PV788343 | PV788501 |
| T43001 | *Lecidea andersonii* | Tr_A02 | CA | 525 | -77.0865 | 161.7238 | PV788331 | PV788527 |
| T43002 | *Lecidea andersonii* | Tr_A02 | CA | 525 | -77.0866 | 161.7233 | GU074449 | PV788528 |
| T43003 | *Lecidea andersonii* | Tr_A12 | CA | 525 | -77.0522 | 161.7236 | GU074451 | JN204775 |
| T43006 | *Lecidea andersonii* | Tr_A02 | CA | 48 | -77.0069 | 162.5325 | GU074453 | JN204777 |
| T43008 | *Lecidea andersonii* | Tr_I01 | CA | 20 | -77.006 | 162.5585 | GU074450 | JN204778 |
| T43009 | *Lecidea andersonii* | Tr_A02 | CA | 19 | -77.0059 | 162.5509 | PV788332 | PV788502 |

| | | | | | | | | |
|---|---|---|---|---|---|---|---|---|
| T43010 | *Lecidea andersonii* | Tr_A12 | CA | 20 | -77.0061 | 162.5586 | PV788333 | PV788554 |
| T43011 | *Lecidea andersonii* | Tr_A02 | CA | 20 | -77.0061 | 162.5561 | GU074452 | JN204779 |
| T43012 | *Lecidea andersonii* | Tr_I01 | CA | 55 | -77.0068 | 162.5603 | PV788334 | PV788639 |
| T43018 | *Lecidea andersonii* | - | CA | 8 | -77.0078 | 162.5188 | - | - |
| T43021 | *Lecidea andersonii* | Tr_A02 | CA | 9 | -77.0074 | 162.5206 | PV788335 | PV788515 |
| T43022 | *Lecidea andersonii* | Tr_A02 | CA | 40 | -77.0144 | 162.5809 | GU074454 | JN204785 |
| T43024 | *Lecidea andersonii* | Tr_A02 | CA | 17 | -77.0119 | 162.4976 | PV788336 | PV788518 |
| T43025 | *Lecidea andersonii* | Tr_A02 | CA | 16 | -77.0175 | 162.4648 | PV788337 | PV788503 |
| T43026 | *Lecidea andersonii* | Tr_A02 | CA | 19 | -77.0131 | 162.4886 | PV788338 | JN204786 |
| T43027 | *Lecidea andersonii* | Tr_I01 | CA | 15 | -77.0103 | 162.5066 | PV788339 | PV788627 |
| T43090 | *Lecidea andersonii* | - | CA | 40 | -77.8486 | 166.7431 | - | - |
| T48807 | *Lecidea andersonii* | Tr_A02 | CA | 554 | -78.1409 | 163.6307 | MK970673 | MK970702 |
| UR00249 | *Lecidea andersonii* | - | CA | 829 | -76.9167 | 160.9167 | PV788340 | - |
| AAS_Convey00458 | *Lecidea atrobrunnea* | Tr_I01 | MA | 50 | -68.838 | -67.3476 | MK620076 | PV788644 |
| AAS_Smith12030 | *Lecidea atrobrunnea* | Tr_I01 | MA | 120 | -64.83 | -62.52 | MK620079 | PV788643 |
| AAS_Smith12031 | *Lecidea atrobrunnea* | Tr_I01 | MA | 120 | -64.83 | -62.52 | PV788378 | PV788646 |
| AAS_Smith12032 | *Lecidea atrobrunnea* | - | MA | 120 | -64.83 | -62.52 | - | - |
| LE_A060124 | *Lecidea atrobrunnea* | Tr_I01 | MA | 50 | -64.8972 | -62.871 | GU074457 | JN204819 |
| LE_A060137 | *Lecidea atrobrunnea* | Tr_I01 | MA | 50 | -64.8972 | -62.871 | MK620083 | PV788647 |
| LE_A060206 | *Lecidea atrobrunnea* | Tr_I01 | MA | 10 | -64.8283 | -63.4917 | PV788304 | JN204820 |
| LE_A060330 | *Lecidea atrobrunnea* | Tr_A02 | MA | 40 | -65.2614 | -63.9874 | PV788305 | JN204821 |

| | | | | | | | | |
|---|---|---|---|---|---|---|---|---|
| LE_A060414 | *Lecidea atrobrunnea* | Tr_I01 | MA | 30 | -65.1 | -64.05 | PV788313 | JN204822 |
| T36121 | *Lecidea atrobrunnea* | Tr_I01 | MA | 10 | -62.6717 | -61.1047 | GU074455 | JN204759 |
| UR00878 | *Lecidea atrobrunnea* | Tr_I01 | MA | 40 | -62.6618 | -60.3787 | PV788308 | PV788653 |
| UR00932 | *Lecidea atrobrunnea* | - | MA | 383 | -62.6816 | -60.3422 | PV788309 | - |
| UR00933 | *Lecidea atrobrunnea* | Tr_S02 | MA | 383 | -62.6816 | -60.3422 | PV788310 | PV788596 |
| UR00979 | *Lecidea atrobrunnea* | Tr_I01 | MA | 46 | -62.6432 | -60.3726 | PV788311 | PV788654 |
| UR00981 | *Lecidea atrobrunnea* | Tr_I01 | MA | 46 | -62.6432 | -60.3726 | PV788312 | PV788655 |
| UR01203 | *Lecidea atrobrunnea* | Tr_I01 | MA | 51 | -62.7481 | -60.3236 | PV788306 | PV788565 |
| UR01212 | *Lecidea atrobrunnea* | Tr_I01 | MA | 51 | -62.7481 | -60.3236 | PV788307 | PV788656 |
| AAS_Convey00489 | *Lecidea cancriformis* | Tr_S02 | CA | 1323 | -75.8333 | -69.4354 | PV788303 | PV788572 |
| AAS_Convey00490 | *Lecidea cancriformis* | - | CA | 1323 | -75.83 | -69.37 | - | - |
| AAS_Convey00491 | *Lecidea cancriformis* | Tr_S02 | CA | 1323 | -75.8407 | -69.4201 | PV788355 | PV788599 |
| AAS_Convey00509B | *Lecidea cancriformis* | - | CA | 860 | -75.83 | -69.37 | - | - |
| AAS_Convey00551 | *Lecidea cancriformis* | - | CA | 791 | -75.43 | -72.6 | - | - |
| AAS_Convey00553 | *Lecidea cancriformis* | - | CA | 791 | -75.43 | -72.6 | - | - |
| AAS_Convey00572A | *Lecidea cancriformis* | - | CA | 791 | -75.43 | -72.6 | - | - |
| AAS_Convey00604 | *Lecidea cancriformis* | - | CA | 1167 | -77.03 | -78.27 | - | - |
| AAS_Convey00605 | *Lecidea cancriformis* | Tr_S02 | CA | 1167 | -77.03 | -78.27 | PV788356 | PV788601 |
| AAS_Convey00606 | *Lecidea cancriformis* | - | CA | 1167 | -77.03 | -78.27 | - | - |
| AAS_Convey00608 | *Lecidea cancriformis* | - | CA | 1167 | -77.03 | -78.27 | - | - |
| AAS_Convey00609 | *Lecidea cancriformis* | - | CA | 1167 | -77.03 | -78.27 | - | - |

| | | | | | | | | |
|---|---|---|---|---|---|---|---|---|
| AAS_Convey00672 | *Lecidea cancriformis* | Tr_S02 | CA | 1760 | -66.78 | -64.88 | PV788386 | PV788584 |
| AAS_Convey00801 | *Lecidea cancriformis* | Tr_S02 | CA | NA | -80.52 | -20.42 | PV788357 | PV788605 |
| AAS_Convey01112 | *Lecidea cancriformis* | Tr_I01 | CA | 1395 | -79.85 | -82.92 | PV788358 | PV788630 |
| AAS_FFE00009 | *Lecidea cancriformis* | Tr_S02 | CA | 908 | -79.824 | -83.051 | PV788359 | PV788585 |
| AAS_FFE00012 | *Lecidea cancriformis* | Tr_S18 | CA | 908 | -79.824 | -83.051 | MK620078 | PV788606 |
| AAS_FFE00014 | *Lecidea cancriformis* | Tr_S18 | CA | 908 | -79.824 | -83.051 | PV788360 | PV788607 |
| AAS_FFE00017 | *Lecidea cancriformis* | Tr_S18 | CA | 908 | -79.824 | -83.051 | PV788361 | PV788622 |
| AAS_Smith07039A | *Lecidea cancriformis* | - | MA | 150 | -60.6221 | -45.5777 | - | - |
| AAS_Smith09771 | *Lecidea cancriformis* | Tr_A02 | CA | 20 | -74.33 | 165.13 | PV788365 | PV788504 |
| AAS_Smith09782 | *Lecidea cancriformis* | Tr_S02 | CA | 100 | -73.3038 | 167.4247 | PV788366 | PV788579 |
| AAS_Smith09787 | *Lecidea cancriformis* | - | CA | 100 | -73.35 | 167.92 | - | - |
| AAS_Smith09982 | *Lecidea cancriformis* | Tr_I01 | CA | 200 | -74.83 | 162.55 | PV788367 | - |
| AAS_Smith10057B | *Lecidea cancriformis* | Tr_I01 | CA | 1494 | -86.48 | -145.93 | PV788368 | PV788645 |
| AAS_Smith10603B | *Lecidea cancriformis* | - | MA | 140 | -67.55 | -68.18 | - | - |
| AAS_Smith11341 | *Lecidea cancriformis* | Tr_I01 | MA | 25 | -63 | -60.58 | PV788374 | PV788657 |
| AAS_Smith11544 | *Lecidea cancriformis* | Tr_S02 | MA | 450 | -70.85 | -68.58 | PV788375 | PV788578 |
| AAS_Smith11568 | *Lecidea cancriformis* | - | MA | 500 | -70.9 | -68.48 | - | - |
| AAS_Smith11604 | *Lecidea cancriformis* | Tr_I01 | MA | 582 | -70.82 | -68.42 | PV788376 | PV788658 |
| ADT_RS18317 | *Lecidea cancriformis* | | CA | NA | -66.2833 | 110.5167 | - | JN204840 |
| ADT_RS19162 | *Lecidea cancriformis* | | CA | NA | -76.9333 | 162.3333 | - | JN204841 |
| ADT_RSN2_1 | *Lecidea cancriformis* | | CA | NA | -67.8 | 66.7 | - | JN204842 |

| | | | | | | | | |
|---|---|---|---|---|---|---|---|---|
| ADT_RSRL | *Lecidea cancriformis* | | CA | NA | -69.4083 | 76.4 | - | JN204843 |
| ADT24118 | *Lecidea cancriformis* | | CA | NA | -66.2825 | 110.5394 | - | JN204829 |
| ADT25151 | *Lecidea cancriformis* | Tr_A02 | CA | 151 | -72.3604 | 169.9096 | GU074429 | JN204830 |
| ADT5902 | *Lecidea cancriformis* | | CA | NA | -66.5818 | 110.6922 | - | JN204832 |
| LE_A040202 | *Lecidea cancriformis* | Tr_A13 | CA | 50 | -69.4296 | 76.38333 | - | PV788564 |
| LE_A041102 | *Lecidea cancriformis* | - | CA | 70 | -69.7489 | 73.68248 | - | - |
| LE_A042607 | *Lecidea cancriformis* | Tr_A13 | CA | 120 | -70.7667 | 11.83333 | PV788422 | PV788561 |
| LE_A045001 | *Lecidea cancriformis* | Tr_I01 | CA | 90 | -69.4083 | 76.4 | PV788410 | JN204813 |
| LE_A046401 | *Lecidea cancriformis* | Tr_A12 | CA | 70 | -69.4064 | 76.36014 | PV788379 | PV788556 |
| LE_A0511101 | *Lecidea cancriformis* | - | CA | 40 | -70.8333 | 68.25 | GU074440 | - |
| LE_A0513401 | *Lecidea cancriformis* | Tr_I01 | CA | 80 | -70.8667 | 67.95 | - | |
| LE_A0514005 | *Lecidea cancriformis* | - | CA | 350 | -70.9 | 67.91667 | - | - |
| LE_A057501 | *Lecidea cancriformis* | Tr_A02 | CA | 30 | -69.4083 | 76.4 | - | JN204818 |
| LE_A100101 | *Lecidea cancriformis* | Tr_I01 | CA | 142 | -70.7724 | 11.75837 | PV788411 | PV788628 |
| LE_A105803 | *Lecidea cancriformis* | Tr_I01 | CA | 110 | -70.762 | 11.74723 | PV788412 | PV788640 |
| LE_A105804 | *Lecidea cancriformis* | Tr_I01 | CA | 110 | -70.762 | 11.74723 | PV788413 | PV788629 |
| LE_A110804 | *Lecidea cancriformis* | Tr_I01 | CA | 57 | -67.6802 | 45.83198 | PV788380 | PV788634 |
| LE_A111006 | *Lecidea cancriformis* | Tr_I01 | CA | 55 | -67.6805 | 45.82288 | PV788423 | PV788635 |
| LE_A111717 | *Lecidea cancriformis* | Tr_S02 | CA | 32 | -67.661 | 45.84795 | PV788381 | PV788570 |
| LE_A119001 | *Lecidea cancriformis* | Tr_S02 | CA | 98 | -67.6754 | 45.8816 | PV788382 | PV788571 |
| LE_A11no | *Lecidea cancriformis* | Tr_S02 | CA | 758 | -70.9382 | 67.80742 | PV788387 | LE_A11no |

| | | | | | | | | |
|---|---|---|---|---|---|---|---|---|
| LE_A130405 | *Lecidea cancriformis* | Tr_I01 | CA | 129 | -72.2235 | 68.80912 | PV788388 | PV788637 |
| LE_A131006 | *Lecidea cancriformis* | Tr_S02 | CA | 127 | -72.218 | 68.81952 | PV788395 | PV788593 |
| LE_A131302 | *Lecidea cancriformis* | Tr_A13 | CA | 124 | -72.2142 | 68.82712 | PV788414 | PV788562 |
| LE_A131401 | *Lecidea cancriformis* | Tr_I01 | CA | 166 | -72.2281 | 68.7841 | PV788415 | PV788638 |
| LE_A132205 | *Lecidea cancriformis* | Tr_I01 | CA | 89 | -72.2199 | 68.8046 | PV788416 | PV788651 |
| LE_A133105 | *Lecidea cancriformis* | Tr_I01 | CA | 205 | -72.2184 | 68.79777 | PV788417 | PV788652 |
| LE_A150307 | *Lecidea cancriformis* | Tr_S02 | CA | 265 | -72.1022 | 68.73928 | PV788418 | - |
| LE_A151501 | *Lecidea cancriformis* | Tr_S02 | CA | 1252 | -73.0459 | 65.73447 | MK620087 | PV788591 |
| LE_A151701 | *Lecidea cancriformis* | Tr_S18 | CA | 711 | -73.1129 | 66.20897 | PV788389 | PV788608 |
| LE_A151901 | *Lecidea cancriformis* | Tr_I01 | CA | 680 | -73.1097 | 66.21585 | PV788383 | PV788636 |
| LE_A151902 | *Lecidea cancriformis* | Tr_S18 | CA | 680 | -73.1097 | 66.21585 | MK620088 | PV788609 |
| LE_A152002 | *Lecidea cancriformis* | Tr_S18 | CA | 692 | -73.1061 | 66.21518 | PV788384 | PV788610 |
| LE_A152101 | *Lecidea cancriformis* | Tr_S18 | CA | 687 | -73.104 | 66.22078 | PV788390 | PV788611 |
| LE_A152601 | *Lecidea cancriformis* | Tr_S18 | CA | 525 | -73.0907 | 66.43533 | PV788391 | PV788620 |
| LE_A153501 | *Lecidea cancriformis* | Tr_S18 | CA | 751 | -72.9423 | 65.6961 | PV788393 | PV788621 |
| LE_A153601 | *Lecidea cancriformis* | Tr_S18 | CA | 751 | -72.938 | 65.70153 | PV788394 | PV788612 |
| LE_A153801 | *Lecidea cancriformis* | Tr_A02 | CA | 774 | -72.9355 | 65.6404 | PV788421 | PV788534 |
| LE_A154701 | *Lecidea cancriformis* | Tr_A02 | CA | 526 | -73.01 | 66.46623 | PV788344 | PV788535 |
| LE_A890510 | *Lecidea cancriformis* | Tr_A02 | CA | 70 | -66.5448 | 93.00279 | PV788396 | PV788505 |
| LE_A893902 | *Lecidea cancriformis* | Tr_S02 | CA | 144 | -66.3 | 100.75 | PV788385 | PV788589 |
| LE_Ru21_3 | *Lecidea cancriformis* | Tr_A12 | CA | 142 | -74.7732 | -136.792 | PV788322 | PV788555 |

| | | | | | | | | |
|---|---|---|---|---|---|---|---|---|
| MAF_DP1_01 | *Lecidea cancriformis* | Tr_S02 | CA | 370 | -85.5392 | -151.15 | MK208709 | MK226844 |
| MAF_DP1_04 | *Lecidea cancriformis* | Tr_S02 | CA | 370 | -85.5392 | -151.15 | MK208711 | MK226846 |
| MAF_DP1_06 | *Lecidea cancriformis* | Tr_S02 | CA | 370 | -85.5392 | -151.15 | MK208712 | PV788575 |
| MAF_DP1_09 | *Lecidea cancriformis* | Tr_S02 | CA | 370 | -85.5392 | -151.15 | MK208715 | MK226849 |
| MAF_DP1_10 | *Lecidea cancriformis* | Tr_S02 | CA | 370 | -85.5392 | -151.15 | MK208716 | MK226850 |
| MAF_DP1_11 | *Lecidea cancriformis* | Tr_S02 | CA | 370 | -85.5392 | -151.15 | MK208717 | MK226851 |
| MAF_DP1_19 | *Lecidea cancriformis* | - | CA | 370 | -85.5392 | -151.15 | MK208719 | - |
| MAF_DP1_24 | *Lecidea cancriformis* | Tr_I01 | CA | 370 | -85.5392 | -151.15 | MK208722 | MK226855 |
| MAF_DP1_25 | *Lecidea cancriformis* | Tr_A02 | CA | 370 | -85.5392 | -151.15 | MK208723 | MK226856 |
| MAF_DP1_30 | *Lecidea cancriformis* | Tr_S02 | CA | 370 | -85.5392 | -151.15 | MK208725 | MK226858 |
| MAF_DP1_33 | *Lecidea cancriformis* | Tr_S02 | CA | 370 | -85.5392 | -151.15 | MK208727 | MK226860 |
| MAF_DP1_34 | *Lecidea cancriformis* | Tr_S02 | CA | 370 | -85.5392 | -151.15 | MK208728 | MK226861 |
| MAF_DP1_36 | *Lecidea cancriformis* | Tr_S02 | CA | 370 | -85.5392 | -151.15 | MK208730 | MK226863 |
| MAF_DP1_39 | *Lecidea cancriformis* | Tr_S02 | CA | 370 | -85.5392 | -151.15 | MK208731 | PV788587 |
| MAF_DP1_42 | *Lecidea cancriformis* | Tr_A02 | CA | 370 | -85.5392 | -151.15 | - | - |
| MAF_GR1_29 | *Lecidea cancriformis* | - | CA | 151 | -83.4868 | 170.7899 | MK208737 | - |
| MAF_MG1_47 | *Lecidea cancriformis* | Tr_I01 | CA | 230 | -84.5589 | -175.009 | MK208751 | MK226883 |
| MAF_MK1_05 | *Lecidea cancriformis* | Tr_S02 | CA | 859 | -83.7748 | 171.8276 | - | - |
| MAF_MK1_26 | *Lecidea cancriformis* | Tr_S02 | CA | 859 | -83.7748 | 171.8276 | MK208755 | MK226888 |
| MAF_MK1_31 | *Lecidea cancriformis* | Tr_I01 | CA | 859 | -83.7748 | 171.8276 | MK208757 | MK226890 |
| MAF_MK1_37 | *Lecidea cancriformis* | Tr_I01 | CA | 859 | -83.7748 | 171.8276 | MK208758 | MK226891 |

| | | | | | | | | |
|---|---|---|---|---|---|---|---|---|
| MAF_MK1_38 | *Lecidea cancriformis* | Tr_S02 | CA | 859 | -83.7748 | 171.8276 | MK208759 | MK226892 |
| MAF_MK1_48 | *Lecidea cancriformis* | Tr_S02 | CA | 859 | -83.7748 | 171.8276 | MK208762 | MK226895 |
| MAF_MK1_55 | *Lecidea cancriformis* | - | CA | 859 | -83.7748 | 171.8276 | MK208764 | - |
| MAF_Sancho2 | *Lecidea cancriformis* | Tr_A04 | CA | NA | -83.7606 | 172.755 | GU074439 | JN204838 |
| T33348 | *Lecidea cancriformis* | - | CA | 810 | -83.8033 | 172.2067 | PV788397 | - |
| T33446 | *Lecidea cancriformis* | Tr_S18 | CA | 490 | -83.8275 | 172.7489 | PV788398 | MK226901 |
| T33599 | *Lecidea cancriformis* | Tr_A02 | CA | 604 | -77.5739 | 163.1336 | PV788407 | JN204732 |
| T33711 | *Lecidea cancriformis* | Tr_S02 | CA | 1143 | -77.6964 | 162.2599 | PV788404 | JN204743 |
| T33712 | *Lecidea cancriformis* | Tr_S02 | CA | 1143 | -77.6964 | 162.2599 | EU257677 | JN204744 |
| T33717 | *Lecidea cancriformis* | Tr_S18 | CA | 1180 | -77.6964 | 162.2599 | PV788406 | PV788613 |
| T33718 | *Lecidea cancriformis* | Tr_S18 | CA | 1180 | -77.6964 | 162.2599 | PV788420 | PV788614 |
| T35533 | *Lecidea cancriformis* | - | CA | 402 | -79.8383 | 159.3322 | - | - |
| T35534 | *Lecidea cancriformis* | Tr_A02 | CA | 402 | -79.8383 | 159.3322 | PV788408 | PV788532 |
| T35543 | *Lecidea cancriformis* | - | CA | 421 | -79.8381 | 159.3408 | - | - |
| T35559 | *Lecidea cancriformis* | Tr_S18 | CA | 402 | -79.8383 | 159.2208 | MK208771 | PV788617 |
| T35604 | *Lecidea cancriformis* | Tr_S18 | CA | 260 | -79.8356 | 159.3167 | EU257671 | JN204749 |
| T35620 | *Lecidea cancriformis* | Tr_I01 | CA | 260 | -79.835 | 159.385 | EU257672 | JN204750 |
| T35622 | *Lecidea cancriformis* | Tr_I01 | CA | 345 | -79.835 | 159.3922 | MK208772 | PV788631 |
| T35634 | *Lecidea cancriformis* | - | CA | 410 | -79.84 | 159.3486 | - | - |
| T35635 | *Lecidea cancriformis* | - | CA | 410 | -79.84 | 159.3486 | - | - |
| T35662 | *Lecidea cancriformis* | Tr_I01 | CA | 361 | -79.84 | 159.3322 | EU257673 | JN204753 |

| | | | | | | | | |
|---|---|---|---|---|---|---|---|---|
| T42988 | *Lecidea cancriformis* | Tr_S18 | CA | 1112 | -79.8894 | 156.7636 | GU074435 | MK226902 |
| T42990 | *Lecidea cancriformis* | Tr_S18 | CA | 963 | -79.9261 | 156.89 | GU170841 | JN204770 |
| T42991 | *Lecidea cancriformis* | Tr_S18 | CA | 935 | -79.9169 | 156.7586 | GU170842 | MK226903 |
| T42992 | *Lecidea cancriformis* | Tr_S18 | CA | 935 | -79.9169 | 156.7506 | GU074436 | JN204771 |
| T43000 | *Lecidea cancriformis* | Tr_S18 | CA | 741 | -77.0339 | 161.7292 | GU074433 | PV788615 |
| T43005 | *Lecidea cancriformis* | - | CA | 12 | -77.006 | 162.5318 | MK620096 | - |
| T43016 | *Lecidea cancriformis* | Tr_A02 | CA | 8 | -77.0073 | 162.5215 | PV788405 | JN204783 |
| T43019 | *Lecidea cancriformis* | Tr_A02 | CA | 9 | -77.0078 | 162.5188 | PV788409 | PV788519 |
| T43020 | *Lecidea cancriformis* | Tr_A02 | CA | 397 | -77.0104 | 162.6054 | GU074431 | PV788520 |
| T43023 | *Lecidea cancriformis* | Tr_S18 | CA | 397 | -77.0129 | 162.5945 | GU074432 | PV788616 |
| T43028 | *Lecidea cancriformis* | Tr_A02 | CA | 12 | -77.0068 | 162.5285 | GU074430 | PV788531 |
| T44625 | *Lecidea cancriformis* | Tr_A02 | CA | 492 | -79.8683 | 159.3603 | MK208773 | MK226905 |
| T44633 | *Lecidea cancriformis* | Tr_S02 | CA | 490 | -79.8683 | 159.3583 | MK208775 | PV788588 |
| T44634 | *Lecidea cancriformis* | Tr_S18 | CA | 533 | -79.8686 | 159.3406 | GU074434 | JN204801 |
| T44638 | *Lecidea cancriformis* | Tr_S18 | CA | 517 | -79.8686 | 159.3575 | PV788399 | MK226907 |
| T44640 | *Lecidea cancriformis* | Tr_S02 | CA | 517 | -79.8686 | 159.3575 | MK208777 | MK226908 |
| T44643 | *Lecidea cancriformis* | Tr_S18 | CA | 1114 | -79.8594 | 159.2383 | MK208778 | PV788618 |
| T44645 | *Lecidea cancriformis* | Tr_S02 | CA | 746 | -79.8567 | 159.2831 | MK208779 | MK226909 |
| T44646 | *Lecidea cancriformis* | Tr_S02 | CA | 746 | -79.8567 | 159.2831 | MK208780 | MK226910 |
| T44647 | *Lecidea cancriformis* | Tr_S02 | CA | 682 | -79.8578 | 159.2881 | MK208781 | MK226911 |
| T44648 | *Lecidea cancriformis* | - | CA | 682 | -79.8578 | 159.2881 | MK208782 | - |

| | | | | | | | | |
|---|---|---|---|---|---|---|---|---|
| T44649 | *Lecidea cancriformis* | Tr_S02 | CA | 682 | -79.8578 | 159.2881 | MK208783 | MK226912 |
| T44651 | *Lecidea cancriformis* | Tr_S02 | CA | 682 | -79.8578 | 159.2881 | MK208785 | MK226913 |
| T44652 | *Lecidea cancriformis* | Tr_S02 | CA | 682 | -79.8578 | 159.2881 | MK208786 | MK226914 |
| T44656 | *Lecidea cancriformis* | Tr_S02 | CA | 436 | -79.8633 | 159.3733 | MK208788 | MK226916 |
| T44659 | *Lecidea cancriformis* | Tr_S02 | CA | 517 | -79.8642 | 159.3678 | MK208789 | MK226917 |
| T44665 | *Lecidea cancriformis* | Tr_S02 | CA | 534 | -79.8689 | 159.3447 | MK208790 | MK226918 |
| T44666 | *Lecidea cancriformis* | - | CA | 534 | -79.8689 | 159.3447 | MK2087917 | - |
| T44667 | *Lecidea cancriformis* | Tr_S02 | CA | 534 | -79.8689 | 159.3447 | MK208792 | MK226919 |
| T44669 | *Lecidea cancriformis* | Tr_S02 | CA | 487 | -79.8628 | 159.3775 | MK208793 | MK226920 |
| T44670 | *Lecidea cancriformis* | Tr_I01 | CA | 487 | -79.8628 | 159.3775 | MK208794 | MK226921 |
| T44673 | *Lecidea cancriformis* | Tr_S02 | CA | 401 | -79.8772 | 159.3264 | - | - |
| T44674 | *Lecidea cancriformis* | Tr_S02 | CA | 401 | -79.8772 | 159.3264 | MK208795 | MK226923 |
| T44676 | *Lecidea cancriformis* | Tr_S18 | CA | 146 | -79.8831 | 159.3483 | MK208796 | MK226924 |
| T44677 | *Lecidea cancriformis* | Tr_A02 | CA | 146 | -79.8831 | 159.3483 | MK208797 | MK226925 |
| T44678 | *Lecidea cancriformis* | Tr_A02 | CA | 146 | -79.8831 | 159.3483 | - | - |
| T44679 | *Lecidea cancriformis* | Tr_S02 | CA | 146 | -79.8772 | 159.3314 | MK208798 | MK226927 |
| T44686 | *Lecidea cancriformis* | Tr_S02 | CA | 486 | -79.8628 | 159.3789 | - | - |
| T44689 | *Lecidea cancriformis* | Tr_S18 | CA | 486 | -79.865 | 159.37 | - | - |
| T44690 | *Lecidea cancriformis* | Tr_S02 | CA | 486 | -79.8628 | 159.3817 | MK208800 | MK226930 |
| T44691 | *Lecidea cancriformis* | Tr_S18 | CA | 486 | -79.8628 | 159.3817 | - | JN204808 |
| T44692 | *Lecidea cancriformis* | Tr_S02 | CA | 491 | -79.8656 | 159.3656 | MK208801 | JN204809 |

| | | | | | | | | |
|---|---|---|---|---|---|---|---|---|
| T44694 | *Lecidea cancriformis* | Tr_I01 | CA | 815 | -79.7553 | 158.5033 | MK208802 | MK226931 |
| T44698 | *Lecidea cancriformis* | Tr_S18 | CA | 817 | -79.755 | 158.6267 | MK208805 | MK226933 |
| T44701 | *Lecidea cancriformis* | Tr_S18 | CA | 363 | -79.7569 | 158.6075 | MK208808 | MK226936 |
| T44703 | *Lecidea cancriformis* | Tr_S18 | CA | 395 | -79.7561 | 158.6144 | PV788400 | MK226938 |
| T44704 | *Lecidea cancriformis* | Tr_S18 | CA | 396 | -79.7558 | 158.6172 | PV788401 | MK226939 |
| T44707 | *Lecidea cancriformis* | Tr_S18 | CA | 365 | -79.7578 | 158.5981 | MK208811 | MK226941 |
| T44708 | *Lecidea cancriformis* | Tr_S18 | CA | 464 | -79.7525 | 158.5489 | MK208812 | PV788619 |
| T44709 | *Lecidea cancriformis* | Tr_S18 | CA | 493 | -79.7525 | 158.5406 | MK208813 | MK226942 |
| T44710 | *Lecidea cancriformis* | Tr_S18 | CA | 622 | -79.7553 | 158.5211 | - | - |
| T44711 | *Lecidea cancriformis* | Tr_S18 | CA | 642 | -79.7553 | 158.5167 | - | - |
| T44712 | *Lecidea cancriformis* | Tr_S18 | CA | 749 | -79.7594 | 158.5069 | GU074438 | JN204811 |
| T44713 | *Lecidea cancriformis* | Tr_S18 | CA | 718 | -79.7586 | 158.5111 | MK208814 | MK226945 |
| T44714 | *Lecidea cancriformis* | Tr_S18 | CA | 904 | -79.7625 | 158.4969 | MK208815 | MK226946 |
| T44715 | *Lecidea cancriformis* | Tr_S18 | CA | 380 | -79.7578 | 158.6022 | PV788402 | MK226947 |
| T44716 | *Lecidea cancriformis* | Tr_S18 | CA | 371 | -79.7578 | 158.6058 | PV788403 | MK226948 |
| T44721 | *Lecidea cancriformis* | Tr_S18 | CA | 434 | -79.7539 | 158.6358 | MK208818 | MK226952 |
| T44723 | *Lecidea cancriformis* | Tr_S18 | CA | 927 | -79.8781 | 157.5266 | MK208819 | MK226953 |
| T44727 | *Lecidea cancriformis* | Tr_S18 | CA | 1620 | -79.892 | 157.5238 | GU074437 | JN204812 |
| T44787 | *Lecidea cancriformis* | Tr_S18 | CA | 1280 | -79.9293 | 156.7048 | MK208820 | MK226955 |
| T46677 | *Lecidea cancriformis* | Tr_A02 | CA | 843 | -78.0189 | 163.9706 | MK970681 | MK970696 |
| T46684 | *Lecidea cancriformis* | Tr_S02 | CA | 788 | -78.0344 | 163.9636 | MK970677 | MK970693 |

| | | | | | | | | |
|---|---|---|---|---|---|---|---|---|
| T46685 | *Lecidea cancriformis* | Tr_S02 | CA | 870 | -78.044 | 163.986 | MK970677 | MK970693 |
| T48776 | *Lecidea cancriformis* | Tr_A02 | CA | 783 | -78.165 | 163.7532 | MK970679 | MK970702 |
| T48777 | *Lecidea cancriformis* | Tr_S02 | CA | 766 | -78.1655 | 163.7545 | MK970679 | MK970692 |
| T48782 | *Lecidea cancriformis* | Tr_S18 | CA | 679 | -78.1228 | 163.6417 | MK970679 | MK970695 |
| T48793a | *Lecidea cancriformis* | Tr_S02 | CA | 831 | -78.1529 | 163.7312 | MK970677 | MK970692 |
| T48799b | *Lecidea cancriformis* | Tr_A02 | CA | 743 | -78.1453 | 163.6203 | MK970679 | MK970698 |
| T48799c | *Lecidea cancriformis* | Tr_S02 | CA | 743 | -78.1453 | 163.6203 | MK970677 | MK970692 |
| T48843a | *Lecidea cancriformis* | Tr_S02 | CA | 836 | -78.0659 | 163.8699 | MK970677 | MK970693 |
| T48843c | *Lecidea cancriformis* | - | CA | 836 | -78.0659 | 163.8699 | MK970678 | - |
| T48855b | *Lecidea cancriformis* | Tr_S02 | CA | 688 | -78.0362 | 163.8365 | MK970680 | MK970692 |
| T48862 | *Lecidea cancriformis* | - | CA | 875 | -78.0374 | 163.9782 | MK970682 | - |
| T48864 | *Lecidea cancriformis* | Tr_A02 | CA | 850 | -78.0393 | 163.9893 | MK970679 | MK970702 |
| T48887a | *Lecidea cancriformis* | Tr_S02 | CA | 710 | -78.0699 | 163.7112 | MK970679 | MK970692 |
| UR00906 | *Lecidea cancriformis* | - | MA | 142 | -62.6666 | -60.392 | PV788424 | - |
| UR00922 | *Lecidea cancriformis* | - | MA | 383 | -62.6816 | -60.3422 | PV788425 | - |
| LE_A890206 | *Lecidea lapicida* | - | CA | 20 | -67.9623 | 44.58029 | - | - |
| LE_A893504 | *Lecidea lapicida* | Tr_I01 | CA | 144 | -66.3 | 100.75 | PV788345 | PV788632 |
| LE_A894101 | *Lecidea lapicida* | Tr_S02 | CA | 144 | -66.3 | 100.75 | PV788346 | PV788590 |
| MAF_LI_2f | *Lecidea lapicida* | Tr_A02 | CA | 15 | -62.6618 | -60.3817 | MK620093 | PV788506 |
| MAF_LI_3b | *Lecidea lapicida* | Tr_A02 | CA | 15 | -62.6618 | -60.3817 | MK620094 | PV788507 |
| T48875 | *Lecidea lapicida* | Tr_A02 | CA | 375 | -78.0239 | 163.8999 | MK620098 | MK970699 |

| | | | | | | | | |
|---|---|---|---|---|---|---|---|---|
| UR00851 | *Lecidea lapicida* | Tr_S02 | MA | 29 | -62.6643 | -60.3898 | PV788349 | PV788597 |
| UR00892 | *Lecidea lapicida* | - | MA | 271 | -62.6643 | -60.3933 | PV788350 | - |
| UR00903 | *Lecidea lapicida* | - | MA | 142 | -62.6666 | -60.392 | PV788351 | - |
| UR00907 | *Lecidea lapicida* | - | MA | 123 | -62.6677 | -60.392 | PV788353 | - |
| UR00974 | *Lecidea lapicida* | - | MA | 46 | -62.6432 | -60.3726 | PV788354 | - |
| UR00984 | *Lecidea lapicida* | - | MA | NA | -62.693 | -60.3887 | PV788352 | - |
| MAF_LI_1c | *Lecidea lithophila* | Tr_A02 | CA | 5 | -62.6624 | -60.3877 | MK620090 | - |
| LE_A0511701 | *Lecidea polypycnidophora* | Tr_A02 | CA | 100 | -70.85 | 68.08333 | MK620081 | PV788494 |
| LE_A0513002 | *Lecidea polypycnidophora* | Tr_A02 | CA | 160 | -70.8167 | 68.08333 | GU074441 | JN204814 |
| LE_A150105 | *Lecidea polypycnidophora* | Tr_A02 | CA | 124 | -71.0982 | 71.47928 | - | - |
| T33632 | *Lecidea polypycnidophora* | Tr_A02 | CA | 25 | -77.6135 | 163.0718 | EU257681 | PV788521 |
| T33640 | *Lecidea polypycnidophora* | | CA | 32 | -77.6154 | 163.0639 | EU257682 | JN204737 |
| T33648 | *Lecidea polypycnidophora* | Tr_A02 | CA | 45 | -77.6269 | 163.0799 | EU257680 | JN204739 |
| T33703 | *Lecidea polypycnidophora* | - | CA | 39 | -77.6069 | 163.1126 | - | - |
| T46680 | *Lecidea polypycnidophora* | Tr_A02 | CA | 381 | -78.0275 | 163.8242 | MK970663 | MK970699 |
| T46713 | *Lecidea polypycnidophora* | Tr_A02 | CA | 350 | -78.0278 | 163.8433 | MK970663 | MK970698 |
| T46716 | *Lecidea polypycnidophora* | Tr_A02 | CA | 412 | -78.0403 | 163.8017 | MK970663 | MK970698 |
| T46717 | *Lecidea polypycnidophora* | Tr_A02 | CA | 442 | -78.04 | 163.8061 | MK970663 | MK970698 |
| T48773 | *Lecidea polypycnidophora* | Tr_A02 | CA | 377 | -78.1266 | 163.6737 | MK970663 | MK970699 |
| T48784 | *Lecidea polypycnidophora* | Tr_A02 | CA | 423 | -78.1214 | 163.6826 | MK970663 | MK970699 |
| T48859 | *Lecidea polypycnidophora* | Tr_A02 | CA | 319 | -78.0254 | 163.9857 | MK970663 | MK970698 |

| | | | | | | | | |
|---|---|---|---|---|---|---|---|---|
| T48880 | *Lecidea polypycnidophora* | - | CA | 443 | -78.0577 | 163.7403 | MK970674 | - |
| T48882 | *Lecidea polypycnidophora* | Tr_A02 | CA | 575 | -78.0568 | 163.8168 | MK970663 | MK970699 |
| MAF_LI_2a | *Lecidea promiscens* | Tr_I01 | CA | 15 | -62.6618 | -60.3817 | MK620091 | PV788660 |
| AAS_Convey00492A | *Lecidea silacea* | - | CA | 1323 | -75.83 | -69.37 | - | - |
| AAS_Convey00496 | *Lecidea silacea* | Tr_S02 | CA | 860 | -75.83 | -69.37 | PV788427 | - |
| AAS_Convey00507B | *Lecidea silacea* | Tr_S02 | CA | 860 | -75.83 | -69.37 | MK620077 | - |
| AAS_Convey00554b | *Lecidea silacea* | - | CA | 791 | -75.43 | -72.6 | - | - |
| AAS_Smith05103 | *Lecidea silacea* | Tr_A08 | MA | 100 | -60.6221 | -45.5777 | PV788428 | - |
| T48823b | *Lecidea sp. T48883b* | - | CA | 401 | -78.0981 | 163.7096 | MK620099 | - |
| UR01208 | *Lecidea sp. UR1208* | Tr_S02 | MA | 51 | -62.7481 | -60.3236 | PV788429 | - |
| UR00978 | *Lecidea sp.1* | - | MA | 46 | -62.6432 | -60.3726 | PV788431 | - |
| LE_A111504 | *Lecidea sp.2* | Tr_S02 | CA | 43 | -67.6734 | 45.81475 | MK620085 | - |
| UR00957 | *Lecidea sp.4* | - | MA | 50 | -62.6463 | -60.593 | PV788426 | - |
| LE_A152901 | *Lecidea sp.5* | Tr_A02 | CA | 888 | -72.963 | 65.63015 | PV788348 | - |
| T48883b | *Lecidea sp.5* | Tr_A02 | CA | 671 | -78.0576 | 163.8467 | MK620099 | MK970700 |
| T48781 | *Lecidea sp.6* | Tr_A02 | CA | 745 | -78.1276 | 163.6203 | MK620097 | MK970699 |
| T48857a | *Lecidea sp.6* | Tr_A02 | CA | 688 | -78.0363 | 163.827 | MK970684 | MK970699 |
| T48860b | *Lecidea sp.6* | Tr_A02 | CA | 694 | -78.0363 | 163.836 | MK970684 | MK970699 |
| LE_A133003 | *Lecidea UCR1* | Tr_A02 | CA | 134 | -72.197 | 68.82937 | MK620086 | PV788546 |
| T33619 | *Lecidea UCR1* | Tr_A04 | CA | 44 | -77.6116 | 163.0826 | EU263927 | JN204734 |
| T33658 | *Lecidea UCR1* | Tr_A02 | CA | 75 | -77.6162 | 163.0472 | - | - |

| | | | | | | | | |
|---|---|---|---|---|---|---|---|---|
| T33704 | *Lecidea UCR1* | Tr_A04 | CA | 62 | -77.6093 | 163.0796 | EU263928 | JN204741 |
| T48813 | *Lecidea UCR1* | Tr_A02 | CA | 557 | -78.1422 | 163.6571 | MK970699 | MK970699 |
| T48874 | *Lecidea UCR1* | Tr_A02 | CA | 375 | -78.024 | 163.8994 | MK970676 | MK970697 |
| UR01210 | *Lecidella drakensis* | Tr_I01 | MA | 51 | -62.7481 | -60.3236 | PV788486 | - |
| UR01214 | *Lecidella drakensis* | Tr_I01 | MA | 17 | -62.7512 | -60.3229 | PV788487 | - |
| LE_A0510504 | *Lecidella greenii* | Tr_A02 | CA | 50 | -70.8333 | 68.05 | - | PV788536 |
| LE_A130201 | *Lecidella greenii* | Tr_A02 | CA | 104 | -71.0991 | 71.47602 | - | PV788547 |
| LE_A131207 | *Lecidella greenii* | Tr_A02 | CA | 132 | -72.2149 | 68.82547 | - | PV788548 |
| LE_A132606 | *Lecidella greenii* | Tr_A02 | CA | 124 | -72.2114 | 68.80773 | - | PV788549 |
| LE_A133305 | *Lecidella greenii* | Tr_A02 | CA | 283 | -72.1948 | 68.8057 | - | - |
| LE_A150201 | *Lecidella greenii* | Tr_A02 | CA | 275 | -72.1023 | 68.739 | - | - |
| MAF_DP1_15 | *Lecidella greenii* | Tr_A02 | CA | 370 | -85.5392 | -151.15 | MK208718 | MK226852 |
| T33586 | *Lecidella greenii* | - | CA | 717 | -77.5772 | 163.0922 | JN873883 | - |
| T33611 | *Lecidella greenii* | - | CA | 60 | -77.6091 | 163.0813 | - | - |
| T33612 | *Lecidella greenii* | Tr_A02 | CA | 60 | -77.6091 | 163.0813 | JN873884 | PV788508 |
| T33620 | *Lecidella greenii* | | CA | 58 | -77.7144 | 163.0825 | - | JN204735 |
| T33628 | *Lecidella greenii* | - | CA | 60 | -77.6 | 163.0667 | JN873885 | - |
| T33630 | *Lecidella greenii* | Tr_A02 | CA | 60 | -77.6 | 163.0667 | - | JN204736 |
| T33649 | *Lecidella greenii* | - | CA | 45 | -77.6269 | 163.0799 | JN873886 | - |
| T33655 | *Lecidella greenii* | - | CA | 773 | -77.5544 | 163.3511 | JN873906 | - |
| T33700 | *Lecidella greenii* | Tr_A02 | CA | 45 | -77.4406 | 163.0923 | JN873887 | JN204740 |

| | | | | | | | | |
|---|---|---|---|---|---|---|---|---|
| T33706 | *Lecidella greenii* | Tr_A02 | CA | 60 | -77.6 | 163.1167 | JN873888 | JN204742 |
| T42999 | *Lecidella greenii* | Tr_A02 | CA | 755 | -77.0359 | 161.7345 | JN873889 | JN204774 |
| T43007 | *Lecidella greenii* | Tr_A02 | CA | 27 | -77.0066 | 162.5303 | JN873890 | PV788509 |
| T43013 | *Lecidella greenii* | Tr_A02 | CA | 94 | -77.0068 | 162.5452 | - | JN204780 |
| T43014 | *Lecidella greenii* | Tr_A02 | CA | 94 | -77.0068 | 162.5375 | - | JN204781 |
| T43015 | *Lecidella greenii* | Tr_A02 | CA | 6 | -77.0066 | 162.5246 | HQ287871 | JN204782 |
| T43017 | *Lecidella greenii* | Tr_A02 | CA | 8 | -77.0076 | 162.5192 | JN873891 | JN204784 |
| T43029 | *Lecidella greenii* | Tr_A02 | CA | 72 | -77.0068 | 162.5343 | JN873892 | JN204787 |
| T44636 | *Lecidella greenii* | Tr_A02 | CA | 492 | -79.8686 | 159.3575 | MK208776 | MK226906 |
| T46659 | *Lecidella greenii* | Tr_A02 | CA | 363 | -78.0231 | 163.903 | MK970671 | MK970698 |
| T46676 | *Lecidella greenii* | Tr_A02 | CA | 359 | -78.028 | 163.851 | MK970671 | MK970699 |
| T46706 | *Lecidella greenii* | Tr_A02 | CA | 379 | -78.028 | 163.821 | MK970671 | MK970698 |
| T48769 | *Lecidella greenii* | Tr_A02 | CA | 370 | -78.126 | 163.6998 | MK970671 | MK970669 |
| T48785 | *Lecidella greenii* | Tr_S02 | CA | 429 | -78.1327 | 163.6662 | MK970671 | MK970692 |
| T48787 | *Lecidella greenii* | Tr_A02 | CA | 361 | -78.127 | 163.678 | MK970671 | MK970699 |
| T48791a | *Lecidella greenii* | Tr_A02 | CA | 373 | -78.1198 | 163.6864 | MK970671 | MK970699 |
| T48801c | *Lecidella greenii* | Tr_A02 | CA | 732 | -78.1436 | 163.6258 | MK970671 | MK970699 |
| T48805 | *Lecidella greenii* | Tr_A02 | CA | 732 | -78.1436 | 163.6258 | MK970671 | MK970698 |
| T48809 | *Lecidella greenii* | Tr_A02 | CA | 540 | -78.1482 | 163.6565 | MK970671 | MK970699 |
| T48811a | *Lecidella greenii* | Tr_A02 | CA | 557 | -78.1422 | 163.6571 | MK970671 | MK970698 |
| T48812a | *Lecidella greenii* | Tr_A02 | CA | 540 | -78.1376 | 163.6186 | MK970671 | MK970699 |

| | | | | | | | | |
|---|---|---|---|---|---|---|---|---|
| T48812b | *Lecidella greenii* | Tr_A02 | CA | 540 | -78.1376 | 163.6186 | MK970671 | MK970699 |
| T48828a | *Lecidella greenii* | - | CA | 406 | -78.1096 | 163.8578 | MK970671 | - |
| T48828b | *Lecidella greenii* | Tr_A02 | CA | 406 | -78.1096 | 163.8578 | MK970671 | MK970698 |
| T48829 | *Lecidella greenii* | Tr_A02 | CA | 453 | -78.1113 | 163.8585 | MK970671 | MK970699 |
| T48837 | *Lecidella greenii* | Tr_A02 | CA | 179 | -78.0981 | 163.7772 | MK970671 | MK970699 |
| T48839 | *Lecidella greenii* | Tr_A02 | CA | 514 | -78.1138 | 163.8543 | MK970671 | MK970699 |
| T48844 | *Lecidella greenii* | Tr_A02 | CA | 749 | -78.0667 | 163.8633 | MK970671 | MK970698 |
| T48858 | *Lecidella greenii* | - | CA | 319 | -78.0251 | 163.9751 | MK970671 | - |
| T48869 | *Lecidella greenii* | - | CA | 849 | -78.0367 | 163.9783 | MK970671 | - |
| T48872 | *Lecidella greenii* | - | CA | 673 | -78.0303 | 163.9514 | MK970671 | - |
| T48873 | *Lecidella greenii* | Tr_A02 | CA | 455 | -78.0245 | 163.8932 | MK970671 | MK970698 |
| T48876a | *Lecidella greenii* | Tr_A02 | CA | 380 | -78.024 | 163.9 | MK970671 | MK970698 |
| T48876b | *Lecidella greenii* | Tr_A02 | CA | 380 | -78.024 | 163.9 | MK970671 | MK970699 |
| T48879 | *Lecidella greenii* | - | CA | 419 | -78.0567 | 163.747 | MK970671 | - |
| UR00772 | *Lecidella greenii* | Tr_S07 | CA | 5 | -77.0167 | 162.5333 | PV788439 | PV788514 |
| AAS_Smith05524 | *Lecidella siplei* | - | CA | 1875 | -81.83 | -89.5 | - | - |
| LE_A0513410 | *Lecidella siplei* | - | CA | 80 | -70.8667 | 67.95 | - | - |
| LE_A131001 | *Lecidella siplei* | Tr_A02 | CA | 127 | -72.218 | 68.81952 | - | - |
| LE_A150504 | *Lecidella siplei* | Tr_A02 | CA | 248 | -72.102 | 68.74618 | - | - |
| LE_A152701 | *Lecidella siplei* | Tr_S02 | CA | 338 | -73.0912 | 66.44153 | PV788392 | PV788604 |
| MAF_DP1_22 | *Lecidella siplei* | Tr_A02 | CA | 370 | -85.5392 | -151.15 | MK208721 | MK226854 |

| | | | | | | | | |
|---|---|---|---|---|---|---|---|---|
| MAF_DP1_35 | *Lecidella siplei* | Tr_A02 | CA | 370 | -85.5392 | -151.15 | MK208729 | MK226862 |
| MAF_DP1_57 | *Lecidella siplei* | Tr_A02 | CA | 370 | -85.5392 | -151.15 | MK208736 | MK226869 |
| MAF_GS1_13 | *Lecidella siplei* | Tr_A02 | CA | 162 | -84.5355 | -174.954 | MK208739 | MK226872 |
| MAF_GS1_58 | *Lecidella siplei* | Tr_A02 | CA | 162 | -84.5355 | -174.954 | MK208742 | MK226875 |
| MAF_GS1_60 | *Lecidella siplei* | Tr_A02 | CA | 162 | -84.5355 | -174.954 | MK208743 | MK226876 |
| MAF_GS1_62 | *Lecidella siplei* | Tr_A02 | CA | 162 | -84.5355 | -174.954 | MK208745 | MK226878 |
| T32987 | *Lecidella siplei* | Tr_A02 | CA | 15 | -77.0167 | 162.5333 | JN873895 | PV788510 |
| T32991 | *Lecidella siplei* | Tr_A02 | CA | 15 | -77.0167 | 162.5333 | JN873895 | JN204726 |
| T32992 | *Lecidella siplei* | - | CA | 15 | -77.0167 | 162.5333 | JN873896 | - |
| T33449 | *Lecidella siplei* | Tr_A02 | CA | 530 | -83.7606 | 172.755 | JN873897 | JN204729 |
| T33454 | *Lecidella siplei* | Tr_S02 | CA | 530 | -83.7606 | 172.755 | - | PV788576 |
| T33455 | *Lecidella siplei* | Tr_A02 | CA | 530 | -83.7606 | 172.755 | - | JN204730 |
| T33457 | *Lecidella siplei* | Tr_A02 | CA | 530 | -83.7606 | 172.755 | JN873898 | JN204731 |
| T33458 | *Lecidella siplei* | - | CA | 530 | -83.7606 | 172.755 | - | - |
| T33667 | *Lecidella siplei* | Tr_A02 | CA | 40 | -77.6021 | 163.1174 | - | - |
| T35540 | *Lecidella siplei* | Tr_A02 | CA | 405 | -79.8417 | 159.3631 | JN873898 | PV788533 |
| T35895 | *Lecidella siplei* | Tr_A02 | CA | 15 | -77.0071 | 162.6298 | JN873899 | JN204757 |
| T35909 | *Lecidella siplei* | - | CA | 15 | -77.0071 | 162.6298 | - | - |
| T43004 | *Lecidella siplei* | Tr_A02 | CA | 12 | -77.006 | 162.5318 | - | JN204776 |
| T43033 | *Lecidella siplei* | Tr_A02 | CA | 72 | -77.0068 | 162.5343 | - | JN204791 |
| UR01202 | *Lecidella siplei* | Tr_A12 | MA | 51 | -62.7481 | -60.3236 | PV788444 | PV788557 |

| | | | | | | | | |
|---|---|---|---|---|---|---|---|---|
| UR01204 | *Lecidella siplei* | Tr_A12 | MA | 51 | -62.7481 | -60.3236 | PV788445 | PV788559 |
| AAS_Smith03154B | *Lecidella sp.* | - | MA | 85 | -60.6221 | -45.5777 | - | - |
| LE_A0512002 | *Lecidella sp.* | - | CA | 100 | -70.85 | 68.08333 | - | - |
| LE_A0512301 | *Lecidella sp.* | Tr_A02 | CA | 40 | -70.7833 | 68.16667 | PV788347 | - |
| LE_A0513310 | *Lecidella sp.* | - | CA | 80 | -70.8667 | 67.95 | - | - |
| LE_A0513801 | *Lecidella sp.* | - | CA | 100 | -70.8667 | 67.93333 | - | - |
| LE_A0514504 | *Lecidella sp.* | - | CA | 20 | -70.85 | 68.01667 | - | - |
| LE_A058001 | *Lecidella sp. B* | - | CA | 30 | -69.4083 | 76.4 | - | - |
| LE_RU1_5 | *Lecidella sp. B* | - | CA | 127 | -74.7652 | -136.803 | - | - |
| LE_A041305 | *Lecidella sp. LE A041305* | Tr_A13 | CA | 70 | -69.7489 | 73.68248 | - | - |
| LE_A060415 | *Lecidella sp. LE A060415* | Tr_I01 | CA | 30 | -65.1 | -64.05 | - | - |
| UR00899 | *Lecidella sp.1* | - | MA | 250 | -62.6683 | -60.3799 | PV788475 | - |
| UR00970 | *Lecidella sp.1* | - | MA | 55 | -62.6671 | -60.394 | PV788476 | - |
| UR00985 | *Lecidella sp.1* | Tr_S02 | MA | NA | -62.693 | -60.3887 | PV788477 | - |
| MAF_GS1_64 | *Lecidella sp.2* | Tr_S02 | CA | 162 | -84.5355 | -174.954 | MK208746 | MK226879 |
| MAF_HS7_59 | *Lecidella sp.2* | Tr_S02 | CA | 1012 | -83.8057 | 172.2625 | MK208747 | MK226880 |
| MAF_MK1_18 | *Lecidella sp.2* | Tr_S02 | CA | 859 | -83.7748 | 171.8276 | MK208753 | MK226886 |
| MAF_MK1_21 | *Lecidella sp.2* | Tr_S02 | CA | 859 | -83.7748 | 171.8276 | MK208754 | MK226887 |
| MAF_MK1_27 | *Lecidella sp.2* | Tr_S02 | CA | 859 | -83.7748 | 171.8276 | MK208756 | MK226889 |
| MAF_MK1_43 | *Lecidella sp.2* | Tr_S02 | CA | 859 | -83.7748 | 171.8276 | MK208761 | MK226894 |
| MAF_MK1_49 | *Lecidella sp.2* | Tr_S02 | CA | 859 | -83.7748 | 171.8276 | MK208763 | MK226896 |

| | | | | | | | | |
|---|---|---|---|---|---|---|---|---|
| T33335 | *Lecidella sp.2* | Tr_S18 | CA | 810 | -83.8033 | 172.2067 | MK208768 | MK226899 |
| T33338 | *Lecidella sp.2* | Tr_S02 | CA | 810 | -83.8033 | 172.2067 | MK208769 | MK226900 |
| UR00860 | *Lecidella sp.3* | Tr_I01 | MA | 2 | -62.6606 | -60.3816 | PV788446 | PV788649 |
| UR00873 | *Lecidella sp.3* | Tr_S02 | MA | 61 | -62.665 | -60.3784 | PV788451 | PV788602 |
| UR00879 | *Lecidella sp.3* | Tr_S02 | MA | 40 | -62.6618 | -60.3787 | PV788447 | PV788594 |
| UR00880 | *Lecidella sp.3* | - | MA | 35 | -62.6609 | -60.3799 | PV788448 | - |
| UR00882 | *Lecidella sp.3* | - | MA | 35 | -62.6609 | -60.3799 | PV788449 | - |
| UR00883 | *Lecidella sp.3* | - | MA | 35 | -62.6609 | -60.3799 | PV788463 | - |
| UR00884 | *Lecidella sp.3* | - | MA | 35 | -62.6609 | -60.3799 | PV788462 | - |
| UR00886 | *Lecidella sp.3* | - | MA | 44 | -62.6628 | -60.3784 | PV788452 | - |
| UR00921 | *Lecidella sp.3* | - | MA | 383 | -62.6816 | -60.3422 | PV788454 | - |
| UR00930 | *Lecidella sp.3* | - | MA | 383 | -62.6816 | -60.3422 | PV788456 | - |
| UR00959 | *Lecidella sp.3* | - | MA | 50 | -62.6463 | -60.593 | PV788457 | - |
| UR00960 | *Lecidella sp.3* | Tr_I01 | MA | 56 | -62.649 | -60.5983 | PV788455 | PV788641 |
| UR00961 | *Lecidella sp.3* | - | MA | 104 | -62.6495 | -60.5998 | PV788464 | - |
| UR00962 | *Lecidella sp.3* | - | MA | 104 | -62.6495 | -60.5998 | PV788465 | - |
| UR00963 | *Lecidella sp.3* | - | MA | 104 | -62.6495 | -60.5998 | PV788466 | - |
| UR00964 | *Lecidella sp.3* | - | MA | 104 | -62.6495 | -60.5998 | PV788467 | - |
| UR00968 | *Lecidella sp.3* | - | MA | 55 | -62.6671 | -60.394 | PV788453 | - |
| UR00972 | *Lecidella sp.3* | Tr_S02 | MA | 46 | -62.6432 | -60.3726 | PV788458 | PV788595 |
| UR00973 | *Lecidella sp.3* | - | MA | 46 | -62.6432 | -60.3726 | PV788461 | - |

| | | | | | | | | |
|---|---|---|---|---|---|---|---|---|
| UR00976 | *Lecidella sp.3* | - | MA | 46 | -62.6432 | -60.3726 | PV788459 | - |
| UR00977 | *Lecidella sp.3* | - | MA | 46 | -62.6432 | -60.3726 | PV788450 | - |
| UR00980 | *Lecidella sp.3* | - | MA | 46 | -62.6432 | -60.3726 | PV788460 | - |
| UR00894 | *Lecidella sp.4* | - | MA | 310 | -62.6691 | -60.3809 | PV788478 | - |
| UR00896 | *Lecidella sp.4* | - | MA | 310 | -62.6691 | -60.3809 | PV788479 | - |
| UR00908 | *Lecidella sp.4* | - | MA | 115 | -62.6666 | -60.3946 | PV788480 | - |
| UR00909 | *Lecidella sp.4* | - | MA | 115 | -62.6666 | -60.3946 | PV788481 | - |
| UR00966 | *Lecidella sp.4* | Tr_S02 | MA | 55 | -62.6671 | -60.394 | PV788482 | - |
| UR00983 | *Lecidella sp.4* | - | MA | NA | -62.693 | -60.3887 | PV788483 | - |
| UR01201 | *Lecidella sp.5* | Tr_S02 | MA | 51 | -62.7481 | -60.3236 | PV788488 | - |
| UR01205 | *Lecidella sp.5* | Tr_S02 | MA | 51 | -62.7481 | -60.3236 | PV788490 | - |
| UR01211 | *Lecidella sp.5* | Tr_S02 | MA | 51 | -62.7481 | -60.3236 | PV788489 | - |
| AAS_Smith06906B | *Lecidella stigmatea* | - | MA | 110 | -62.9827 | -60.711 | - | - |
| LE_A0510701 | *Lecidella stigmatea* | - | CA | 50 | -70.8333 | 68.05 | - | - |
| LE_A0513103 | *Lecidella stigmatea* | Tr_A02 | CA | 80 | -70.8667 | 67.95 | - | |
| LE_A890221 | *Lecidella stigmatea* | - | CA | 20 | -67.9623 | 44.58029 | - | - |
| LE_A892702 | *Lecidella stigmatea* | - | CA | 144 | -66.3 | 100.75 | - | - |
| UR00861 | *Lecidella stigmatea* | Tr_I01 | MA | 2 | -62.6606 | -60.3816 | PV788440 | PV788648 |
| UR00956 | *Lecidella stigmatea* | - | MA | 107 | -62.665 | -60.3784 | PV788441 | - |
| UR01206 | *Lecidella stigmatea* | Tr_A12 | MA | 51 | -62.7481 | -60.3236 | PV788443 | PV788558 |
| UR01209 | *Lecidella stigmatea* | Tr_A13 | MA | 51 | -62.7481 | -60.3236 | PV788442 | PV788563 |

| | | | | | | | | |
|---|---|---|---|---|---|---|---|---|
| UR00902 | *Poeltidea perusta* | - | MA | 142 | -62.6666 | -60.392 | PV788437 | - |
| LE_A0517306 | *Poeltidea sp. 1* | Tr_A02 | MA | 30 | -62.2067 | -58.9667 | PV788432 | - |
| T37137 | *Poeltidea sp. 1* | - | MA | 60 | -62.6667 | -61.1 | PV788433 | - |
| UR00988 | *Poeltidea sp. 1* | - | MA | NA | -62.693 | -60.3887 | PV788434 | - |
| UR00989 | *Poeltidea sp. 1* | - | MA | NA | -62.693 | -60.3887 | PV788435 | - |
| UR00990 | *Poeltidea sp. 1* | - | MA | NA | -62.693 | -60.3887 | PV788436 | - |
| ADT13352 | *Porpidia macrocarpa* | Tr_A04 | CA | NA | -66.3183 | 110.5167 | PV788491 | JN204824 |
| ADT18027 | *Porpidia macrocarpa* | Tr_A04 | CA | NA | -66.2822 | 110.5439 | PV788492 | JN204826 |
| MAF_LI_4a | *Porpidia macrocarpa* | Tr_I01 | MA | 15 | -62.6618 | -60.3817 | MK620095 | PV788642 |
| MAF_LI_1a | *Porpidia sp.1* | Tr_A02 | CA | 5 | -62.6624 | -60.3877 | MK620089 | PV788511 |
| MAF_LI_2c | *Porpidia sp.3* | Tr_I01 | CA | 15 | -62.6618 | -60.3817 | MK620092 | - |
| T33597 | *Rhizoplaca macleanii* | - | CA | 604 | -77.5739 | 163.1336 | JN873905 | - |
| T33626 | *Rhizoplaca macleanii* | Tr_A02 | CA | 718 | -77.5818 | 163.0843 | - | PV788512 |
| T33726 | *Rhizoplaca macleanii* | Tr_A02 | CA | 950 | -77.6699 | 163.1501 | JN873907 | - |
| T33736 | *Rhizoplaca macleanii* | - | CA | 950 | -77.6699 | 163.1501 | - | - |
| T43032 | *Rhizoplaca macleanii* | Tr_A02 | CA | 598 | -77.5701 | 163.157 | JN873908 | JN204790 |
| T43093 | *Rhizoplaca macleanii* | Tr_A02 | CA | 40 | -77.8481 | 166.7503 | - | PV788513 |
| T46643 | *Rhizoplaca macleanii* | Tr_A02 | CA | 602 | -78.0314 | 163.8653 | MK970663 | MK970698 |
| T46647b | *Rhizoplaca macleanii* | Tr_A02 | CA | 794 | -78.0328 | 163.8975 | MK970663 | MK970698 |
| T46672 | *Rhizoplaca macleanii* | Tr_A02 | CA | 397 | -78.0281 | 163.8514 | MK970663 | MK970698 |
| T46673 | *Rhizoplaca macleanii* | Tr_A02 | CA | 365 | -78.0269 | 163.8483 | MK970663 | MK970699 |

| | | | | | | | | |
|---|---|---|---|---|---|---|---|---|
| T46678 | *Rhizoplaca macleanii* | Tr_A02 | CA | 843 | -78.0189 | 163.9706 | MK970669 | MK970669 |
| T46679 | *Rhizoplaca macleanii* | Tr_A02 | CA | 790 | -78.0319 | 163.9506 | MK970664 | MK970696 |
| T46681 | *Rhizoplaca macleanii* | Tr_A02 | CA | 870 | -78.0442 | 163.9861 | MK970663 | MK970698 |
| T46701 | *Rhizoplaca macleanii* | Tr_A02 | CA | 390 | -78.02 | 163.805 | MK970663 | MK970698 |
| T46710 | *Rhizoplaca macleanii* | Tr_A02 | CA | 600 | -78.073 | 163.717 | MK970666 | MK970698 |
| T48774 | *Rhizoplaca macleanii* | Tr_A02 | CA | 526 | -78.1352 | 163.6262 | MK970663 | MK970698 |
| T48778a | *Rhizoplaca macleanii* | Tr_A02 | CA | 700 | -78.149 | 163.7692 | MK970663 | MK970698 |
| T48779 | *Rhizoplaca macleanii* | Tr_A02 | CA | 757 | -78.1641 | 163.7546 | MK970664 | MK970696 |
| T48794b | *Rhizoplaca macleanii* | Tr_A02 | CA | 905 | -78.151 | 163.7347 | MK970663 | MK970689 |
| T48795a | *Rhizoplaca macleanii* | Tr_A02 | CA | 853 | -78.1614 | 163.7136 | MK970668 | MK970698 |
| T48797 | *Rhizoplaca macleanii* | Tr_A02 | CA | 814 | -78.156 | 163.6891 | MK970667 | MK970698 |
| T48798 | *Rhizoplaca macleanii* | Tr_A02 | CA | 911 | -78.1505 | 163.7363 | MK970667 | MK970698 |
| T48804 | *Rhizoplaca macleanii* | Tr_A02 | CA | 775 | -78.1483 | 163.6296 | MK970670 | MK970699 |
| T48817a | *Rhizoplaca macleanii* | Tr_A02 | CA | 546 | -78.1127 | 163.7845 | MK970663 | MK970698 |
| T48821 | *Rhizoplaca macleanii* | Tr_A02 | CA | 589 | -78.1144 | 163.7787 | MK970665 | MK970698 |
| T48841a | *Rhizoplaca macleanii* | Tr_A02 | CA | 722 | -78.0681 | 163.8614 | MK970663 | MK970698 |
| T48843d | *Rhizoplaca macleanii* | Tr_A02 | CA | 836 | -78.0659 | 163.8699 | MK970663 | MK970699 |
| T48843e | *Rhizoplaca macleanii* | Tr_A02 | CA | 836 | -78.0659 | 163.8699 | MK970663 | MK970702 |
| T48865a | *Rhizoplaca macleanii* | Tr_A02 | CA | 874 | -78.0363 | 163.9903 | - | MK970698 |
| T48867a | *Rhizoplaca macleanii* | Tr_A02 | CA | 936 | -78.0428 | 164.1038 | MK970663 | MK970696 |
| T48867c | *Rhizoplaca macleanii* | Tr_A02 | CA | 936 | -78.0428 | 164.1038 | MK970666 | MK970702 |

| | | | | | | | | |
|---|---|---|---|---|---|---|---|---|
| T48881b | *Rhizoplaca macleanii* | Tr_A02 | CA | 541 | -78.0612 | 163.7909 | MK970666 | MK970699 |
| T48885 | *Rhizoplaca macleanii* | Tr_A02 | CA | 666 | -78.0573 | 163.8442 | MK970666 | KM970698 |
| T48900 | *Rhizoplaca macleanii* | Tr_A02 | CA | 610 | -78.0335 | 163.8448 | MK970663 | MK970698 |
| T54141 | *Rhizoplaca macleanii* | - | CA | 15 | -77.0081 | 162.5183 | - | - |
| T54147 | *Rhizoplaca macleanii* | Tr_A02 | CA | 15 | -77.0142 | 162.5878 | - | - |
| T54152 | *Rhizoplaca macleanii* | - | CA | 9 | -77.0172 | 162.4653 | - | - |
| AAS_Smith11907A | *Rhizoplaca melanophthalma* | - | MA | 452 | -70.7 | -69.68 | - | - |
| LE_A0513402 | *Rhizoplaca melanophthalma* | - | CA | 80 | -70.8667 | 67.95 | - | - |
| T35508 | *Rhizoplaca melanophthalma* | Tr_I01 | CA | 90 | -72.3231 | 170.2319 | - | - |

**Table S2** Primers used in this study with references.

| Symbiont | Primer Name | Reference |
|---|---|---|
| Mycobiont general | ITS1 | (White et al., 1990) |
| | ITS1F | (Gardes & Bruns, 1993) |
| | ITS1L | (Ruprecht et al., 2020) |
| | ITS4 | (White et al., 1990) |
| | ITS4L | (Ruprecht et al., 2020) |
| *Trebouxia* | ITS1T | (Kroken & Taylor, 2000) |
| | ITS1aT | (Ruprecht et al., 2014) |
| | ITS4T | (Kroken & Taylor, 2000) |
| | ITS4bT | (Ruprecht et al., 2014) |
| *Chlorella* | Ns7m | (Bhattacharya et al., 1996) |
| | LR1850 | |
| *Asterochloris* | ITS2-sense-A | (Ruprecht et al., 2014) |
| | ITS2-antisense-A | |

**Table S3** Variable Contribution to the PC1 and PC2 axes for current niches (probability >= 0.75) for mycobiont species and photobiont OTU.

| **Bioclim variable** | **PC1** | **PC2** |
|---|---|---|
| bio1 | **0.41** | 0.01 |
| bio12 | **0.38** | -0.04 |
| bio15 | -0.25 | **-0.92** |
| bio18 | **0.41** | 0.01 |
| bio2 | -0.36 | **0.29** |
| bio7 | **-0.42** | 0.04 |
| bio8 | 0.39 | -0.26 |

**Table S4** Delta Area mean, min and max area gain (in 100 $km^2$) of probable niches per species across regions. Underlined vales show the highest delta area mean values per mycobiont species and photobiont OTU. Bold values show the newly acquired ranges for each respective region.

| | | | Delta Area | | |
|---|---|---|---|---|---|
| **Scenario** | **Region** | **Mycobiont species/ Photobiont OTU** | **Mean** | **Min** | **Max** |
| SSP1-2.6 | ANTARCTIC PENINSULA | *Carbonea vorticosa* | -0.09 | -0.09 | -0.09 |
| SSP1-2.6 | ANTARCTIC PENINSULA | *Lecanora fuscobrunnea* | 0.00 | 0.00 | 0.00 |
| SSP1-2.6 | ANTARCTIC PENINSULA | *Lecidea andersonii* | 0.53 | -1.84 | 4.29 |
| SSP1-2.6 | ANTARCTIC PENINSULA | *Lecidea atrobrunnea* | 16.62 | 3.24 | 36.79 |
| SSP1-2.6 | ANTARCTIC PENINSULA | *Lecidea cancriformis* | -4.97 | -9.90 | 2.10 |
| SSP1-2.6 | ANTARCTIC PENINSULA | *Lecidea polypycnidophora* | 0.00 | 0.00 | 0.00 |
| SSP1-2.6 | ANTARCTIC PENINSULA | *Lecidella greenii* | 0.00 | 0.00 | 0.00 |
| SSP1-2.6 | ANTARCTIC PENINSULA | *Lecidella siplei* | 0.00 | 0.00 | 0.00 |
| SSP1-2.6 | ANTARCTIC PENINSULA | *Rhizoplaca macleanii* | 0.00 | 0.00 | 0.00 |
| SSP1-2.6 | ANTARCTIC PENINSULA | *Tr*_A02 | 1.07 | -1.31 | 2.63 |
| SSP1-2.6 | ANTARCTIC PENINSULA | *Tr*_I01 | 39.48 | 18.83 | 60.61 |
| SSP1-2.6 | ANTARCTIC PENINSULA | *Tr*_S02 | -0.09 | -0.35 | 0.44 |
| SSP1-2.6 | ANTARCTIC PENINSULA | *Tr*_S18 | 1.40 | 0.44 | 2.98 |
| SSP1-2.6 | PENSACOLA MOUNTAINS | *Carbonea vorticosa* | 0.63 | -1.49 | 3.15 |
| SSP1-2.6 | PENSACOLA MOUNTAINS | *Lecanora fuscobrunnea* | 4.73 | -0.09 | 12.26 |
| SSP1-2.6 | PENSACOLA MOUNTAINS | *Lecidea andersonii* | 3.17 | 2.01 | 4.38 |
| SSP1-2.6 | PENSACOLA MOUNTAINS | *Lecidea atrobrunnea* | 0.00 | 0.00 | 0.00 |
| SSP1-2.6 | PENSACOLA MOUNTAINS | *Lecidea cancriformis* | 0.98 | -0.44 | 3.07 |
| SSP1-2.6 | PENSACOLA MOUNTAINS | *Lecidea polypycnidophora* | 0.00 | 0.00 | 0.00 |
| SSP1-2.6 | PENSACOLA MOUNTAINS | *Lecidella greenii* | 0.58 | 0.26 | 0.88 |
| SSP1-2.6 | PENSACOLA MOUNTAINS | *Lecidella siplei* | 0.46 | 0.18 | 0.88 |
| SSP1-2.6 | PENSACOLA MOUNTAINS | *Rhizoplaca macleanii* | 0.00 | 0.00 | 0.00 |
| SSP1-2.6 | PENSACOLA MOUNTAINS | *Tr*_A02 | 2.49 | 0.70 | 5.26 |
| SSP1-2.6 | PENSACOLA MOUNTAINS | *Tr*_I01 | 0.72 | -0.09 | 2.45 |
| SSP1-2.6 | PENSACOLA MOUNTAINS | *Tr*_S02 | 7.06 | 4.12 | 10.77 |

| | | | | | |
|---|---|---|---|---|---|
| SSP1-2.6 | PENSACOLA MOUNTAINS | *Tr*_S18 | 3.80 | 0.88 | 7.44 |
| SSP1-2.6 | PRINCE CHARLES MOUNTAINS | *Carbonea vorticosa* | 1.51 | -0.61 | 4.82 |
| SSP1-2.6 | PRINCE CHARLES MOUNTAINS | *Lecanora fuscobrunnea* | 3.15 | 1.58 | 5.96 |
| SSP1-2.6 | PRINCE CHARLES MOUNTAINS | *Lecidea andersonii* | <u>10.05</u> | 5.96 | 13.31 |
| SSP1-2.6 | PRINCE CHARLES MOUNTAINS | *Lecidea atrobrunnea* | 0.00 | 0.00 | 0.00 |
| SSP1-2.6 | PRINCE CHARLES MOUNTAINS | *Lecidea cancriformis* | 3.47 | 0.26 | 6.39 |
| SSP1-2.6 | PRINCE CHARLES MOUNTAINS | *Lecidea polypycnidophora* | 1.24 | 0.09 | 2.54 |
| SSP1-2.6 | PRINCE CHARLES MOUNTAINS | *Lecidella greenii* | 5.17 | 2.36 | 10.77 |
| SSP1-2.6 | PRINCE CHARLES MOUNTAINS | *Lecidella siplei* | 0.51 | 0.00 | 1.23 |
| SSP1-2.6 | PRINCE CHARLES MOUNTAINS | *Rhizoplaca macleanii* | 0.07 | -0.79 | 1.05 |
| SSP1-2.6 | PRINCE CHARLES MOUNTAINS | *Tr*_A02 | <u>10.49</u> | 8.58 | 12.79 |
| SSP1-2.6 | PRINCE CHARLES MOUNTAINS | *Tr*_I01 | 3.75 | 2.54 | 5.08 |
| SSP1-2.6 | PRINCE CHARLES MOUNTAINS | *Tr*_S02 | 8.93 | 5.96 | 13.84 |
| SSP1-2.6 | PRINCE CHARLES MOUNTAINS | *Tr*_S18 | 4.48 | 0.53 | 9.90 |
| SSP1-2.6 | TRANSANTARCTIC MOUNTAINS | *Carbonea vorticosa* | 2.68 | -0.53 | 7.09 |
| SSP1-2.6 | TRANSANTARCTIC MOUNTAINS | *Lecanora fuscobrunnea* | 13.84 | 5.52 | 33.28 |
| SSP1-2.6 | TRANSANTARCTIC MOUNTAINS | *Lecidea andersonii* | <u>23.77</u> | 10.25 | 42.30 |
| SSP1-2.6 | TRANSANTARCTIC MOUNTAINS | *Lecidea atrobrunnea* | 0.00 | 0.00 | 0.00 |
| SSP1-2.6 | TRANSANTARCTIC MOUNTAINS | *Lecidea cancriformis* | 14.12 | 3.24 | 36.96 |
| SSP1-2.6 | TRANSANTARCTIC MOUNTAINS | *Lecidea polypycnidophora* | 0.63 | -1.58 | 2.63 |
| SSP1-2.6 | TRANSANTARCTIC MOUNTAINS | *Lecidella greenii* | 12.72 | 7.18 | 23.82 |
| SSP1-2.6 | TRANSANTARCTIC MOUNTAINS | *Lecidella siplei* | 11.19 | 3.50 | 26.19 |
| SSP1-2.6 | TRANSANTARCTIC MOUNTAINS | *Rhizoplaca macleanii* | 0.79 | -0.70 | 3.33 |
| SSP1-2.6 | TRANSANTARCTIC MOUNTAINS | *Tr*_A02 | <u>22.32</u> | 9.11 | 39.68 |
| SSP1-2.6 | TRANSANTARCTIC MOUNTAINS | *Tr*_I01 | 15.89 | 3.07 | 35.73 |
| SSP1-2.6 | TRANSANTARCTIC MOUNTAINS | *Tr*_S02 | 18.34 | 10.25 | 31.97 |
| SSP1-2.6 | TRANSANTARCTIC MOUNTAINS | *Tr*_S18 | 10.63 | -2.10 | 34.07 |
| SSP5-8.5 | ANTARCTIC PENINSULA | *Carbonea vorticosa* | -0.09 | -0.09 | -0.09 |
| SSP5-8.5 | ANTARCTIC PENINSULA | *Lecanora fuscobrunnea* | 0.00 | 0.00 | 0.00 |
| SSP5-8.5 | ANTARCTIC PENINSULA | *Lecidea andersonii* | -2.68 | -5.43 | -0.44 |
| SSP5-8.5 | ANTARCTIC PENINSULA | *Lecidea atrobrunnea* | <u>65.48</u> | 15.59 | 137.16 |
| SSP5-8.5 | ANTARCTIC PENINSULA | *Lecidea cancriformis* | -6.85 | -11.12 | -0.26 |
| SSP5-8.5 | ANTARCTIC PENINSULA | *Lecidea polypycnidophora* | 0.00 | 0.00 | 0.00 |
| SSP5-8.5 | ANTARCTIC PENINSULA | *Lecidella greenii* | 0.00 | 0.00 | 0.00 |
| SSP5-8.5 | ANTARCTIC PENINSULA | *Lecidella siplei* | 0.00 | 0.00 | 0.00 |

| | | | | | |
|---|---|---|---|---|---|
| SSP5-8.5 | ANTARCTIC PENINSULA | *Rhizoplaca macleanii* | 0.00 | 0.00 | 0.00 |
| SSP5-8.5 | ANTARCTIC PENINSULA | *Tr*_A02 | 15.73 | -4.29 | 50.80 |
| SSP5-8.5 | ANTARCTIC PENINSULA | *Tr*_I01 | 72.75 | -27.33 | 114.47 |
| SSP5-8.5 | ANTARCTIC PENINSULA | *Tr*_S02 | -0.35 | -0.35 | -0.35 |
| SSP5-8.5 | ANTARCTIC PENINSULA | *Tr*_S18 | -0.11 | -0.79 | 1.40 |
| SSP5-8.5 | PENSACOLA MOUNTAINS | *Carbonea vorticosa* | 1.54 | -1.05 | 3.94 |
| SSP5-8.5 | PENSACOLA MOUNTAINS | *Lecanora fuscobrunnea* | 9.41 | -6.66 | 21.11 |
| SSP5-8.5 | PENSACOLA MOUNTAINS | *Lecidea andersonii* | **13.72** | 0.26 | 19.71 |
| SSP5-8.5 | PENSACOLA MOUNTAINS | *Lecidea atrobrunnea* | 0.00 | 0.00 | 0.00 |
| SSP5-8.5 | PENSACOLA MOUNTAINS | *Lecidea cancriformis* | 3.64 | 0.53 | 6.31 |
| SSP5-8.5 | PENSACOLA MOUNTAINS | *Lecidea polypycnidophora* | **1.00** | 0.00 | 1.75 |
| SSP5-8.5 | PENSACOLA MOUNTAINS | *Lecidella greenii* | **3.38** | 1.23 | 8.32 |
| SSP5-8.5 | PENSACOLA MOUNTAINS | *Lecidella siplei* | **4.99** | 0.00 | 11.91 |
| SSP5-8.5 | PENSACOLA MOUNTAINS | *Rhizoplaca macleanii* | 0.40 | 0.00 | 0.88 |
| SSP5-8.5 | PENSACOLA MOUNTAINS | *Tr*_A02 | 16.83 | 4.38 | 23.74 |
| SSP5-8.5 | PENSACOLA MOUNTAINS | *Tr*_I01 | 4.76 | -0.09 | 11.21 |
| SSP5-8.5 | PENSACOLA MOUNTAINS | *Tr*_S02 | 17.90 | 0.88 | 24.00 |
| SSP5-8.5 | PENSACOLA MOUNTAINS | *Tr*_S18 | 0.00 | -15.06 | 7.97 |
| SSP5-8.5 | PRINCE CHARLES MOUNTAINS | *Carbonea vorticosa* | 0.82 | -1.66 | 4.99 |
| SSP5-8.5 | PRINCE CHARLES MOUNTAINS | *Lecanora fuscobrunnea* | 3.92 | -1.84 | 10.51 |
| SSP5-8.5 | PRINCE CHARLES MOUNTAINS | *Lecidea andersonii* | 17.57 | -3.50 | 30.65 |
| SSP5-8.5 | PRINCE CHARLES MOUNTAINS | *Lecidea atrobrunnea* | 0.00 | 0.00 | 0.00 |
| SSP5-8.5 | PRINCE CHARLES MOUNTAINS | *Lecidea cancriformis* | 5.75 | -2.54 | 17.95 |
| SSP5-8.5 | PRINCE CHARLES MOUNTAINS | *Lecidea polypycnidophora* | 4.15 | 1.05 | 7.18 |
| SSP5-8.5 | PRINCE CHARLES MOUNTAINS | *Lecidella greenii* | 9.51 | 6.13 | 11.12 |
| SSP5-8.5 | PRINCE CHARLES MOUNTAINS | *Lecidella siplei* | **1.89** | 0.00 | 3.85 |
| SSP5-8.5 | PRINCE CHARLES MOUNTAINS | *Rhizoplaca macleanii* | -0.11 | -0.79 | 0.96 |
| SSP5-8.5 | PRINCE CHARLES MOUNTAINS | *Tr*_A02 | 18.13 | 0.88 | 25.75 |
| SSP5-8.5 | PRINCE CHARLES MOUNTAINS | *Tr*_I01 | 7.85 | -5.69 | 21.90 |
| SSP5-8.5 | PRINCE CHARLES MOUNTAINS | *Tr*_S02 | 12.38 | -4.47 | 22.60 |
| SSP5-8.5 | PRINCE CHARLES MOUNTAINS | *Tr*_S18 | 1.56 | -4.20 | 8.93 |
| SSP5-8.5 | TRANSANTARCTIC MOUNTAINS | *Carbonea vorticosa* | -10.18 | -18.31 | 5.52 |
| SSP5-8.5 | TRANSANTARCTIC MOUNTAINS | *Lecanora fuscobrunnea* | 23.60 | -61.13 | 66.39 |
| SSP5-8.5 | TRANSANTARCTIC MOUNTAINS | *Lecidea andersonii* | 60.68 | -22.16 | 94.85 |
| SSP5-8.5 | TRANSANTARCTIC MOUNTAINS | *Lecidea atrobrunnea* | 0.00 | 0.00 | 0.00 |

| | | | | | |
|---|---|---|---|---|---|
| SSP5-8.5 | TRANSANTARCTIC MOUNTAINS | *Lecidea cancriformis* | 47.14 | 14.63 | 67.00 |
| SSP5-8.5 | TRANSANTARCTIC MOUNTAINS | *Lecidea polypycnidophora* | 5.15 | -4.90 | 10.07 |
| SSP5-8.5 | TRANSANTARCTIC MOUNTAINS | *Lecidella greenii* | 30.16 | 18.74 | 41.87 |
| SSP5-8.5 | TRANSANTARCTIC MOUNTAINS | *Lecidella siplei* | 20.63 | -22.68 | 47.30 |
| SSP5-8.5 | TRANSANTARCTIC MOUNTAINS | *Rhizoplaca macleanii* | -5.06 | -7.62 | 0.88 |
| SSP5-8.5 | TRANSANTARCTIC MOUNTAINS | *Tr*_A02 | <u>64.78</u> | 3.33 | 91.53 |
| SSP5-8.5 | TRANSANTARCTIC MOUNTAINS | *Tr*_I01 | 42.88 | -45.72 | 88.64 |
| SSP5-8.5 | TRANSANTARCTIC MOUNTAINS | *Tr*_S02 | 51.53 | -31.36 | 79.26 |
| SSP5-8.5 | TRANSANTARCTIC MOUNTAINS | *Tr*_S18 | 11.54 | -72.17 | 47.73 |

**Table S5** Distance from Coast in km for each pixel with occurrence probability > 500 for SSP1-2.6 and SSP5-8.5

| **Number of overlapping species ranges** | **Current** | **SSP1-2.6** | **SSP5-8.5** | **Difference current to SSP1-2.6** | **Difference current to SSP5-8.5** |
|---|---|---|---|---|---|
| 1 | 20.56 | 17.85 | 12.02 | -2.70 | -8.54 |
| 2 | 18.75 | 22.55 | 27.65 | 3.81 | 8.90 |
| 3 | 13.30 | 14.37 | 23.90 | 1.06 | 10.59 |
| 4 | 9.58 | 12.20 | 20.93 | 2.62 | 11.35 |
| 5 | 6.78 | 8.70 | 14.32 | 1.92 | 7.54 |
| 6 | 4.42 | 6.24 | 9.39 | 1.82 | 4.96 |
| 7 | 4.02 | 4.40 | 8.41 | 0.38 | 4.39 |
| 8 | 3.27 | 3.99 | 5.13 | 0.71 | 1.85 |
